# Prion propagation is controlled by a hierarchical network involving the nuclear Tfap2c and hnRNP K factors and the cytosolic mTORC1 complex

**DOI:** 10.1101/2024.10.21.619371

**Authors:** Stefano Sellitto, Davide Caredio, Matteo Bimbati, Giovanni Mariutti, Martina Cerisoli, Lukas Frick, Vangelis Bouris, Carlos Omar Oueslati Morales, Dalila Laura Vena, Sandesh Neupane, Federico Baroni, Kathi Ging, Jiang-An Yin, Elena De Cecco, Andrea Armani, Adriano Aguzzi

**Affiliations:** Institute of Neuropathology, University of Zurich, Schmelzbergstrasse 12, CH-8091 Zurich, Switzerland; Department of Biomedical Sciences, University of Padova, Via Ugo Bassi 58/B - 35131 Padova, Italy

## Abstract

Heterogeneous Nuclear Ribonucleoprotein K (hnRNP K) is a limiting factor for prion propagation. However, little is known about the function of hnRNP K except that it is essential to cell survival. Here, we performed a synthetic-viability CRISPR ablation screen to identify epistatic interactors of *HNRNPK*. We found that deletion of Transcription Factor AP-2γ (*TFAP2C*) suppressed the death of hnRNP K-depleted LN-229 and U-251 MG cells, whereas its overexpression hypersensitized cells to hnRNP K loss. *HNRNPK* ablation decreased cellular ATP, downregulated genes related to lipid and glucose metabolism, and enhanced autophagy. Co-occurrent deletion of *TFAP2C* reversed these effects, restoring transcriptional balance and alleviating energy deficiency. We linked *HNRNPK* and *TFAP2C* interaction to mTOR signaling, observing that *HNRNPK* ablation inhibited mTORC1 activity through downregulation of mTOR and Rptor, while *TFAP2C* overexpression enhanced mTORC1 downstream functions. In prion-infected cells, *TFAP2C* activation reduced prion levels and countered the increased prion propagation caused by *HNRNPK* suppression. Short-term pharmacological inhibition of mTOR also elevated prion levels and partially mimicked the effects of *HNRNPK* silencing. Our study identifies *TFAP2C* as a genetic interactor of *HNRNPK*, implicates their roles in mTOR metabolic regulation, and establishes a causative link between these activities and prion propagation.

## Introduction

hnRNP K is a highly conserved multifunctional protein expressed in nearly all mammalian tissues (*1–3*). hnRNP K has been described as a DNA/RNA binding protein involved in several stages of RNA metabolism through mechanisms that are not fully understood (*4–11*). *HNRNPK* can act as an oncogene or tumor suppressor in numerous malignancies (*1, 12, 13*) and is linked to various neuronal functions (*14–16*). Its mutations and dysregulated expression are implicated in neurodevelopmental and neurodegenerative conditions such as Au-Kline syndrome (*17, 18*), Spinocerebellar Ataxia Type 10 (*19*), Amyotrophic Lateral Sclerosis, and Frontotemporal Lobar Degeneration (*20–25*). We recently reported a role of hnRNP K in limiting the conversion of the cellular prion protein PrP^C^ into transmissible prions (PrP^Sc^) (*26*), a process referred to as prion propagation or replication (*27, 28*).

The involvement of hnRNP K in disparate neurodegenerative proteinopathies suggests a broad role in protein folding and homeostasis. A better understanding of these functions may help elucidate shared mechanisms of genetic and molecular abnormalities among different neurodegenerative disorders. However, the essentiality of *HNRNPK,* whose genetic ablation is lethal to cells (*29–31*), and its tightly regulated expression limit the usefulness of loss/gain-of-function studies to investigate *HNRNPK*’s functions.

Here, we performed a genome-wide synthetic-viability CRISPR screen to discover epistatic interactors that might suppress the lethality of hnRNP K loss-of-function and provide insights into its cellular roles. We found that the ablation of Transcription Factor AP-2γ (*TFAP2C*) mitigated the death of *HNRNPK*-ablated cells, whereas its overexpression further sensitized cells to the loss of hnRNP K. Also, we found that *HNRNPK* deletion reduced the transcription of genes related to fatty acid, sterol, and glucose metabolism, lowered intracellular ATP, and increased autophagic flux through mTORC1 downregulation and AMPK activation; all these perturbations were partially prevented by *TFAP2C* co-deletion. Conversely, *TFAP2C* overexpression enhanced mTORC1 signaling.

We previously found that shifts in energy metabolism accompany *HNRNPK*-modulated prion propagation (*26*). Accordingly, mTOR inhibition increased prion propagation, partially reproducing the effect of *HNRNPK* silencing, while *TFAP2C* overexpression reduced PrP^Sc^ replication and limited its *HNRNPK*-induced accumulation.

These findings suggest that *TFAP2C*, *HNRNPK*, and mTORC1 interact to regulate cell death, metabolic homeostasis, and prion propagation.

## Results

### A cellular model to study HNRNPK essentiality

We aimed to identify genes whose loss alleviates, or exacerbates, the impaired cellular fitness caused by the depletion of hnRNP K. As model systems, we chose the human glioblastoma-derived LN-229 and U-251 MG cell lines which express high levels of *HNRNPK* (*2, 3*). We employed a plasmid harboring quadruple non-overlapping single-guide RNAs (qgRNAs), driven by four distinct constitutive promoters, to target the human *HNRNPK* gene in polyclonal LN-229 and U-251 MG cells stably expressing Cas9 (*32*). Seven days after qgRNA lentiviral delivery, we observed a substantial reduction in hnRNP K protein followed, as expected, by a drop in cell viability (Fig. 1A-D). A minor fraction of LN-229 cells exhibiting low or no Cas9 expression did not undergo *HNRNPK* ablation, resulting in incomplete cell death (Fig. 1A, 1D). To address this issue, we isolated by limiting dilutions LN-229 clones expressing high Cas9 levels (Supp. Fig. 1A). We compared Cas9 activity in seven individual clones using an *eGFP* reporter and selected LN-229 clone C3 (Supp. Fig. 1B). When we tested the *HNRNPK* ablation efficiency in LN-229 C3 cells, we observed complete protein depletion and cell death (Fig. 1A-B). Interestingly, U-251 MG Cas9 cells showed delayed cell death compared to LN-229 C3 (Fig. 1A).

**Figure 1.**
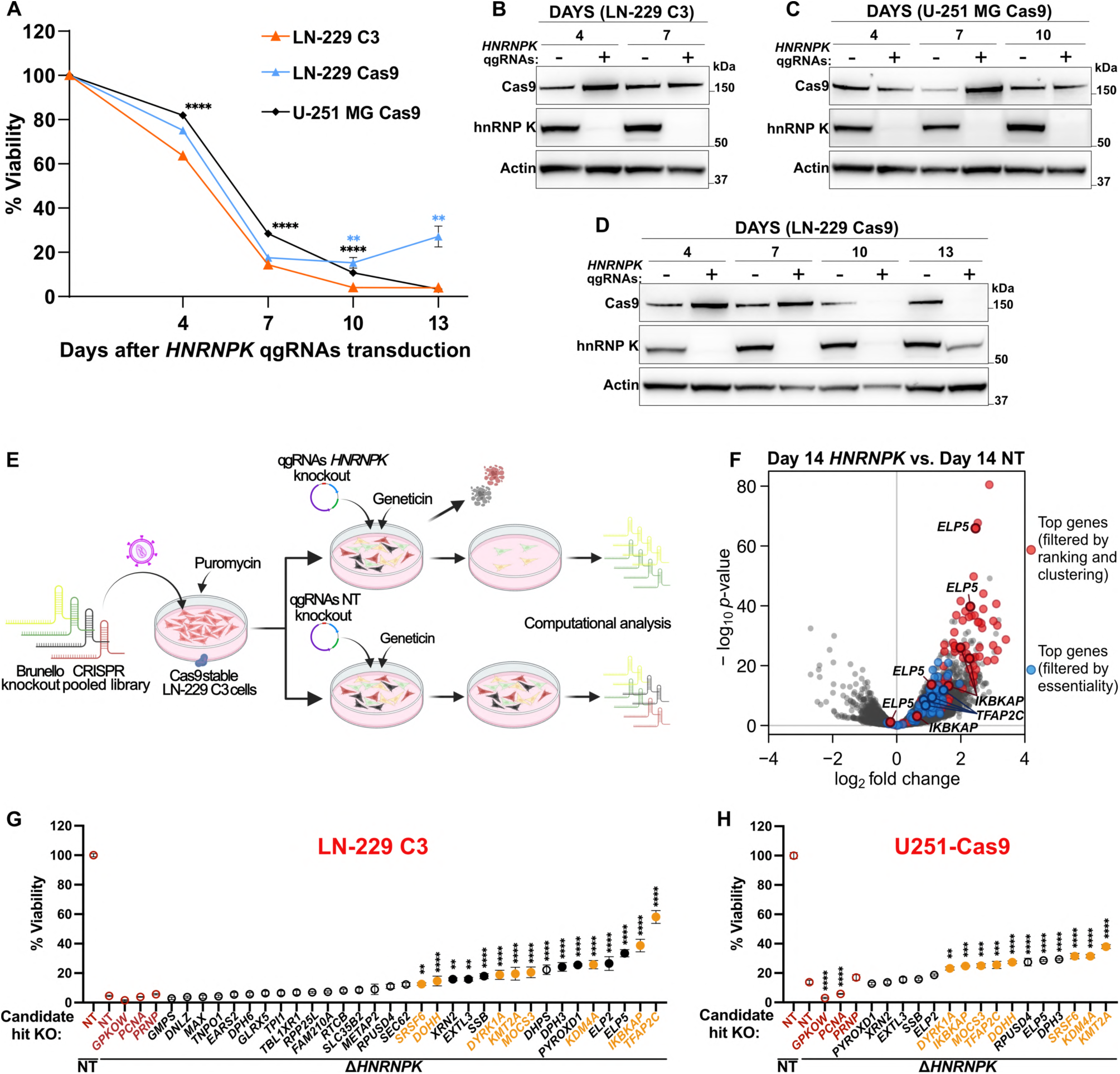
**A.** Cell viability after delivery of *HNRNPK* or non-targeting (NT) qgRNAs (CellTiter-Glo assay). Results are normalized against the NT condition. *n* = 10. **B-D.** Cas9 and hnRNP K protein upon delivery of *HNRNPK* (+) or NT (-) qgRNA. **E.** Genome-wide CRISPR deletion screen. **F.** Volcano plot showing differential sgRNAs abundance in the *HNRNPK* vs. NT comparison at day 14. **G-H.** Cell viability after sequential co-deletion of each candidate gene and *HNRNPK* (CellTiter-Glo assay). Results are normalized against the seeded cell density before *HNRNPK* ablation and compared to the double NT+NT condition. Red empty circles: non-targeting control (NT) and non-specific genes (*PCNA*, *GPKOW*, *PRNP*). Black-filled circles: genes confirmed only in LN-229 C3 cells. Yellow-filled circles: genes confirmed in both LN-229 C3 and U251-Cas9 cells. *n* ≥ 3. **Data information:** *n* represents independent cultures. Mean ± SEM. ^✱✱^: *p* < 0.01; ^✱✱✱^: *p* < 0.001; ^✱✱✱✱^: *p* < 0.0001 (Two-way ANOVA Tukey’s test in A. One-way ANOVA Dunnett’s test in G-H).

To confirm that the observed lethality resulted from the absence of hnRNP K, we transduced LN-229 C3 cells with constructs encoding the *HNRNPK* ORF sequence under transcriptional control of the elongation factor 1α (EF-1α) promoter. We then utilized intron-targeting single-guide RNAs (sgRNAs) to selectively ablate the endogenous *HNRNPK* gene (Supp. Table 1). The cell death resulting from *HNRNPK* deletion was suppressed by the exogenous constructs, confirming the specificity of the lethal phenotype and the reliability of this cellular model (Supp. Fig. 1C-D).

### Genome-wide CRISPR ablation screen for the identification of HNRNPK epistatic interactors

To identify functionally relevant epistatic interactors of *HNRNPK*, we conducted a whole-genome ablation screen in LN-229 C3 cells using the Human CRISPR Brunello pooled library (*33*), which targets 19,114 genes with an average of four distinct sgRNAs per gene (total = 76,441 sgRNAs). The lentiviral transduction of the Brunello library was followed by six days of antibiotic selection and subsequent lentiviral delivery of qgRNA vectors containing either *HNRNPK*-specific or non-targeting (NT) qgRNA guides. Cells underwent antibiotic selection for six more days before harvesting and gDNA extraction (Fig. 1E, Supp. Fig. 2A). sgRNAs distribution was analyzed by next-generation Illumina sequencing (NGS) at the onset of the screen after the library transduction (Day 1) and at the screen endpoint (Day 14; Supp. Fig. 2A).

Two independent screens were conducted on different days and yielded a robust correlation indicative of satisfactory technical performance (Supp. Fig. 2B). When using the DepMap repository (*30, 31*) to compare the representation of essential genes in LN-229 cells at Day 14 NT vs. Day 1 (Supp. Table 2), we found that 75% of the sgRNAs targeting known essential genes were efficiently depleted (log_2_ fold change ≤ −1, FDR ≥ 0.01) with >92% of those essential genes having ≥2 sgRNAs dropped below threshold (Supp. Fig 2C-E).

We then listed genes whose sgRNAs were over- or underrepresented in the *HNRNPK* vs. NT at day 14, reasoning that their deletion would modify the lethality resulting from hnRNP K removal. We obtained a list of 763 and 37 significantly enriched and depleted genes, respectively (log_2_ fold change ≥1 or ≤ −1, FDR ≥0.01; Supp. Fig. 2F, Supp. Table 3). Pathway analysis of genes enriched with ≥2 sgRNAs yielded gene ontology (GO) terms related to ribosomal biogenesis, tRNA processing, non-coding RNA metabolism, and translation, consistent with the known roles of hnRNP K in RNA metabolism (Supp. Fig. 2G). Accordingly, ablation of *HNRNPK* in LN-229 C3 cells showed a progressive reduction in global protein synthesis (Supp. Fig. 2H). Also, the GO analysis highlighted “tRNA wobble base modification” as the most overrepresented GO term (Supp. Fig. 2G). Genes encoding for Elongator complex proteins (ELPs), which are included in this pathway, were significantly enriched in the screen (Fig. 1F, Supp. Table 3), suggesting that their deletion counteracts the deleterious effects of *HNRNPK* ablation. Previous CRISPR screens showed that the absence of ELPs prevents apoptosis in metastatic gallbladder cancer (GBC) (*34*). Accordingly, our screen also showed enrichment of other general proapoptotic factors, including *AIFM1*, *MFN2,* and *FADD* (Supp. Table 3).

Among the most profoundly depleted genes were *PCBP1*, *PCBP2,* and *HNRNPA1*, all of which belong to the same genetic superfamily as *HNRNPK* (Supp. Table 3) (*35, 36*). The synthetic lethality deriving from their deletion suggests that these genes cooperate with hnRNP K in cell-essential processes. Conversely, the screen was enriched for sgRNAs targeting *CPSF6*, *NUDT21*, and *XRN2*, which form protein complexes with hnRNP K and regulate RNA maturation processes (Supp. Table 3) (*7, 37*). Hence, our screen identified *HNRNPK* functional partners sensitively and specifically despite the detection of additional, less specific cell death modulators.

### TFAP2C ablation suppresses HNRNPK loss-of-function

To prioritize biologically relevant hits among the 763 enriched genes, we focused on those showing enrichment of all four sgRNAs (Supp. Table 3). We applied the STRING database (*38*) to assess protein-protein interactions and biological pathways associated with these genes. Next, we examined whether any other genes enriched in the screen scored as interactors. This allowed us to identify and select hierarchical functional clusters among our hits (Supp. Table 4). In parallel, we ranked the 763 enriched genes by multiplying their False Discovery Rate (-log_10_ FDR) with their effect size (log_2_ fold change; Supp. Table 4). We focused on genes with ≥2 sgRNA scoring in the top 100 rankings, with ≥1 sgRNA among the top 35. The intersection of this ordered list with the STRING clusters yielded 19 genes (Fig. 1F, Supp. Table 4). We generated a second list including those genes that, independently from the ranking and clustering, had ≥2 sgRNAs enriched and scored as non-essential in the Day 14 NT vs. Day 1 comparison (maximally one sgRNA with log_2_ fold change ≤ −1, FDR ≥ 0.01; Fig. 1F, Supp. Table 4).

Based on these two lists, 32 genes were selected for validation in LN-229 C3 cells. We ablated each gene individually and then deleted *HNRNPK*. Two different non-targeting (NT) qgRNAs were used as controls for two sequential ablations.

We excluded genes whose deletion from *HNRNPK^+/+^*cells resulted in >50% enhanced or impaired cell viability (Supp. Fig. 2I-J). 16 of the initial 32 hits increased viability of Δ*HNRNPK* cells by >2-fold (*p* < 0.01) (Fig. 1G). We then tested these 16 genes also in U-251 MG Cas9 cells (henceforth abbreviated as U251-Cas9 cells) at a log_2_ fold threshold of ≥ 0.5. We confirmed a total of 9 hits (Fig. 1H), including the ELPs gene *IKBAKP* and the transcription factor *TFAP2C*, the two strongest hits identified in LN-229 C3 cells.

*TFAP2C* (Transcription Factor AP-2γ) was particularly interesting because it regulates the expression of several long non-coding RNAs (lncRNAs) (*39–41*) and has critical roles in neurodevelopmental processes (*42, 43*) similar to *HNRNPK* (*9, 17, 18, 37, 44, 45*). Moreover, both *HNRNPK* and *TFAP2C* have been described to modulate glucose metabolism (*46–48*). Therefore, we elected to explore the epistatic interaction between these two genes.

To consolidate the observations above, we repeated the experiments described in Fig. 2G-H by individually deleting only *TFAP2C* in 20 distinct technical replicas (Supp. Fig. 3A-B). As an orthogonal means of confirmation, we assessed the clonogenic potential of the respective ablated cells (Fig. 2A-B). Again, the deletion of *TFAP2C* suppressed the cell death induced by removing hnRNP K in both LN-229 C3 and U251-Cas9 cells, whereas *TFAP2C* ablation alone only slightly reduced their growth rate. Thus, the loss of *TFAP2C* did not induce any intrinsic pro-survival effect, pointing to a specific epistatic interaction between *TFAP2C* and *HNRNPK*.

**Figure 2.**
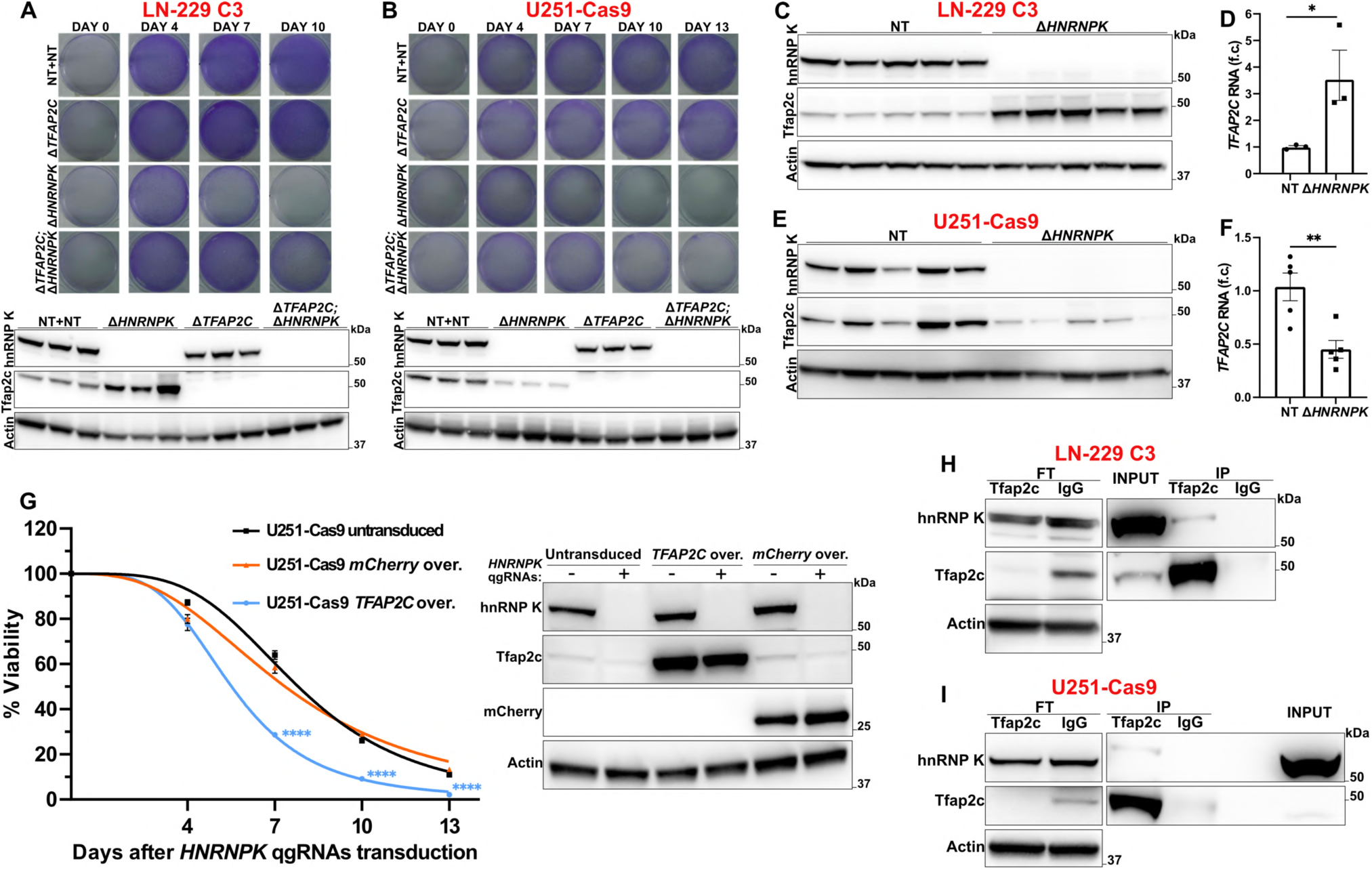
**A-B.** Clonogenic fitness of Δ*TFAP2C* or NT-unmodified cells after delivery of *HNRNPK* or NT qgRNAs (Crystal violet staining). Western blot: hnRNP K and Tfap2c protein 7 (A) and 10 (B) days after qgRNAs delivery. *n* = 3. **C, E.** Tfap2c protein after *HNRNPK* ablation. *n* = 5. **D, F.** qRT-PCR showing *TFAP2C* RNA in LN-229 C3 (D) and U251-Cas9 (F) cells upon *HNRNPK* deletion. *n* = 3 and 5. **G.** Viability course of U251-Cas9 untransduced or overexpressing *TFAP2C* or mCherry after delivery of *HNRNPK* (+) or NT (-) qgRNAs (CellTiter-Glo assay). Results are normalized against the NT condition. *n* = 10. Western blot: hnRNP K, Tfap2c, and mCherry protein 10 days after qgRNAs delivery. **H-I.** Co-immunoprecipitation of Tfap2c and hnRNP K. IP: Immunoprecipitated Protein; FT: Flow-through after immunoprecipitation. **Data information:** qRT-PCR results are normalized against *GAPDH* expression. *n* represents independent cultures. f.c.: fold change. Mean ± SEM. ^✱^: *p* < 0.05, ^✱✱^: *p* < 0.01, ^✱✱✱✱^: *p* < 0.0001 (Unpaired t-test in D and F. Two-way ANOVA Šídák’s test in G).

### TFAP2C upregulation sensitizes cells to the loss of HNRNPK

Following *HNRNPK* ablation, we observed an increase in *TFAP2C* RNA and protein amount in LN-229 C3 cells (Fig. 2C-D). This suggested that the toxicity caused by hnRNP K deletion may be due to *TFAP2C* upregulation. However, *TFAP2C* overexpression using quadruple-guide CRISPR activation (*32*) in *HNRNPK^+/+^* LN-229-dCas9-VPR cells did not impair viability (Supp. Fig. 3C). In U251-Cas9 cells, *HNRNPK* ablation had the opposite effect and decreased both the RNA and protein levels of *TFAP2C* (Fig. 2E-F), potentially explaining their relative resilience to *HNRNPK* ablation (Fig. 1A) and the smaller protective effect mediated by *TFAP2C* deletion in this cell line (Fig. 2A-B, Supp. Fig. 3A-B).

To test if *TFAP2C* overexpression sensitizes cells to the loss of *HNRNPK*, we produced stable U251-Cas9 lines overexpressing *TFAP2C* or mCherry for control and measured their viability after hnRNP K removal. U251-Cas9 cells overexpressing *TFAP2C* experienced a significant acceleration of cell death (*p* < 0.0001; Fig. 2G), confirming the genetic relationship between *TFAP2C* and *HNRNPK* and highlighting a causative dependency between their expression levels and cell death.

Although we efficiently ablated *HNRNPK* in LN-229 and U-251 MG cells, we were unable to overexpress it, possibly due to tight autoregulative feedback loops. However, while Tfap2c levels changed upon *HNRNPK* deletion, neither the ablation nor the overexpression of *TFAP2C* modified the levels of hnRNP K in LN-229 and U-251 MG cells (Supp. Fig. 3D-I), disproving the existence of transcriptional feedback loop between *HNRNPK* and *TFAP2C*.

### Nuclear colocalization and interaction between hnRNP K and Tfap2c

We then asked whether hnRNP K and Tfap2c proteins physically interact and modulate their reciprocal subcellular localization. When staining hnRNP K and Tfap2c by immunofluorescence in LN-229 C3 and U251-Cas9 cells, we noticed a nuclear overlap between the two proteins but no change in their subcellular distribution after ablation of *TFAP2C* or *HNRNPK* (Supp. Fig. 3J-K). We also observed specific co-immunoprecipitation of hnRNP K and Tfap2c in LN-229 C3 and U251-Cas9 cells (Fig. 2H-I, Supp. Fig. 3L), suggesting that the two proteins form a complex inside the nucleus.

### HNRNPK and TFAP2C control the activation of caspase-dependent apoptosis but not ferroptosis

We asked if the deletion of *HNRNPK* causes apoptosis. We detected increased levels of PARP and Caspase 3 cleavage in LN-229 C3 cells upon ablation of *HNRNPK* (Fig. 3A). Conversely, the prior removal of *TFAP2C* limited the cleavage of these two proteins (Fig. 3A), suggesting that *TFAP2C* ablation prevents *HNRNPK*-induced apoptosis. PARP cleavage had the same pattern in U251-Cas9 cells; however, these cells did not show cleavage of Caspase 3 upon deletion of *HNRNPK*, *TFAP2C,* or both (Fig. 3B). We then asked whether the pan-caspase inhibitor Z-VAD-FMK reduced the lethality resulting from *HNRNPK* deletion. Z-VAD-FMK decreased cell death consistently and significantly in LN-229 C3 and U251-Cas9 cells transduced with *HNRNPK* ablation qgRNAs (Fig. 3C-D). These data confirm that *HNRNPK* deletion promotes cell apoptosis. We then treated LN-229 C3^Δ*TFAP2C*^ and U251-Cas9^Δ*TFAP2C*^ cells with increasing concentrations of staurosporine, a potent apoptosis inducer. Both cell lines were only mildly resistant to the apoptotic action of staurosporine (Fig. 3E-F), indicating that *TFAP2C* ablation exerts its protective effect on *HNRNPK*-deleted cells by modulating upstream processes which eventually converge on apoptosis.

**Figure 3.**
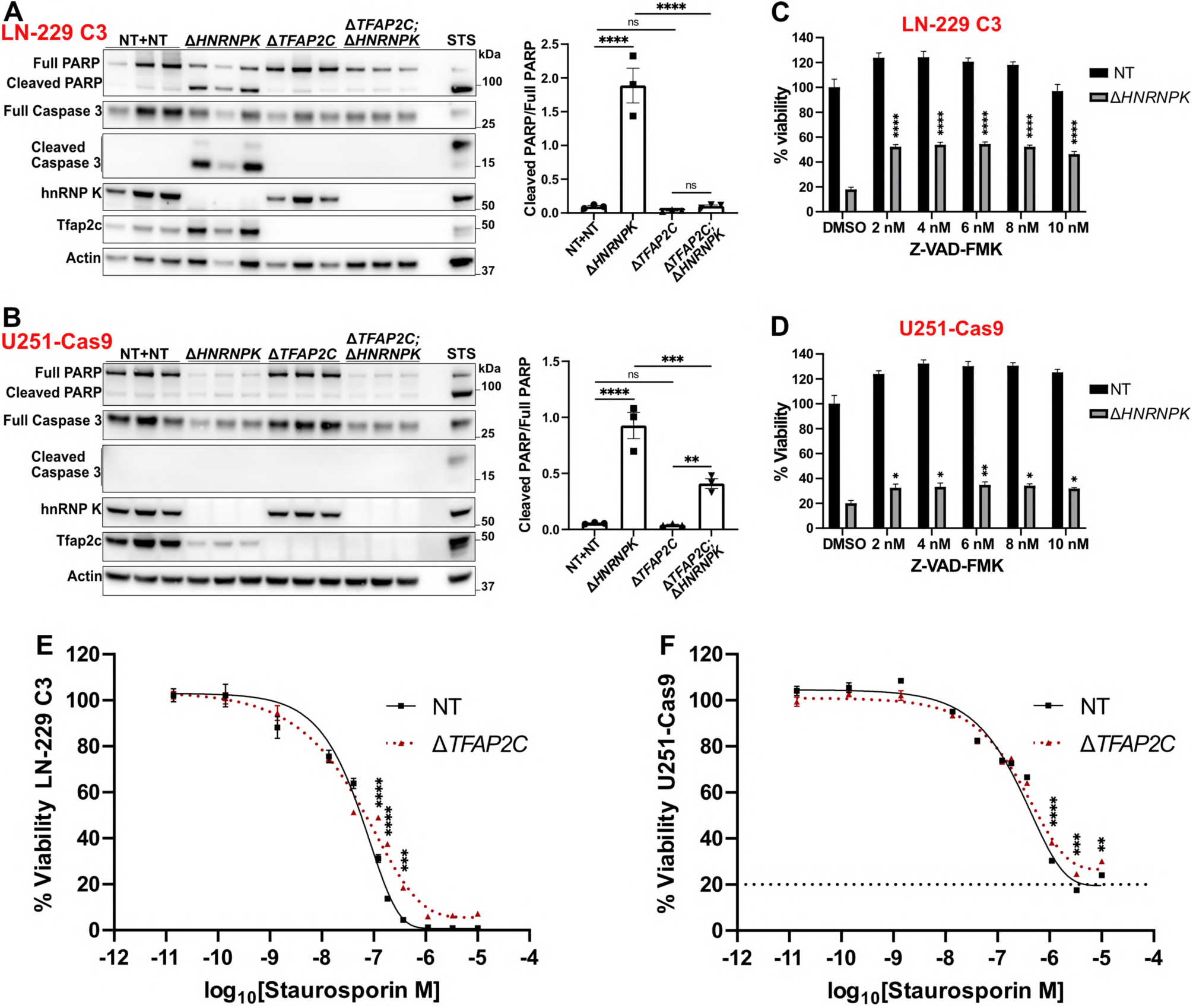
**A-B.** PARP and Caspase 3 protein cleavage ratio after *HNRNPK* and *TFAP2C* deletion. 4 hours 1 μM staurosoprine (STS) was used as positive control. *n* = 3. **C-D.** Viability of cells 7 (C) or 10 (D) days after *HNRNPK* or NT qgRNAs transduction and supplementation of Z-VAD-FMK (CellTiter-Glo assay). Results are normalized against the DMSO/NT condition *n* = 6. **E-F.** Viability of Δ*TFAP2C* cells treated with staurosporine or DMSO (CellTiter-Glo assay). Results are normalized against the DMSO-treated cells. *n* = 4. **Data information:** *n* represents independent cultures. Mean ± SEM. ^✱^: *p* < 0.05, ^✱✱^: *p* < 0.01, ^✱✱✱^: *p* < 0.001, ^✱✱✱✱^: *p* < 0.0001 (Two-way ANOVA Uncorrected Fisher’s LSD in A-B, Dunnett’s test in C-D, and Šídák’s test in E-F).

To investigate whether *HNRNPK* and *TFAP2C* ablation modulate additional cell death pathways, we explored their potential involvement in ferroptosis, a form of cell death marked by lipid peroxides accumulation. *TFAP2C* regulates ferroptosis by enhancing the expression of the Glutathione Peroxidase 4 (*GPX4*) upon selenium detection (*41, 49*). We inquired whether ferroptosis was activated upon ablation of *HNRNPK*. In LN-229 C3 and U251-Cas9 cells, the deletion of *HNRNPK* reduced the protein level of GPX4, whereas *TFAP2C* deletion increased it (Supp. Fig. 4A-B). Moreover, the ablation of *HNRNPK* led to higher lipid peroxidation in LN-229 C3 cells, which was increased in the absence of *TFAP2C* (Supp. Fig. 4C). To investigate this further, we challenged LN-229 C3^Δ*TFAP2C*^ and U251-Cas9^Δ*TFAP2C*^ cells with increasing doses of erastin, a commonly used ferroptosis inducer. *TFAP2C* ablation did not prevent erastin toxicity (Supp. Fig. 4D-E). Additionally, different anti-ferroptosis drugs did not suppress the lethality of hnRNP K-depleted LN-229 C3 cells (Supp. Fig. 4F). These results suggest a role for *HNRNPK* and *TFAP2C* in balancing the protein levels of GPX4. However, ferroptosis seems only marginally connected to the essentiality of *HNRNPK* and is unlikely to be the primary toxic pathway activated by its removal.

### HNRNPK deletion perturbs the transcriptional regulation of genes involved in lipid and glucose metabolism

Apoptosis is often activated downstream of various stress signaling pathways. To gain a mechanistic understanding of the early events shaping the genetic interaction between *TFAP2C* and *HNRNPK*, we sequenced total RNA in LN-229 C3 cells depleted of hnRNP K, Tfap2c, or both (Supp. Table 5). We lysed cells four days after delivery of *HNRNPK* qgRNAs. At this time point, ablation was already extensive (Fig. 1B, Supp. Fig. 5A), yet cell viability was mostly preserved (Fig. 1A, 2A). Gene Set Enrichment Analysis (GSEA) showed significant downregulation of genes involved in sterol biosynthesis, secondary alcohol metabolism, and fatty acids catabolism (FDR < 0.05) after removal of hnRNP K (Δ*HNRNPK* vs. NT) (Fig. 4A). Conversely, the most upregulated genes in LN-229 C3^Δ*TFAP2C;*Δ*HNRNPK*^ versus LN-229 C3^Δ*HNRNPK*^ cells pertained to lipid metabolism and lysosomal functions including sterol and secondary alcohol metabolic processes (Fig. 4B). The intersection of these upregulated genes (log_2_ fold change ≥ 0.5) with genes downregulated upon depletion of hnRNP K (log_2_ fold change ≤ 0.5) yielded a significant enrichment of GO terms related to energy production and catabolic functions, including carbohydrate metabolism (Fig. 4C-D, Supp. Table 5). Factors involved in lipid metabolism and cholesterol biosynthesis like *FASN*, *SREBF1*, *LSS,* and *DHCR7*, and genes related to glycolysis and the pentose phosphate pathway, including *PGLS*, *G6PD*, *TPI1, H6PD,* and *GPI,* underwent consistent bidirectional regulation (Fig. 4C, Supp. Table 5). These data suggest that the transcriptional regulation of lipid and glucose metabolism is imbalanced by the removal of hnRNP K and is partially restored by *TFAP2C* deletion.

**Figure 4.**
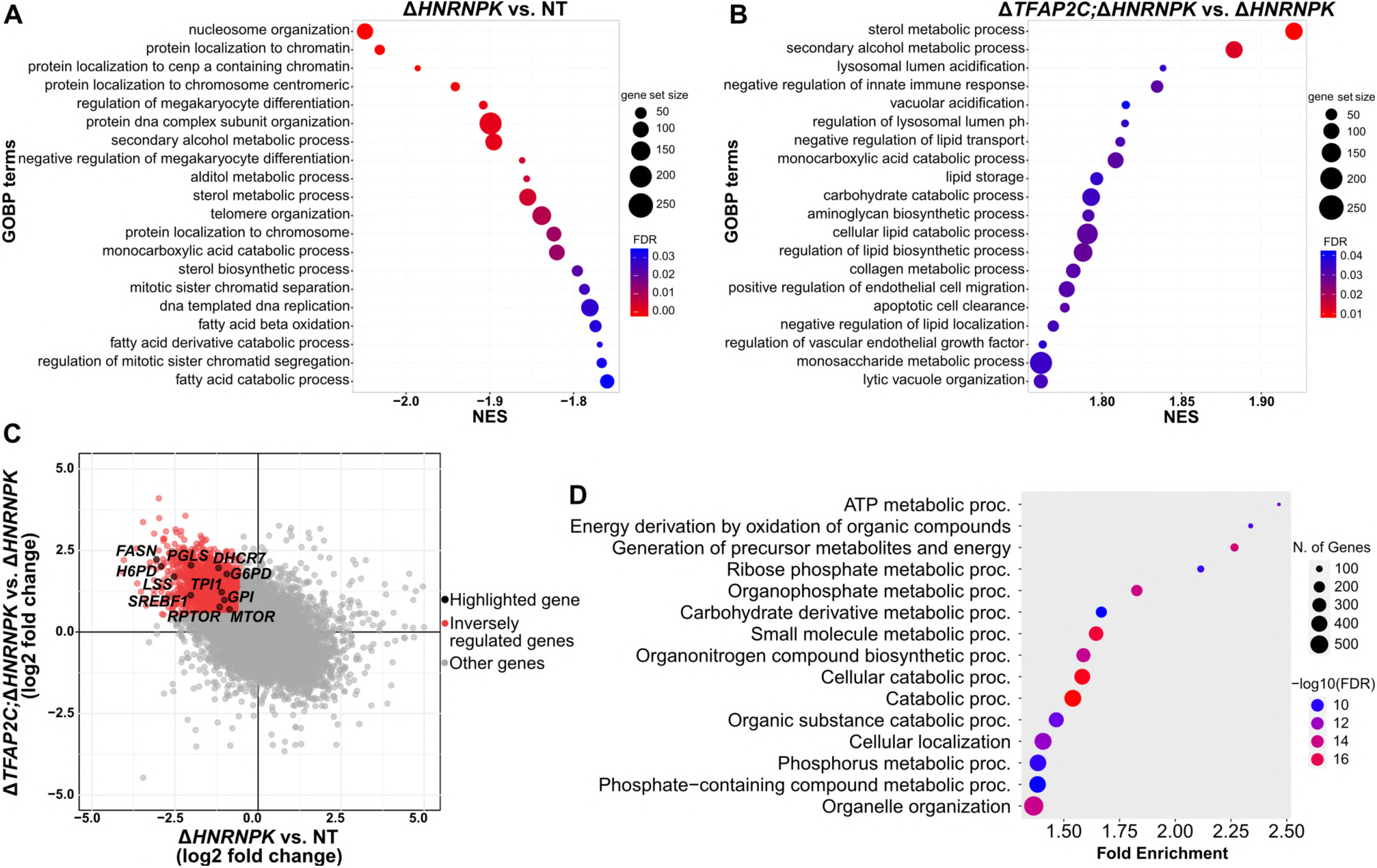
**A-B.** RNA-seq Gene Set Enrichment Analysis. Shown are the 20 most negatively (A) and positively (B) enriched GOBP terms (Gene Ontology Biological Process), respectively, in Δ*HNRNPK* vs. NT (A) and Δ*TFAP2C;*Δ*HNRNPK* vs. Δ*HNRNPK* (B). NES: Normalized Enrichment Score. **C.** Intersection of RNA-seq data resulting from the Δ*TFAP2C;*Δ*HNRNPK* vs. Δ*HNRNPK* and the Δ*HNRNPK* vs. NT comparisons. **D.** Gene enrichment of the biological process for the inversely regulated genes highlighted in C.

### HNRNPK and TFAP2C regulate cell metabolism and bioenergetics via mTOR and AMPK

The deletion of *TFAP2C* and *HNRNPK* caused a broad transcriptional rewiring of cell bioenergetics and metabolism (Fig. 4A-D, Supp. Table 5). Accordingly, LN-229 C3 cells had reduced intracellular ATP four days after delivery of *HNRNPK* ablation qgRNAs. Conversely, ATP levels slightly increased after *TFAP2C* deletion and remained high when both genes were ablated (Fig. 5A). AMP-activated protein kinase (AMPK) is a well-established ATP/AMP sensor, and it is phosphorylated and activated in the context of low cellular energy (*50*). Accordingly, we observed increased AMPK phosphorylation (pAMPK) upon ablation of *HNRNPK*, which was consistently reduced in LN-229 C3^Δ*TFAP2C*^ cells (Supp. Fig. 5B). LN-229 C3^Δ*TFAP2C;*Δ*HNRNPK*^ cells also showed a partial reduction of pAMPK relative to LN-229 C3^Δ*HNRNPK*^ cells (Supp. Fig. 5B). These results suggest that hnRNP K depletion causes an energy shortfall, leading to cell death.

**Figure 5.**
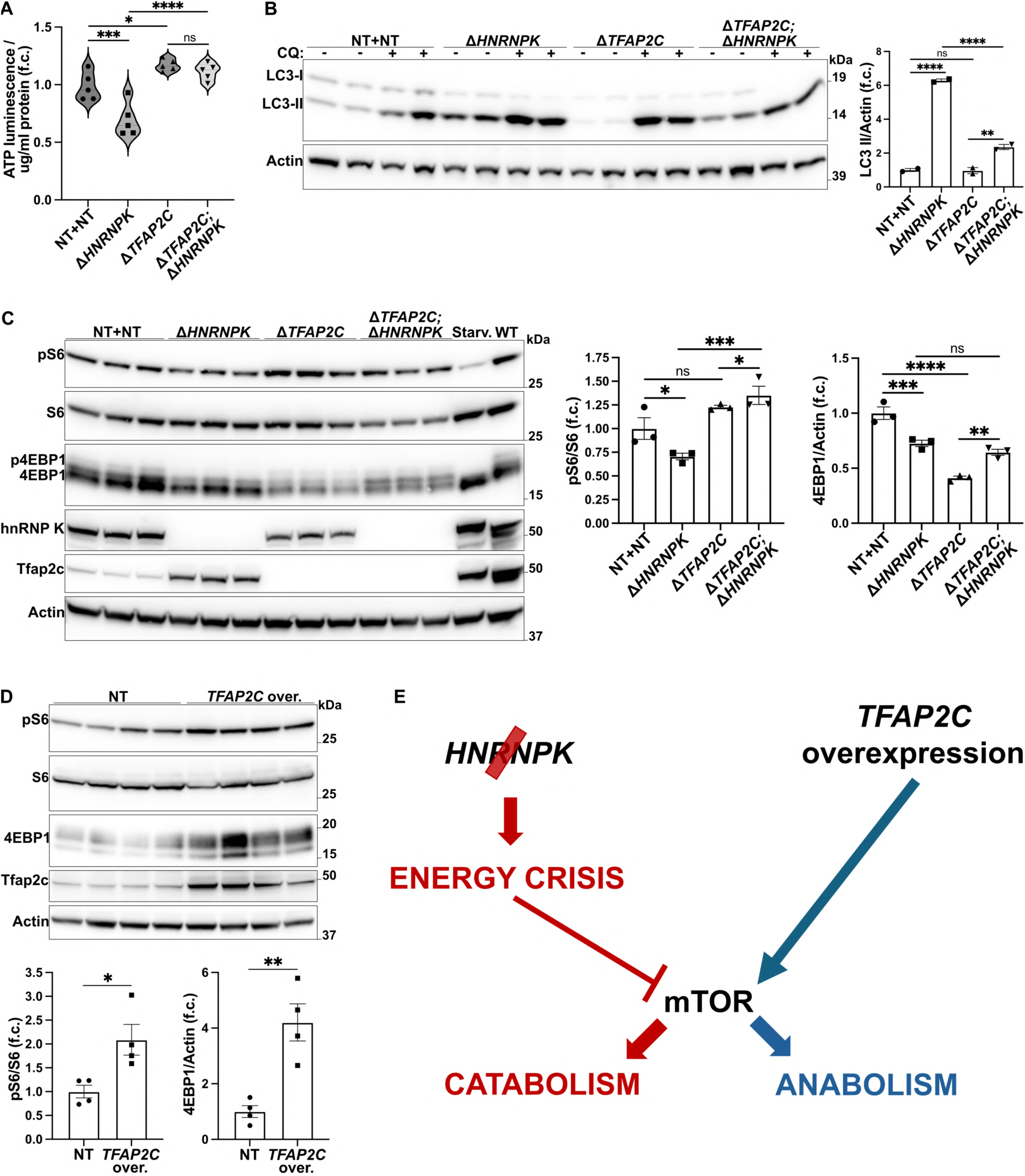
**A.** Intracellular ATP level in Δ*TFAP2C* or NT-unmodified LN-229 C3 cells 4 days upon delivering *HNRNPK* and NT qgRNAs. *n* = 5. **B.** LC3-II protein in LN-229 C3 cells after *HNRNPK* and *TFAP2C* ablation. 4 hours 100 μM chloroquine (CQ) was used to monitor the autophagic flux. Basal autophagy (without chloroquine) is quantified. *n* = 2. Deletions are shown in Supp. Fig. 5C. **C.** 4EBP1 and S6 protein phosphorylation after *HNRNPK* and *TFAP2C* ablation in LN-229 C3 cells. 6h HBSS-starvation (starv.) was used as positive control. *n* = 3. **D.** 4EBP1 and S6 protein phosphorylation in LN-229 dCas9-VPR cells upon *TFAP2C* overexpression. *n* = 4. **E.** Hypothesized mechanism of *TFAP2C* and *HNRNPK* in cell metabolism regulation. **Data information:** *n* represents independent cultures. f.c.: fold change. Mean ± SEM. ns: *p* > 0.05, ^✱^: *p* < 0.05, ^✱✱^: *p* < 0.01, ^✱✱✱^: *p* < 0.001, ^✱✱✱✱^: *p* < 0.0001 (Two-way ANOVA Uncorrected Fisher’s LSD in A-C. Unpaired t-test in D).

Autophagy is a central process modulated by AMPK downstream activity. Consistently, and as previously reported (*51*), the LC3-II autophagy marker increased significantly (*p* < 0.0001) in LN-229 C3^Δ*HNRNPK*^ cells, suggestive of an energy crisis triggering autophagy, but less in LN-229 C3^Δ*TFAP2C;*Δ*HNRNPK*^ cells (Fig. 5B, Supp. Fig. 5C). Together with AMPK, the mechanistic target of rapamycin (mTOR) is a key regulator of cell anabolic and catabolic processes that senses and integrates upstream inputs related to cellular oxygen, nutrients, and energy levels (*52*). mTOR and AMPK control autophagy by regulating the phosphorylation of several downstream targets (*50, 52, 53*). Under nutrient sufficiency, mTOR phosphorylates Ulk1 at Ser758 (pUlk1), preventing its interaction with AMPK and inhibiting autophagosome formation (*53*). To test if mTOR directly modulated the observed autophagy alterations, we measured the phosphorylation of Ulk1. As anticipated, Ulk1 phosphorylation was reduced by *HNRNPK* ablation, homeostatically restored by concomitant Δ*TFAP2C;*Δ*HNRNPK* double ablation, and increased by single *TFAP2C* removal (Supp. Fig. 5D).

These results suggest that mTOR activity is central to the functional interaction between *HNRNPK* and *TFAP2C*. Interestingly, RNA and protein levels of mTOR were downregulated in LN-229 C3^Δ*HNRNPK*^ cells but were partially rebalanced by the Δ*TFAP2C;*Δ*HNRNPK* double deletion (Fig. 4C, Supp. Fig. E). RNA levels of the mTOR associated protein Rptor, which assembles with mTOR in the mTORC1 complex, followed the same trend (Fig. 4C). 4EBP1 and S6 are two downstream targets of mTORC1 kinase activity. Deletion of *HNRNPK* diminished the highly phosphorylated forms of 4EBP1, which instead were preserved in both LN-229 C3^Δ*TFAP2C*^ and LN-229 C3^Δ*TFAP2C;*Δ*HNRNPK*^ cells (Fig. 5C). Similarly, the S6 phosphorylation ratio was reduced in LN-229 C3^Δ*HNRNPK*^ cells and was restored in the Δ*TFAP2C;*Δ*HNRNPK* double-ablated cells (Fig. 5C). Total expression of 4EBP1 was reduced in both *HNRNPK* and *TFAP2C* single ablation and only partially rescued upon double deletion (Fig. 5C).

We conclude that *HNRNPK* and *TFAP2C* play an essential role in co-regulating cell metabolism homeostasis by influencing mTOR and AMPK activity and expression.

### TFAP2C promotes mTORC1 activity

The above results suggest that *HNRNPK* ablation causes an energy crisis parallel to, or followed by, the inhibition of mTORC1 activity and a shift toward catabolism. *TFAP2C* deletion did not induce energetic impairment, yet it affected the mTORC1 pathway by decreasing the expression of 4EBP1. To clarify these observations, we overexpressed *TFAP2C* in LN-229-dCas9-VPR cells. We found that both the phosphorylation ratio of S6 and expression levels of 4EBP1 increased upon *TFAP2C* upregulation (Fig. 5D), although we did not observe changes in mTOR protein expression (Supp. Fig. 5F). These data specify a role for *TFAP2C* in promoting mTORC1-mediated cell anabolism and suggest that its overexpression might hypersensitize cell to *HNRNPK* ablation by depleting the already limited ATP available, thus making its deletion advantageous (Fig. 5E).

### TFAP2C overexpression reduces prion levels and prevents HNRNPK-induced prion propagation

Transcriptional silencing of hnRNP K causes increased generation of infectious prions (PrP^Sc^) in various cell lines (*26*), whereas psammaplysene A (PSA), a compound that interacts with hnRNP K (*54*), reduces prion levels. *HNRNPK* knockdown and PSA treatment induce an inversely symmetric transcriptional profile with downregulation and upregulation of genes related to glycolysis and energy metabolism (*26*). These results suggest that cell stress and energy homeostasis may impact prion propagation and protein aggregation (*55–63*). As we observed an inverse relationship between *HNRNPK* ablation and *TFAP2C* activation in mTORC1 metabolic signaling (Fig. 5A, 5D-E, Supp. Fig. 5B, 5F), we wondered whether *TFAP2C* overexpression might influence prion propagation in a manner opposite to *HNRNPK* downregulation.

Because human prions are highly biohazardous and replicate poorly in most human cells, we used CRISPR/Cas9 to produce an LN-229^Δ*PRNP*^ subline devoid of human PrP^C^ (Supp. Fig. 6A-B). We then expressed a vector plasmid encoding the VRQ allele of the ovine prion protein (*ovPRNP*) to generate an isogenic ovinized line termed “HovL” (for “human ovinized LN-229”; Supp. Fig. 6A-B). As reported for other ovinized cell models (*64*), HovL cells were susceptible to infection by the PG127 strain of ovine prions and capable of sustaining chronic prion propagation (Supp. Fig. 6C-E).

We then acutely downregulated *HNRNPK* and overexpressed *TFAP2C* in PG127-infected HovL cells: 96 hours after inducing *TFAP2C* overexpression (or mCherry for control), we delivered *HNRNPK* siRNA (or non-targeting NT siRNA for control) and incubated HovL cells for the next 96 hours. As expected, proteinase K (PK) digested western blots showed a significant increase of PrP^Sc^ level upon *HNRNPK* downregulation (Fig. 6A-B, Supp. Fig. 7A), with cell viability only minimally affected (Supp. Fig. 7B). Conversely, overexpression of *TFAP2C* consistently reduced PrP^Sc^ level and prevented its accumulation following *HNRNPK* suppression (Fig. 6A-B, Supp. Fig. 7A). Importantly, neither *HNRNPK* silencing nor *TFAP2C* activation perturbed *ovPRNP* RNA or protein level, respectively, in PG127-infected or uninfected HovL (Fig. 6C, Supp. Fig. 7C), indicating that they modulate a post-translational step of the prion life cycle. To assess the kinetics of such effects, we measured PrP^Sc^ level using the antibody 6D11 to PrP, which selectively stains prion-infected cells after fixation and treatment with guanidinium (*65*). Single-cell detection was performed by flow cytometry analysis after overexpression of *TFAP2C* for 2 or 7 days, followed by *HNRNPK* downregulation for 72 or 96 hours, respectively. At 2 days + 72 hours, changes in the % 6D11^+^ cells or total 6D11^+^ fluorescence (% 6D11^+^ cells multiplied for the 6D11^+^ mean intensity) were mild or absent (Fig. 6D), whereas flow cytometry at 7 days + 96 hours showed prominent shifts in both % 6D11^+^ cells and total 6D11^+^ fluorescence (Fig. 6D), coherent with the results observed by PK western blots.

**Figure 6.**
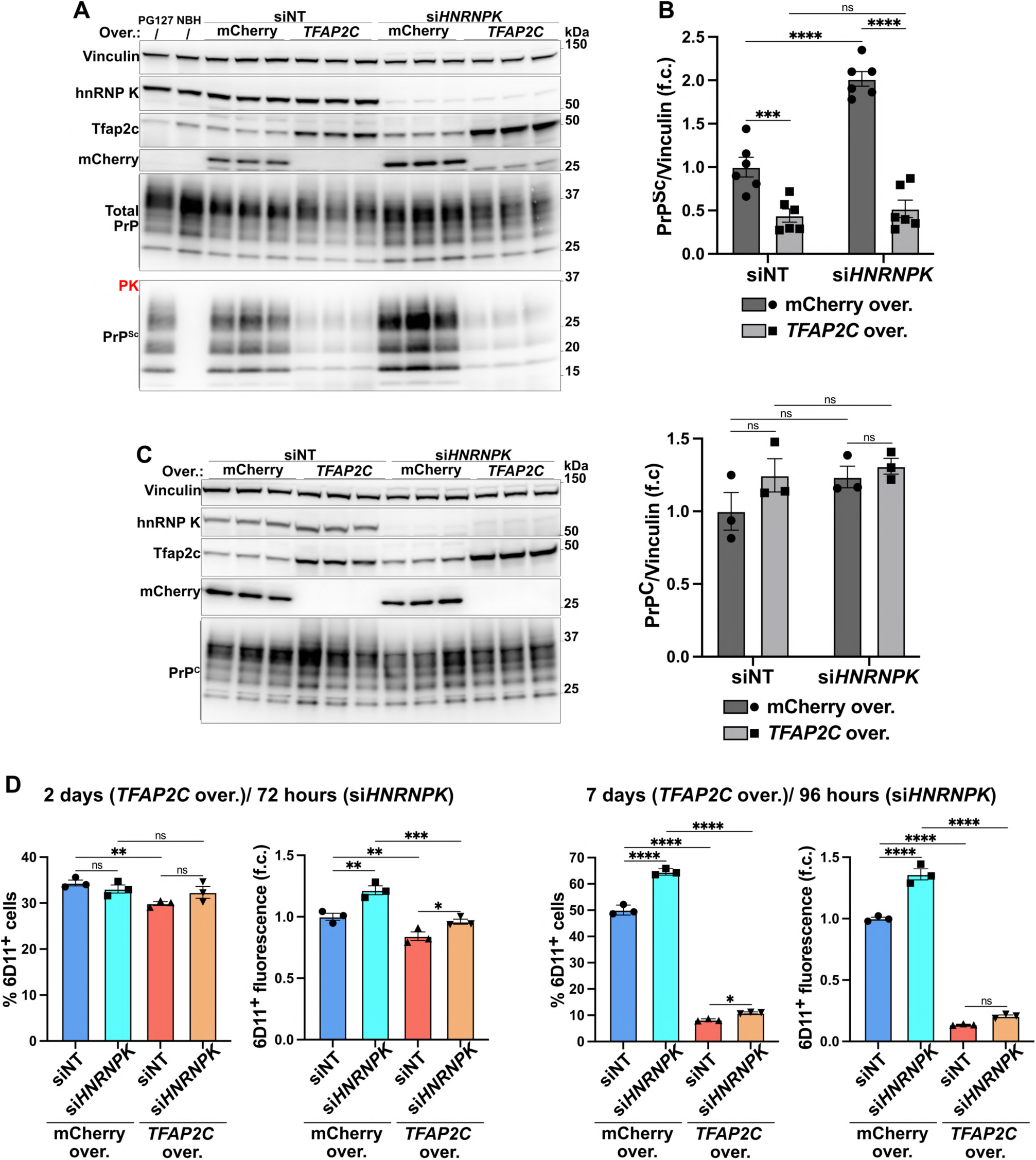
**A.** Proteinase K (PK) digested (bottom) and undigested (top) western blots showing, respectively, PrP^Sc^ and total PrP after *HNRNPK* silencing (96 hours) and *TFAP2C* overexpression (192 hours) in PG127-infected HovL cells. *n* = 3. **B.** Quantification of PrP^Sc^ from B and Supp. Fig. 7A. *n* = 6. **C.** PrP^C^ protein in uninfected HovL cells upon *HNRNPK* knockdown (96 hours) and *TFP2C* overexpression (192 hours). *n* = 3. **D.** Flow cytometry quantification of anti-PrP^Sc^ 6D11 antibody signal in PG127-infected HovL cells upon *HNRNPK* knockdown and *TFP2C* overexpression at different time points. *n* = 3. **Data information:** PG127-infected (PG127) and Not-infectious Brain Homogenate (NBH) were used for PK digestion positive and negative control, respectively. Non-targeting siRNA (siNT) and mCherry overexpression were used as controls. *n* represents independent cultures. f.c.: fold change. Mean ± SEM. ns: *p* > 0.05, ^✱^: *p* < 0.05, ^✱✱^: *p* < 0.01, ^✱✱✱^: *p* < 0.001, ^✱✱✱✱^: *p* < 0.0001 (Two-way ANOVA Uncorrected Fisher’s LSD in B-D).

These data show that *TFAP2C* overexpression and *HNRNPK* downregulation bidirectionally regulate prion propagation. Their combined effect was not additive, indicating a potential convergence on shared molecular pathways ultimately influencing prion propagation.

### Drug inhibition of mTOR partially mimics HNRNPK’s role in driving prion propagation

We showed above that mTORC1 might be a putative pathway at the interface between *HNRNPK* and *TFAP2C* (Fig. 4C, 5A-E, Supp. Fig. 5B-F). *HNRNPK* silencing in PG127-infected HovL reproduced the effects of *HNRNPK* ablation in LN-229 C3 cells, showing reduced RNA levels of mTOR and Rptor, but not Rictor, a scaffold protein required for the formation of the mTORC2 complex (Fig. 7A). Accordingly, *HNRNPK* downregulation caused a consistent reduction in the phosphorylation of S6 and 4EBP1 (Fig. 7B). Thus, we wondered whether mTORC1 activity might mediate or contribute to the effects of *HNRNPK* on prion propagation.

**Figure 7.**
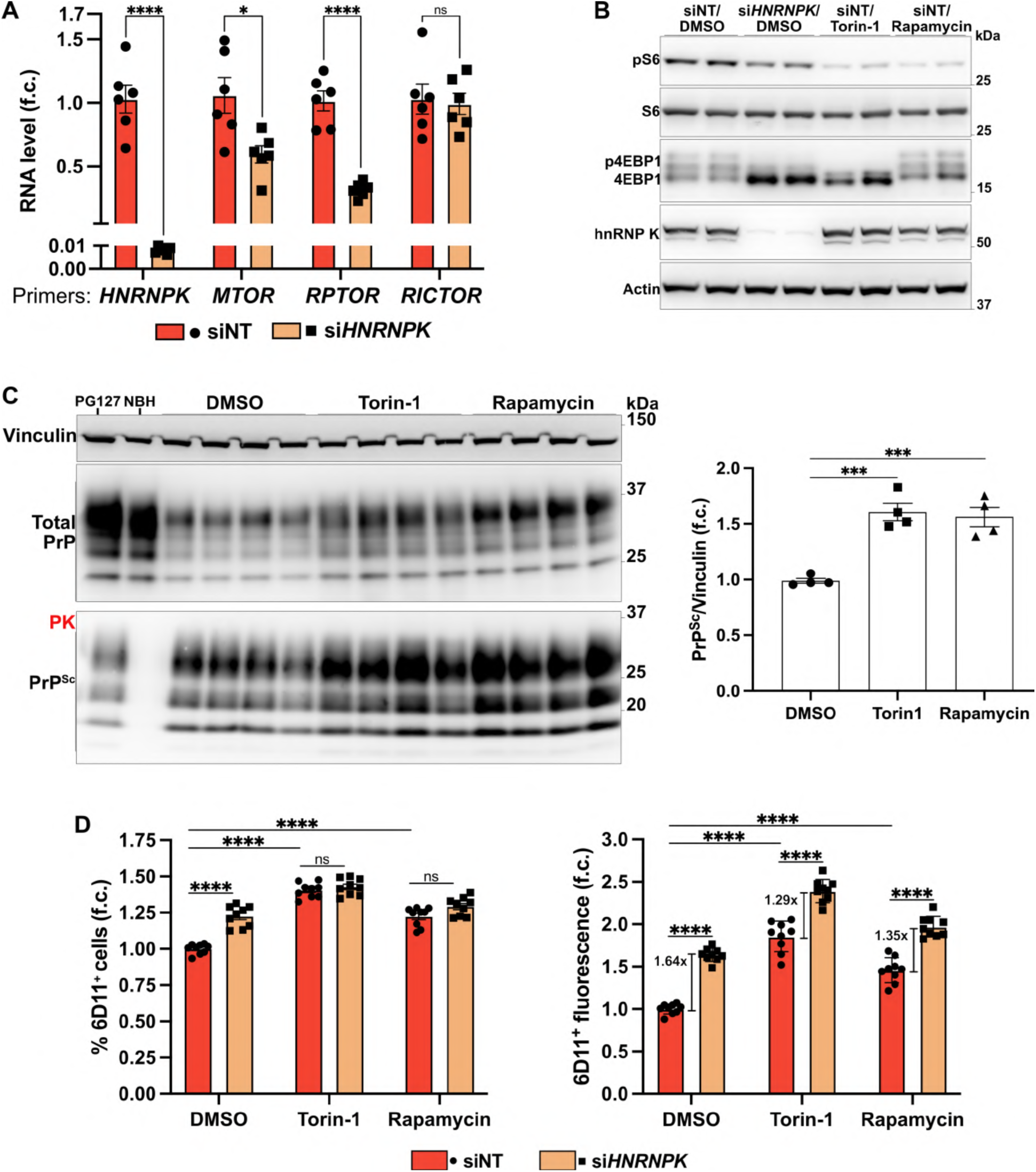
**A.** qRT-PCR showing *HNRNPK*, *MTOR*, *RPTOR*, and *RICTOR* RNA upon *HNRNPK* silencing (96 hours) in PG127-infected HovL cells. *n* = 6. **B.** 4EBP1 and S6 protein phosphorylation upon *HNRNPK* silencing (96 hours) or treatment with 500 nM Torin-1 or Rapamycin (72 hours) in PG127-infected HovL cells. *n* = 2. **C.** PK digested (bottom) and undigested (top) western blots showing, respectively, PrP^Sc^ and total PrP after treatment with 500 nM Torin-1 or Rapamycin (72 hours) in PG127-infected HovL cells. *n* = 4. **D.** Flow cytometry quantification of anti-PrP^Sc^ 6D11 antibody signal in PG127-infected HovL cells upon *HNRNPK* knockdown (96 hours) and treatment with 500 nM Torin-1 or Rapamycin (72 hours). *n* = 9. **Data information:** PG127-infected (PG127) and Not-infectious Brain Homogenate (NBH) were used for PK digestion positive and negative control, respectively. Non-targeting siRNA (siNT) and DMSO were used as controls. qRT-PCR results are normalized against *GAPDH* expression. *n* represents independent cultures. f.c.: fold change. Mean ± SEM. ns: *p* > 0.05, ^✱^: *p* < 0.05, ^✱✱✱^: *p* < 0.001, ^✱✱✱✱^: *p* < 0.0001 (Multiple Unpaired t-test Holm-Šídák method in A. One-way ANOVA Dunnett’s test in C. Two-way ANOVA Šídák’s test in D).

To test this hypothesis, we treated HovL cells with Torin-1 and Rapamycin, two potent mTORC1/2 inhibitors with different mechanisms of action. Torin-1 inhibits both mTORC1 and mTORC2 complexes, whereas short-term administration of Rapamycin blocks only mTORC1 (*52*). Additionally, Torin-1 strongly inhibits the phosphorylation of mTORC1 substrates S6 and 4EBP1, whereas Rapamycin has a less pronounced effect on 4EBP1 phosphorylation, which is often cell-type dependent (*52*) (Fig. 7B). After three days, both drugs caused a consistent increase in PrP^Sc^ levels, resembling the effects of *HNRNPK* downregulation (Fig. 7C, Supp. Fig. 7D).

To dissect the net contribution of mTOR inhibition and *HNRNPK* silencing on prion propagation, we treated PG127-HovL with *HNRNPK* siRNA for 24 hours, followed by Torin-1 or Rapamycin treatment for 72 hours. 6D11 immunostaining and flow cytometry analysis confirmed that *HNRNPK* silencing, Torin-1, and Rapamycin each significantly increased the total 6D11^+^ fluorescent signal and the fraction of 6D11^+^ cells (Fig. 7D). Co-treatment with *HNRNPK* siRNA and the mTOR inhibitors did not further increase the % 6D11^+^ cells compared to the effects of the drugs alone (Fig. 7D). However, the total 6D11^+^ fluorescence increased upon *HNRNPK* downregulation even in the presence of Torin-1 or Rapamycin (Fig. 7D). Nonetheless, the relative effect of *HNRNPK* silencing when combined with the drugs was still smaller than that mediated by *HNRNPK* siRNA alone (Fig. 7D). These findings suggest that *HNRNPK* influences prion propagation at least in part through mTORC1 signaling, although additional mechanisms may be involved.

## Discussion

Because of its role as a prominent limiter of prion propagation (*26*), the molecular pathways in which hnRNP K participates are important to prion biology and, in the best case, may even point to therapeutically actionable targets against prion diseases. We reasoned that an important first step would consist of clarifying the molecular basis of the cell-essentiality of hnRNP K. We opted to perform a CRISPR-based synthetic viability screen because this approach goes beyond the mere description of phenotypes associated with hnRNP K deficiency, and can instead point to genes that are causally involved in mediating the function of hnRNP K.

Large genome-wide perturbation screens assess the phenotypic outcomes of extensive collections of biological samples under various conditions. Consequently, background noise is frequently observed in these types of experiments, and our screen was no exception. We identified genes associated with mitochondrial membrane permeability and apoptosis, such as *FADD*, *MFN2*, *AIFM1*, and Elongator complex proteins (ELPs). These genes are likely to reflect outcomes that are not specific to *HNRNPK*, as the deletion of proapoptotic factors results in enhanced survival upon cellular stress (*34, 66–71*). However, we also recovered genes directly linked to *HNRNPK*, such as *PCBP1*, *PCBP2,* and *HNRNPA1,* which are members of the *HNRNPK* gene superfamily (*35, 36*). Furthermore, our screen enriched for *XRN2*, *NUDT21*, and *CPSF6*, which synergize with *HNRNPK* in modulating post-transcriptional RNA processes (*7, 37*). These results suggest that our experimental approach was effective in identifying true *HNRNPK* genetic interactors despite background noise and genetic confounders.

The Transcription Factor AP-2γ (*TFAP2C*) emerged from our genetic screen as the strongest modulator of *HNRNPK* essentiality. Its effect was bidirectional: Tfap2c removal conspicuously reduced the cell death triggered by *HNRNPK* ablation, whereas its overexpression hypersensitized cells to the loss of *HNRNPK*. Both hnRNP K and Tfap2c control the expression and localization of several lncRNAs (*9, 39, 45*), regulate different glucose metabolic pathways (*46–48*), and modulate neurodevelopment in mice and humans (*17, 18, 42, 43*). Because a direct functional interaction between these two genes had not been reported, we decided to investigate the molecular significance of their genetic link. We found that *HNRNPK* ablation perturbed the transcription of *TFAP2C*, although with opposite effects in different cell lines. Deletion or upregulation of *TFAP2C* did not change *HNRNPK* RNA and protein levels, disproving the presence of a regulatory feedback loop between these two genes and suggesting that hnRNP K acts upstream of Tfap2c. In addition, we found that hnRNP K and Tfap2c co-localize in the nucleus and co-immunoprecipitated from cellular extracts. Hence, multiple converging lines of evidence from forward genetics, immunostaining, and immunochemistry point to a functional connection between these two proteins.

The ablation of *TFAP2C* prevented the induction of caspase-dependent apoptosis triggered by the deletion of *HNRNPK*. However, this finding is not very informative because apoptosis can result from multiple dysfunctional processes, and its prevention does little to explain how Tfap2c mediates the action of hnRNP K. To gain more insight into the relevant cellular events, we performed early-stage RNA sequencing analyses after ablation of *TFAP2C, HNRNPK*, or both. We found that hnRNP K removal dysregulated cell bioenergetics, particularly impairing lipid and glucose metabolism, long before extensive cell death occurred. Instead, Δ*TFAP2C;*Δ*HNRNPK* double-ablated cells retained a more balanced transcriptional profile. Accordingly, we observed that *HNRNPK* ablation reduced intracellular ATP levels and increased autophagy flux by decreasing mTORC1 expression and promoting AMPK activation. Crucially, all these perturbations were largely corrected by Δ*TFAP2C;*Δ*HNRNPK* double ablation. *TFAP2C* deletion alone also perturbed the mTORC1 downstream pathway by diminishing 4EBP1 expression, whereas *TFAP2C* overexpression increased both S6 phosphorylation and 4EBP1 levels. We conclude that *HNRNPK* deletion causes a metabolic impairment leading to a nutritional crisis and a catabolic shift, whereas *TFAP2C* activation promotes mTORC1 anabolic functions. Thus, Tfap2c removal may rewire the bioenergetic needs of cells by modulating the mTOR signaling and augmenting their resilience to metabolic stress like the one induced by *HNRNPK* ablation.

These findings point to a previously unrecognized role of *HNRNPK*, i.e. its impact on cellular bioenergetics and metabolic regulation. Emerging evidence underscores the critical role of cellular energy homeostasis in protein misfolding disorders and neurodegeneration (*72–74*). Specifically, alterations in ATP levels and autophagy have been implicated in regulating the aggregation of pathogenic proteins (*58–63*). Thus, it is plausible that the metabolic imbalance resulting from *HNRNPK* ablation and linked to mTOR and AMPK dysregulation may influence prion propagation dynamics. Consistently, the anti-prion compound psammaplysene A (PSA) (*26*) is known to modulate the Foxo3a-mTOR-AMPK pathway (*75*), and PSA-treated cells upregulate genes related to glycolysis and energy metabolism, which are instead inhibited by *HNRNPK* knockdown (*26*).

Surprisingly, *TFAP2C* overexpression led to a marked reduction of aggregated pathogenic prions without altering the expression of their monomeric physiological precursor PrP^C^. This points to a PrP^C^-independent inhibitory effect of *TFAP2C* on prion propagation. Interestingly, *TFAP2C* upregulation prevented PrP^Sc^ accumulation even when *HNRNPK* was downregulated. Hence, *TFAP2C* operates in parallel or downstream to *HNRNPK* in a pathway influencing prion propagation. Since we found in mTORC1 a shared and inversely modulated pathway at the interface between *HNRNPK* and *TFAP2C*, we tested its role in prion propagation. Pharmacological inhibition of mTOR with Torin-1 and Rapamycin partially reproduced the effects of *HNRNPK* silencing on prion propagation, linking *HNRNPK* and mTORC1 signaling to the regulation of prion replication. However, mTOR inhibition did not explain the whole effect of *HNRNPK* on prion propagation, suggesting that, although involved, other mechanisms may further participate and contribute to such regulation.

These data suggest a role of *TFAP2C* and *HNRNPK* at the crossroads between mTOR cell metabolic control and prion propagation. One potential mechanism could involve the regulation of autophagy and ATP production mediated by mTORC1. Previous studies observed that mTOR inhibition and autophagy activation reduced, rather than increased, PrP^Sc^ aggregation (*76, 77*). However, most of these findings were based on long-term inhibition of mTOR. Acute inhibition, mimicking the time frame of *HNRNPK* and *TFAP2C* manipulation, may produce distinct, short-lived metabolic changes that outweigh the effects of autophagy on prion propagation.

In conclusion, the identification of *TFAP2C* as a novel regulator linked to *HNRNPK* points to functions of this protein in the modulation of mTOR cell metabolism and outlines a possible explanation for its role in prion propagation. From a methodological viewpoint, the present study shows that it is possible to use synthetic-survival CRISPR screens to discover novel functional actions of genes even when their removal causes cell lethality. The fundaments laid here may be instrumental to discovering further pathways regulated by *TFAP2C* and *HNRNPK* and their roles in the modulation of prion propagation and potentially additional proteinopathies.

## Materials and methods

### Cell culture

The LN-229 cell lines were cultured in Dulbecco’s Modified Eagle Medium (DMEM) (Gibco, 11965092) supplemented with 10% fetal bovine serum (FBS) (Clontech Laboratories) and 1% penicillin/streptomycin (P/S) (Gibco, 15140122). The U-251MG cell lines were grown in OptiMEM (Gibco, 31985047) supplemented with 10% FBS, 1% GlutaMax (Gibco, 35050061), 1% MEM Non-Essential Amino acids (MEM-NEAA) (Gibco, 11140050), and 1% P/S. The HEK-293T cell line for lentivirus production was cultured in DMEM supplemented with 10% FBS. For lentiviral delivery, all cells were seeded in antibiotic-free medium and placed under antibiotic selection 24 hours after transduction. All cells were maintained in culture at 37 °C in a 3% oxygen and 5% CO_2_ atmosphere.

### HovL cell line generation

The generation of the human ovinized LN-229 cell line, HovL, was performed as previously described for the ovinized SHSY-5Y cell line, HovS (*64*). LN-229 wild-type cells were co-transfected with two plasmids encoding Cas9 (pSpCas9(BB)-2A-GFP, addgene #48138) and qgRNAs against human *PRNP* (*32*) to achieve *PRNP* ablation. A monoclonal line was isolated via limiting dilution, and the ablation was confirmed by PCR and sequencing. Δ*PRNP* cells were then transfected using Lipofectamine 3000 (Invitrogen) with an expression vector (EF1α promoter) (Sigma-Aldrich, OGS606-5U) containing the *Ovis aries PRNP* VRQ allele coding sequence (Genescript). The insert was cloned with the human ER localization signal and optimized for codon usage in human cells. Cells were kept under geneticin selection (400 μg/ml) (Thermo Fisher Scientific), and a stable monoclonal line was isolated via limiting dilution. Chronic PG127 prion-infected or mock-infected HovL were obtained as previously reported (*64*). HovL were incubated either with PG127 prion-contaminated Brain Homogenate (PG127) or Non-infectious Brain Homogenate (NBH) and split for at least eight passages to dilute to original prion inoculum and enhance *de novo* PrP^Sc^ formation and propagation. PG127 and NBH HovL were also transduced with lentiCas9-Blast plasmid to stably express Cas9 (*78*). Primers’ sequences for PCR (Fwd: 5’-GCACTCATTCATTATGCAGGAAACA-3’, Rev: 5’-AGACACCACCACTAAAAGGGC-3’) and sequencing (5’-GGACTCTGACGTTCTCCTCTTC-3’).

### Immunoblotting

Cell extracts were prepared in lysis buffer (50 mM Tris–HCl, pH 8, 150 mM NaCl, 0.5% sodium deoxycholate, and 0.5% Triton-X 100) supplemented with protease inhibitors and PhosStop (Sigma-Aldrich). In case of Proteinase K (PK) (Roche AG) digestion, protease inhibitors and PhosStop were avoided. The bicinchoninic acid assay (BCA) was used to measure the total protein concentrations according to the manufacturer’s instructions (Pierce). Immunoblots were performed using standard procedures. The samples, boiled at 95 °C in NuPAGE LDS Sample Buffer 1x (Invitrogen) supplemented with 1 mM DTT (Sigma-Aldrich), were loaded onto precast gels (Invitrogen), and blotted onto PVDF membranes (Invitrogen). Proteinase K (PK) digestion was performed at a final concentration of 2.5 μg/ml for 30 minutes at 37°C. Following are the antibodies used and their relative dilutions: anti-Tfap2c 1:20000 (Abcam, ab76007), anti-hnRNP K 1:2000 (Abcam, ab70492), anti-Cas9 (*S. pyogenes*) (Santa Cruz Biotechnology, 7A9-3A3) 1:1000 (Cell Signaling Technology, 14697), anti-Puromycin 1:1000 (Kerafast, EQ0001), anti-PARP 1:1000 (Cell Signaling Technology, 9542S), anti-Cleaved Caspase 3 1:1000 (Cell Signaling Technology, 9661S), anti-Caspase 3 1:1000 (Novus Biologicals, NB100-56708), anti-GPX4 1:1000 (Abcam, ab125066), anti-mCherry 1:1000 (Abcam, ab213511), anti-LC3 1:1000 (Cell Signaling Technology, #2775), anti-Actin-HRP 1:10000 (Sigma-Aldrich, A3854), anti-Vinculin 1:2000 (Abcam, ab129002), anti-PrP POM1 300 ng/ml (for PrP^Sc^ detection) (*79*) or POM2 300 ng/ml (for total PrP or PrP^C^ detection) (*79*), anti-mTOR 1:1000 (Cell Signaling Technology, 29835S), anti-pAMPK 1:1000 (Cell Signaling Technology, 2535S), anti-AMPK 1:1000 (Cell Signaling Technology, 2532S), anti-pUlk1 1:1000 (Cell Signaling Technology, 6888T), anti-Ulk1 1:1000 (Cell Signaling Technology, 8054T), anti-4EBP1 1:1000 (Cell Signaling Technology, 9452S), anti-pS6 1:1000 (Cell Signaling Technology, 2215), anti-S6 1:1000 (Cell Signaling Technology, 2217), anti-Rabbit 1:5000 (Jackson ImmunoResearch, 111.035.045), anti-Mouse 1:5000 (BIO-RAD, STAR87P).

### Lentivirus production

All the lentiviral vectors were produced as follows: HEK-293T cells were seeded at 40% confluency, and 24 hours later the target plasmid was co-transfected together with the pCMV-VSV-G (Addgene plasmid # 8454) (*80*) and psPAX2 plasmids (Addgene plasmid # 12260) using Lipofectamine 3000 transfection reagent (Invitrogen, ThermoFisher Scientific). After 6 hours, the medium was replaced with DMEM supplemented with 10% FBS and 1% HyClone Bovine Serum Albumin (Cytiva). 72 hours after the transfection, the supernatant was collected, centrifuged at 1500 RCF at 4°C for 5 minutes, filtered through a 0.45 μm filter (Whatman), aliquoted, and stored at −80°C. Viral titer was measured as previously reported (*81*). Cells were seeded at known numbers, infected with different volumes of lentivirus, and 24 hours later selected with the relevant antibiotics. The lentiviral Titer Unit (TU) was extrapolated by measuring the fraction of viable cells with CellTiter-Glo 2.0 and GloMax Plate reader (Promega). For plasmids bearing a fluorescent probe, the viral titer was measured by flow cytometry: briefly, cells were seeded in a 24-well plate and infected with different volumes of lentivirus after 6 hours; 72 hours after the infection, the cells were harvested, and the percentage of fluorescent positive cells was acquired by flow cytometry (BD LSRFortessa, Cell Analyzer) and analyzed by FlowJo 10 (Tree Star).

### Whole-genome CRISPR-Cas9 screen and analysis

The human Brunello CRISPR ablation pooled library (Addgene #73178) (*33*) was amplified as previously reported (*81*), packaged into a lentiviral vector, and titrated as described above. The library was transduced into 280 million LN-229 C3 cells with a multiplicity of infection (MOI) of 0.3 at an estimated coverage of around 1100 cells expressing each sgRNA. The screen was performed as follows. Day 0: 1 million cells/ml were seeded in a final volume of 31.25 ml per flask in 9 T-300 flasks. Cell density was defined based on the original titration of the lentiviral packaged Brunello Library. 6 hours later, the cells were transduced with the packaged library. Day 1: 24 hours after the delivery of the library, half of the culture (280 million cells) was transferred into new T-300 flasks in DMEM supplemented with 15 μg/ml blasticidin and 1 μg/ml puromycin (Thermo Fisher Scientific). The other half (280 million cells) was harvested, pelleted, and frozen at −80°C. Day 5: At this point, all the uninfected cells were depleted, and 80 million cells were re-seeded to maintain a library coverage >1000x. Day 7: 160 million cells were seeded at a concertation of 1 million cells/ml. 6 hours later, half of the culture was transduced either with qgRNAs against the *HNRNPK* gene or with the non-targeting control qgRNAs (NT) at an MOI of 10. Day 8: 24 hours after the delivery of the *HNRNPK* or NT qgRNAs, 160 million cells were transferred into 20 T-300 flasks and maintained under selection with 15 μg/ml blasticidin, 1 μg/ml puromycin and 1.5 mg/ml geneticin (Thermo Fisher Scientific). *HNRNPK*-targeted cells were incubated for six more days, changing the medium every three days. Day 11: To avoid over-confluency, the cells transduced with the NT qgRNAs were split according to the library coverage. Day 14: 80 million cells from both conditions were harvested, pelleted, and frozen at −80°C.

The extraction of the genomic DNA (gDNA), the preparation, and the purification of the NGS library were performed as previously reported (*81*). Briefly, the gDNA was harvested from each of the collected samples using the Zymo Research Quick-gDNA MidiPrep (Zymo Research) according to the manufacturer’s protocol. To amplify the sgRNAs for NGS, a PCR reaction was set up for 23 cycles in 96 PCR plates (Sarstedt). Each gDNA sample was processed by using a mix of eight P5 primers in combination with one unique P7 primer. For the purification of the amplified product, the PCR reaction was mixed with five volumes of DNA Binding Buffer (Zymo Research), transferred in the Zymo-Spin V column with Reservoir (Zymo Research), and centrifuged at 500 RCF for 5 minutes at room temperature. The column was washed twice with 2 ml DNA Wash Buffer (Zymo Research) by a second step of centrifugation. The column was transferred to a 2 ml collection tube and spun again at maximum speed for 1 minute to remove residual wash buffer, and finally, the purified PCR reaction was eluted with Nuclease-Free Water (Invitrogen). The PCR product was quantified from agarose gel, and the samples sequenced by Illumina Novaseq 6000 Full SP Flowcell.

The Sushi data analysis platform (*82*) (FGCZ, University of Zurich) or R (version 3.5.2) were utilized for data analysis and graphical visualizations. Reads quality was assessed using FastQC. Reads were aligned to the Human Brunello CRISPR ablation pooled library (*33*) with targeted gene symbols. Read counts were generated using the featureCounts function (*83*) from the Rsubread package in R. A generalized linear model applying Trimmed Means of M-values (TMM) normalization was implemented using the EdgeR package in R (*84*) for differential expression analysis. Clustering analysis was performed with the hclust function from the R stats package. Differential gene enrichment was performed using Edge R (version 3.5.2) by providing the log_2_ fold change and false discovery rate (FDR). sgRNAs have been grouped according to their targeted gene for the generation of the final gene lists.

### Essential genes identification

To identify the set of essential genes in LN-229 cells, the Cancer Dependency Map’s “2021-Q3” release (*49*) served as the primary resource. Genes whose “gene_effect” scored below −1 were classified as essential. The gene symbols were aligned with those referenced in the Brunello Library. The genes missing in the Brunello Library were excluded from the final list and the following analysis (Supp. Table 2).

### CRISPR ablation and activation qgRNAs

Unless otherwise specified, the effects of CRISPR ablation were always analyzed 7 days after transduction of ablation qgRNAs in LN-229 Cas9 cells, and after 10 days in U-251MG Cas9 cells. CRISPR activation was always evaluated five days after delivery of transactivating qgRNAs.

*HNRNPK* intron-targeting sgRNAs were designed with a customized algorithm, and the specificity and efficacy metrics were calculated using the GPP sgRNA designer (*33, 85*), GuideScan (*86*), and CRISPOR (*87*) tools. The selected guides were then cloned inside the pYJA5_G418 backbone (*88*), a modified version of the pYJA5 vector (*32*), containing the geneticin resistance marker instead of puromycin resistance. The sequence of the intron-targeting sgRNAs and their score parameters are reported in Supp. Table 1.

The *HNRNPK* ablation qgRNAs and its non-targeting control (NT) were provided by Jiang-An Yin (*32*) and cloned into the pYJA5_G418 backbone. All the other ablation, activation, and control qgRNAs were cloned in the pYJA5, stocked in-house and kindly provided by Jiang-An Yin (*32*) in the form of glycerol stocks. For double-ablation experiments, cells were always co-treated with pYJA5_G418 and pYJA5 plasmids expressing *HNRNPK* or NT and *TFAP2C* or NT qgRNAs in the four possible combinations.

The non-targeting control NT qgRNAs used for CRISPR ablation and CRISPR activation are listed as *Control_3* and *Control_13* (for the ablation) and *Control_5* (for the activation) in Jiang-An Yin et al. (*32*).

### Cell survival analysis

For the clonogenic assay, cells were seeded in 6-well plates (3 x 10^5^ U-251 MG cells and 1 x 10^5^ LN-229 cells per well) and treated with qgRNAs against different genes the day after. The cells were incubated for different time points, washed twice with PBS, fixed with 4% paraformaldehyde (Thermo Fisher Scientific), stained with 0.5% crystal violet (Sigma-Aldrich) for 1 hour, washed three times with PBS, and dried before imaging.

For cell viability assay, cells were seeded in 96-well plates (1 x 10^4^ U-251 MG cells and 3 x 10^3^ LN-229 cells per well) and treated with drugs or qgRNAs against different genes the following day. The cells were incubated for different time points and the viability was measured using CellTiter-Glo 2.0 and GloMax Plate reader (Promega).

### Translation activity measurement

Translation activity was measured by puromycin labeling as previously described (*89*). Briefly, cells were pulse-chased with 10 µg/ml puromycin (Thermo Fisher Scientific) for 10 minutes and washed three times with PBS before harvesting. Proteins were immunoblotted with anti-Puromycin 1:1000 (Kerafast, EQ0001).

### RNA preparation and qRT-PCR

RNA was isolated using the RNeasy Mini kit (Qiagen). RNA quality and concentration were measured with a NanoDrop spectrophotometer (Thermo Fisher Scientific). Reverse transcription was carried out with the Quantitect Reverse Transcription kit (Qiagen) as per the manufacturer’s guidelines. For each sample, 10 ng of cDNA was loaded into 384-well PCR plates (Life Systems Design) in triplicates, and the detection was conducted using SYBR green (Roche). Readout was executed with ViiA7 Real-Time PCR systems (Thermo Fisher Scientific). The ViiA7 Real-Time PCR system (Thermo Fisher Scientific) was used for the readout, and the qRT-PCR data were processed using the 2^-ΔΔCT^ method. Following are the primers’ sequences: *HNRNPK* (Fwd: 5’-TTCAGTCCCAGACAGCAGTG-3’, Rev: 5’-TCCACAGCATCAGATTCGAG-3’), *TFAP2C* (Fwd: 5’-GCCGTAATGAACCCCACTGA-3’, Rev: 5’-TTCTTTACACAGTTGCTGGGC-3’), *GAPDH* (Fwd: 5’-TGCACCACCAACTGCTTAGC-3’, Rev: 5’-GGCATGGACTGTGGTCATCAG-3’), *MTOR* (Fwd: 5’-AAAGAGCAGAGTGCCCGCAT-3’, Rev: 5’-TCC AGG CCA CTA ACC TGT GC-3’), *ovPRNP* (Fwd: 5’-AACCACCACAAAGGGCGAGA-3’, Rev: 5’-GCACATCTGCTCCACCACTC-3’), *RPTOR* (Fwd: 5’-CTCGCAGTGGACAGCTCGTG-3’, Rev: 5’-CACGGCGAGAATGAAAGCCG-3’), *RICTOR* (Fwd: 5’-TGGATCTGACCCGAGAACCT-3’, Rev: 5’-TCCTCATAGTGAAAGCCCAGTT-3’).

### Immunofluorescence imaging

For immunofluorescence imaging, cells were seeded and grown in a 96-well plate or 8-well Culture Slide (Corning) and fixed in 4% paraformaldehyde for 20 minutes at room temperature. After three washes in PBS, the cells were blocked and permeabilized for 1 hour in FBS 10% supplemented with Triton X-100 0.2%. Cells were then incubated with primary antibodies diluted in the blocking/permeabilizing solution for 2 hours at room temperature. Cells were washed in PBS three times before incubation with the indicated secondary antibody for 1 hour at room temperature. After three more washes, cells were stained with DAPI diluted 1:10000 (Sigma-Aldrich) for 10 minutes at room temperature, washed again in PBS, and mounted. 8-well Culture Slide were imaged by Confocal Laser Scanning microscopy (Leica Stellaris 5 inverse), while 96-well plate images were captured by InCell Analyzer 2500HS high-throughput imaging microscope using a 20x water objective with 2x digital camera pixel binning for a total of 12 fields of view per well. Fluorescence per well was quantified with a custom-made CellProfiler pipeline and normalized to the total number of nuclei per well.

Following are the antibodies used and their relative dilutions: anti-Tfap2c 1:500 (Abcam, ab76007), anti-hnRNP K 1:500 (Abcam, ab39975), anti-Rabbit Alexa488 1:500 (Invitrogen, A-11008), anti-mouse Alexa488 1:500 (Invitrogen, A-11001), anti-Mouse Alexa647 1:500 (Invitrogen, A-21236).

### RNA Sequencing

RNA was isolated with the RNeasy Mini Kit (Qiagen). The Illumina Truseq Total RNA protocol (ribosomal depletion) was used for the preparation of the libraries. The quality of both the RNA and the libraries was assessed using the Agilent 4200 TapeStation System (Agilent). Libraries were pooled equimolar and sequenced on an Illumina NovaSeq 6000 sequencer (single-end 100 bp), achieving an approximate depth of 40 million reads per sample. The experiment was conducted in triplicate, however, due to low RNA quality, downstream analysis was performed only for two replicates except for the *TFAP2C*;*HNRNPK* double-knockout condition for which three replicates could be analyzed. Transcript alignment was carried out using the STAR package. After normalizing the library data (Transcripts Per Million, TPM), we conducted differential gene expression analysis using the DESeq2 package (*90*). We identified upregulated and downregulated genes considering the log_2_ fold change and Bonferroni-corrected *p*-value.

### RNAi silencing

HovL cells were seeded at a density of 2 x 10^5^ cells/well in 6-well plates. The next day, culture media was exchanged with OptiMEM (Gibco) without antibiotics, and siRNAs pre-mixed with RNAiMAX (0.3% final concentration) (Thermo Fisher Scientific) were delivered in a dropwise manner at a final concentration of 10 μM. Cells were incubated for 72 or 96 hours. Media containing siRNAs was aspirated, and cells washed once with PBS and then lysed. Pools of three different siRNAs were used to downregulate *HNRNPK* (s6737, s6738, s6739. Thermo Fisher Scientific, #4392420).

### Co-immunoprecipitation

Cell extracts were prepared in PBS supplemented with protease inhibitors (Sigma-Aldrich) using mechanical disruption. BCA assay was used to measure the total protein concentrations according to the manufacturer’s instructions (Pierce). The protein sample was processed in Protein LowBind Tube (Eppendorf) during each step. For each condition, 1 or 3 mg of protein was incubated for 1 hour at 4°C with 10 μl pre-washed dynabeads (Invitrogen) to clear the lysate from unspecific interactions. The cleared lysate was then incubated overnight at 4°C with 50 μl antibody-conjugated dynabeads and the sample kept mixing on a spinning wheel. The day after, the flow-through was collected and the dynabeads washed three times in PBS supplemented with protease inhibitors. The samples, boiled at 95 °C in LDS 2x (Invitrogen) supplemented with 2 mM DTT (Sigma-Aldrich), were loaded onto a 10% gel (Invitrogen), and blotted onto a PVDF membrane (Invitrogen). Following are the antibodies used and the relative dilutions: anti-Tfap2c 1:30 (Abcam, ab218107), anti-hnRNP K 1:1000 (Abcam, ab39975), anti-Tfap2c 1:5000 (Abcam, ab76007), anti-Actin HRP 1:10000 (Sigma-Aldrich, A3854), Normal Rabbit IgG (Merck, 12-370) diluted to match the concentration of anti-Tfap2c (Abcam, ab218107), VeriBlot 1:10000 (Abcam, ab131366).

### Lipid peroxidation detection

Lipid peroxidation was measured by flow cytometry using Liperfluo (Dojindo, L248-10) according to the manufacturer’s instructions. Δ*TFAP2C* and NT-unmodified LN-229 C3 cells were seeded in a 24-well plate, transduced with NT or *HNRNPK* ablation qgRNAs, and the following day put under antibiotic selection for three more days. On day 4 the cells were stained with Liperfluo, detached, acquired by flow cytometry (BD LSRFortessa, Cell Analyzer), and analyzed with FlowJo 10 (Tree Star).

### ATP level quantification

Intracellular ATP was measured via CellTiter-Glo 2.0 and GloMax Plate reader (Promega) and normalized to total protein synthesis by BCA assay (Pierce). Δ*TFAP2C* and NT-unmodified LN-229 C3 cells were seeded in a 96-well plate, transduced with NT or *HNRNPK* ablation qgRNAs, and the following day put under antibiotic selection for three more days. On day 4, the cells were lysed, and protein and ATP levels quantified as specified above.

### 6D11 staining for imaging and flow cytometry

The immunostaining protocol was based on a previous publication (*65*). For imaging, cells were washed with PBS and fixed with 4% paraformaldehyde for 12 minutes. After PBS washing, cells were incubated with 3.5M guanidine thiocyanate (Sigma-Aldrich) for 10 minutes. Following, cells were washed five times with PBS and incubated with mouse monoclonal (6D11) anti-PrP antibody (BioLegend, 808001) diluted 1:1000 in 25% Superblock (Thermo Fisher Scientific) at room temperature for 1 hour. After washing once with PBS, cells were labeled with secondary antibody (dilution 1:1000, anti-Mouse Alexa488, Invitrogen, A-11001) along with DAPI (Sigma-Aldrich) in 25% Superblock for 1 hour at room temperature. Cells were washed once with PBS and imaged with Fluoview FV10i confocal microscope (Olympus Life Science).

For flow cytometry, HovL cells were dissociated and fixed using the Cyto-Fast™ Fix/Perm buffer set (BioLegend, 426803) according to the manufacturer’s instructions. Additionally, after fixation, samples were incubated in 3.5M guanidine thiocyanate (Sigma-Aldrich) for 10 minutes and then immediately washed with 1 ml 1x Cyto-Fast™ Perm wash solution to ensure PrP^Sc^-specific binding (*65*). Staining was performed using AlexaFluor®-647 mouse monoclonal anti-PrP antibody (6D11) (BioLegend, 808008) diluted 1:200 in 1x Cyto-Fast™ Perm wash solution. Data were acquired on SP6800 spectral analyzer (Sony Biotechnology Inc), and analysis was performed using FlowJo 10 (Tree Star).

### Plasmids

pCMV-VSV-G was a gift from Bob Weinberg (Addgene plasmid # 8454; http://n2t.net/addgene:8454; RRID:Addgene_8454) (*80*). psPAX2 was a gift from Didier Trono (Addgene plasmid # 12260; http://n2t.net/addgene:12260; RRID:Addgene_12260). pXPR_011 was a gift from John Doench & David Root (Addgene plasmid # 59702; http://n2t.net/addgene:59702; RRID:Addgene_59702) (*91*). Human Brunello CRISPR knockout pooled library was a gift from David Root and John Doench (Addgene # 73178) (*33*). pXPR_120 was a gift from John Doench & David Root (Addgene plasmid # 96917; http://n2t.net/addgene:96917; RRID:Addgene_96917) (*92*). lentiCas9-Blast was a gift from Feng Zhang (Addgene plasmid # 52962; http://n2t.net/addgene:52962; RRID:Addgene_52962) (*78*). HA-tagged or untagged full-length *HNRNPK* plasmids were a gift from Ralf Bartenschlager (*93*). TFORF1330 was a gift from Feng Zhang (Addgene plasmid # 143950; http://n2t.net/addgene:143950; RRID:Addgene_143950) (*94*). TFORF3550 was a gift from Feng Zhang (Addgene plasmid # 145026; http://n2t.net/addgene:145026; RRID:Addgene_145026) (*94*). pSpCas9(BB)-2A-GFP (PX458) was a gift from Feng Zhang (Addgene plasmid # 48138; http://n2t.net/addgene:48138; RRID:Addgene_48138) (*95*). PSF-EF1-UB-NEO/G418 ASCI - EF1 ALPHA PROMOTER G418 SELECTION PLASMID (Sigma-Aldrich, OGS606-5U).

### Drugs

Erastin (Merck, E7781-1MG); Baicalein (Merck, 465119-100MG); Liproxstatin-1 (MedChemExpress, HY-12726); Ferrostatin-1 (MedChemExpress, HY-100579); Staurosporin (Abcam, ab120056); Z-VAD(Ome)-FMK (Cayman Chemical, Cay14463-1); Rapamycin (Selleckchem, S1039); Torin-1 (Sigma-Aldrich, 475991-10MG).

### Data availability

Raw sequencing data and processed data from this manuscript are available via GEO accession number GSE279797.

## Authors’ contributions

S.S.: designed, performed, contributed, supervised, and coordinated all the experiments and analysis; generation of the LN-229 C3 cell model; data curation and visualization; manuscript writing. D.C.: performed bioinformatic analysis, data curation, and visualization of the CRISPR screen and RNA-seq results (Fig. 1F, Fig. 4A-B, Supp. Fig. 2B-F, Supp. Table 2 to 5); assisted in cell culture for the generation of LN-229 C3 cell model; experimental design and manuscript writing. M.B.: performed or contributed to all the experiments illustrated from Fig. 3 to 6, and in Supp. Fig. 3, 5, and 7A; performed co-immunoprecipitation assay (Fig. 2H-I, Supp. Fig. 3L); data curation and visualization; manuscript writing. G.M.: contributed to developing the 6D11 staining protocol; performed the 6D11 staining and flow cytometry analysis, data curation, and visualization (Fig. 6D, Supp. Fig. 7D); contributed to the experiments regarding prion propagation (Fig. 6, Supp. Fig. 6-7); contributed to the generation of the HovL cell model (Supp. Fig. 6); manuscript writing. M.C.: assisted with confocal image acquisition (Supp. Fig. 3J-K); performed flow cytometry analysis for isolation of the LN-229 C3 cell line (Supp. Fig. 1B); data curation and visualization; manuscript writing. L.F.: design of *HNRNPK* intron-targeting sgRNAs (Supp. Fig. 1C-D, Supp. Table 1); CRISPR screen data analysis, curation, and visualization (Fig. 1F, Supp. Fig. 2C); manuscript writing. V.B.: contributed to developing the 6D11 staining protocol; HovL cells staining (Supp. Fig. 7B); manuscript writing. C.O.M.: generation of the HovL cell model (Supp. Fig. 6); assisted in cell culture; data curation and visualization. D.L.V.: experimental design; assisted in cell culture; contributed to genomic DNA purification and sgRNA PCR amplification. S.N.: experimental design; assisted in cell culture; contributed to genomic DNA purification and sgRNA PCR amplification. F.B.: performed flow cytometry analysis, data curation, and visualization (Supp. Fig. 4C). K.G.: generation of the LN-229 dCas9-VPR model; experimental design; assisted in cell culture; manuscript writing. J-A.Y.: designed and produced CRISPR qgRNAs. E.D.C.: contributed to developing the 6D11 staining protocol and flow cytometry data collection; experimental design and manuscript writing. A. A (Andrea Armani).: supervision, design, and technical support for the experiment related to mTOR, AMPK, ULK1, and autophagy (Fig. 5; Supp. Fig. 5); RNA-seq data curation and visualization (Fig. 4A-B); experimental design and manuscript writing. A.A. (Adriano Aguzzi): proposed the project, supervised its execution, secured funding, and helped to write and correct the manuscript.

## Funding and acknowledgments

A.A. is supported by institutional core funding by the University of Zurich, a Distinguished Scientist Award of the NOMIS Foundation, and grants from the Michael J. Fox Foundation (grant ID MJFF-022156 and grant ID MJFF-024255), the Swiss National Science Foundation (SNSF grant ID 179040 and grant ID 207872, Sinergia grant ID 183563), the GELU Foundation, swissuniversities (CRISPR4ALL), the Human Frontiers Science Program (grant ID RGP0001/2022), and Parkinson Schweiz. S.S. was supported by UZH Candoc Grant 2022 (No. FK-22036) for the development of this project. D.C. and V.B. were supported by the UZH Candoc Grant 2022 and 2023, respectively. K.G. was the recipient of a grant from the Michael J. Fox Foundation (MJFF-022156). J-A.Y. was the recipient of the UZH postdoc grant 2021 and the Career Development Awards grant of the Synapsis Foundation - Alzheimer Research Switzerland ARS. E.D.C. is the recipient of the Synapsis Career Development Award (2022-CDA01) and the Parkinson Schweiz grant. Andrea A. is the recipient of a Marie Curie postdoctoral fellowship (MSCA GF, 101033310). We thank Dr. Merve Avar and Dr. Daniel Heinzer for their supervision in the Aguzzi’s lab and their initial identification of the role of *HNRNPK* in prion propagation, which was the starting point for this project (*26*). We also thank Dr. Emina Lemes for the help setting up the screening strategy and protocol, Dr. Tingting Liu for support during the amplification of the Brunello library, Dr. Marian Hruska-Plochan, Carolina Appleton, Ana Marques, Lorenzo Maraio, and Marigona Imeri for laboratory assistance, critical discussions, and technical help. Dr. Susanne Kreutzer, Dr. Maria Domenica Moccia, and the Functional Genomics Center Zurich (FGCZ) for preparing sequencing libraries, CRISPR screen NGS sequencing, RNA sequencing, quality control, and technical support, Dr. Aria Maya Minder Pfyl, Silvia Kobel and the Genomic Diversity Centre (GDC) of the ETH Zurich for RNA quality control and technical advice. The funders of the study had no role in study design, data collection, data analysis, data interpretation, or writing of the report. The corresponding author had full access to all the data in the study and had final responsibility for the decision to submit for publication. GSEA was run with GSEA 4.3.2 Broad Institute software (*96, 97*). Gene enrichment analysis was performed using ShinyGO 0.77 and 0.8 (*98*) online tools. Statistical analysis and graphs were generated with R (version 3.5.2) and GraphPad Prism version 10.2.0 for Windows, GraphPad Software, Boston, Massachusetts USA, www.graphpad.com. Fig.1E was created with <u>BioRender.com</u>.

---

**Supplementary Figure 1.**
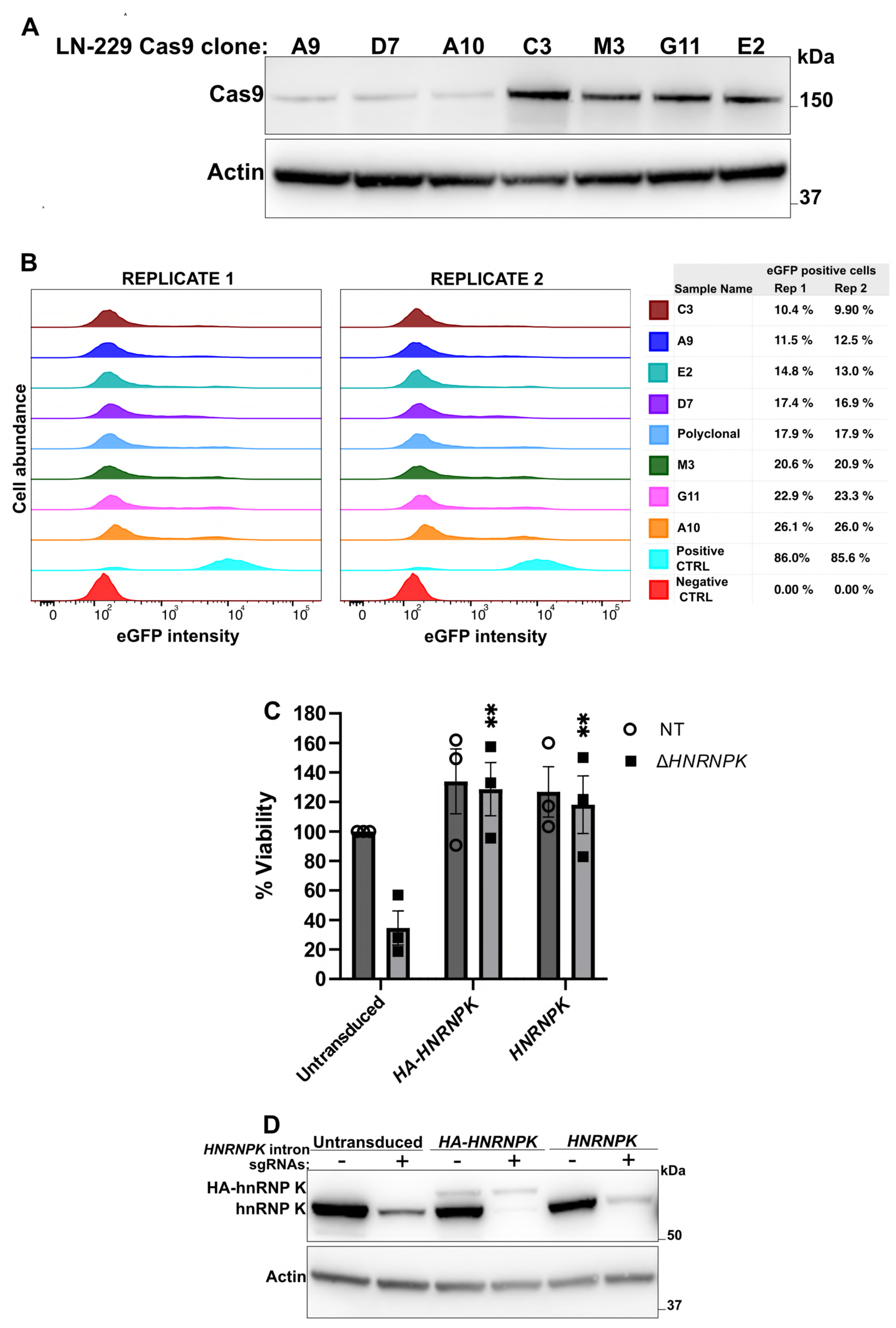
**A.** Cas9 protein in isolated LN-229 Cas9 clones. **B.** Flow cytometry-based determination of Cas9 activity in the LN-229 Cas9 clones by an *eGFP* reporter assay. Cas9 activity was estimated from the percentage of the eGFP-negative cells. LN-229 not expressing Cas9 or the *eGFP* reporter were used as positive and negative controls, respectively. **C.** Viability of LN-229 C3 cells upon ablation of the *HNRNPK* endogenous gene (CellTiter-Glo assay). LN-229 C3 cells expressed a vector carrying either the *HA-HNRNPK* or the *HNRNPK* coding sequence. Untransduced cells were used for control. Results are normalized on the untransduced non-targeting condition (NT). *n* = 3. **D.** The western blot refers to the data shown in C. -: NT, +: *HNRNPK* sgRNAs. **Data information:** *n* represents independent cultures. Mean ± SEM. ^✱✱^: *p* < 0.01 (Two-way ANOVA Dunnett’s test).

**Supplementary Figure 2.**
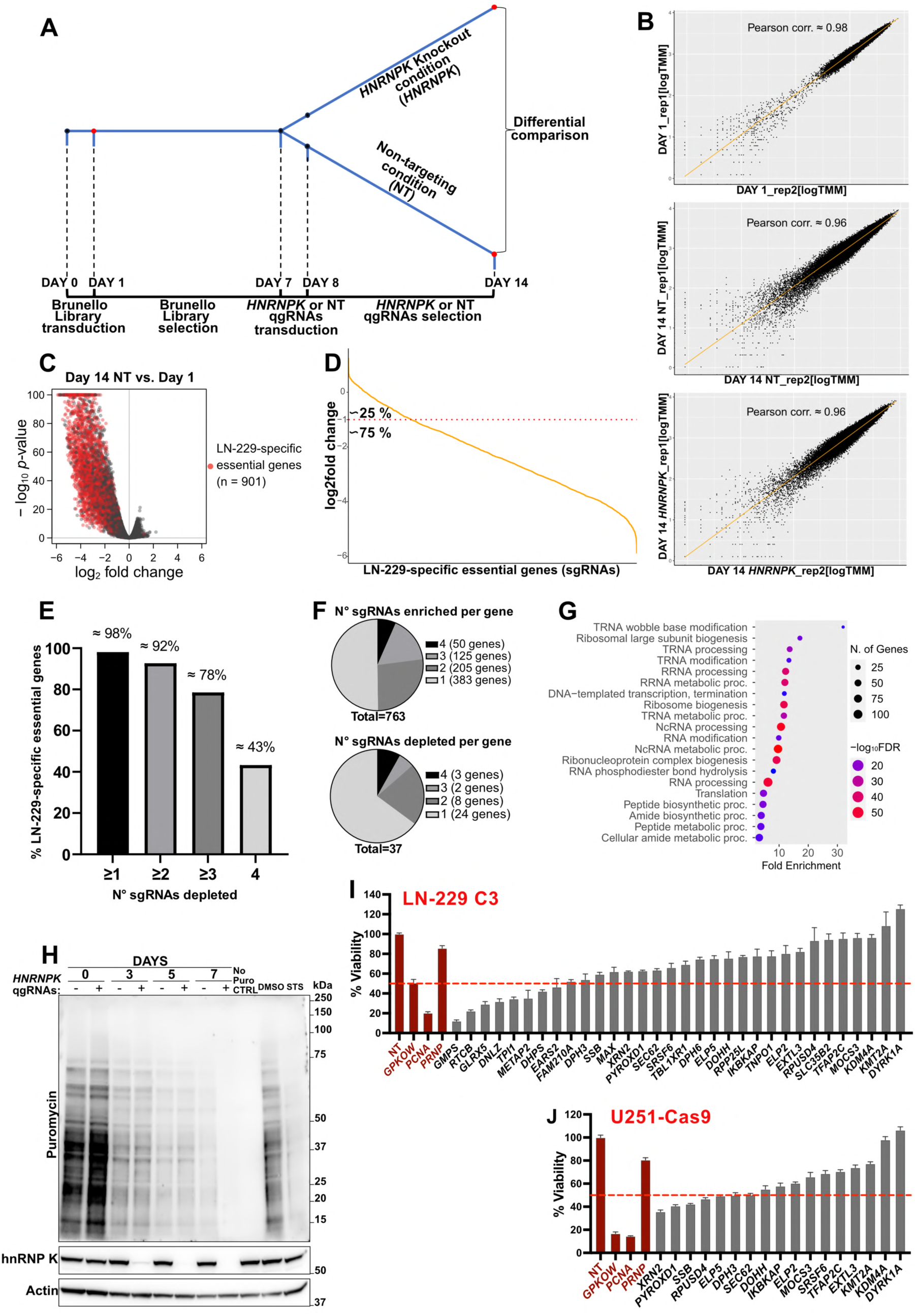
**A.** Workflow of the genome-wide CRISPR deletion screen. The red dots highlight the three time points subjected to next-generation sequencing (NGS) and analysis (Day 1, Day 14 NT, Day 14 *HNRNPK*). **B.** Correlation between the two experimental replicates of the screen for the three analyzed conditions: Day 1, Day 14 NT, and Day 14 *HNRNPK* **C.** Volcano plot showing the differential sgRNAs abundance in Day 14 NT vs. Day 1. Red-filled circles indicate the sgRNAs targeting LN-229 essential genes. **D.** Distribution of sgRNAs targeting LN-229 essential genes in the Day 14 NT vs. Day 1 comparison. **E.** Percentage of LN-229 essential genes with at least one, two, three, or four sgRNAs depleted in the Day 14 NT vs. Day 1 comparison. **F.** Distribution of the number of sgRNAs per gene significantly enriched or depleted. **G.** Gene enrichment biological process analysis of the genes with ≥2 sgRNAs enriched in *HNRNPK* vs. NT at day 14. **H.** Puromycin labeling and detection of global protein synthesis in LN-229 C3 cells transduced with *HNRNPK* (+) or NT (-) qgRNAs. 4 hours, 1 μM staurosoprine (STS) was used for control. **I, J.** Cell viability upon individual deletion of each of the candidate genes obtained from the screen (CellTiter-Glo assay). Results are normalized against the seeded cell density and compared to the NT condition. Red columns indicate the control groups: non-targeting control (NT), non-specific genes (*PCNA*, *GPKOW*, *PRNP*). The red dashed line highlights the viability threshold set at 50% of the NT condition. Mean ± SEM, *n* ≥ 3 independent cultures.

**Supplementary Figure 3.**
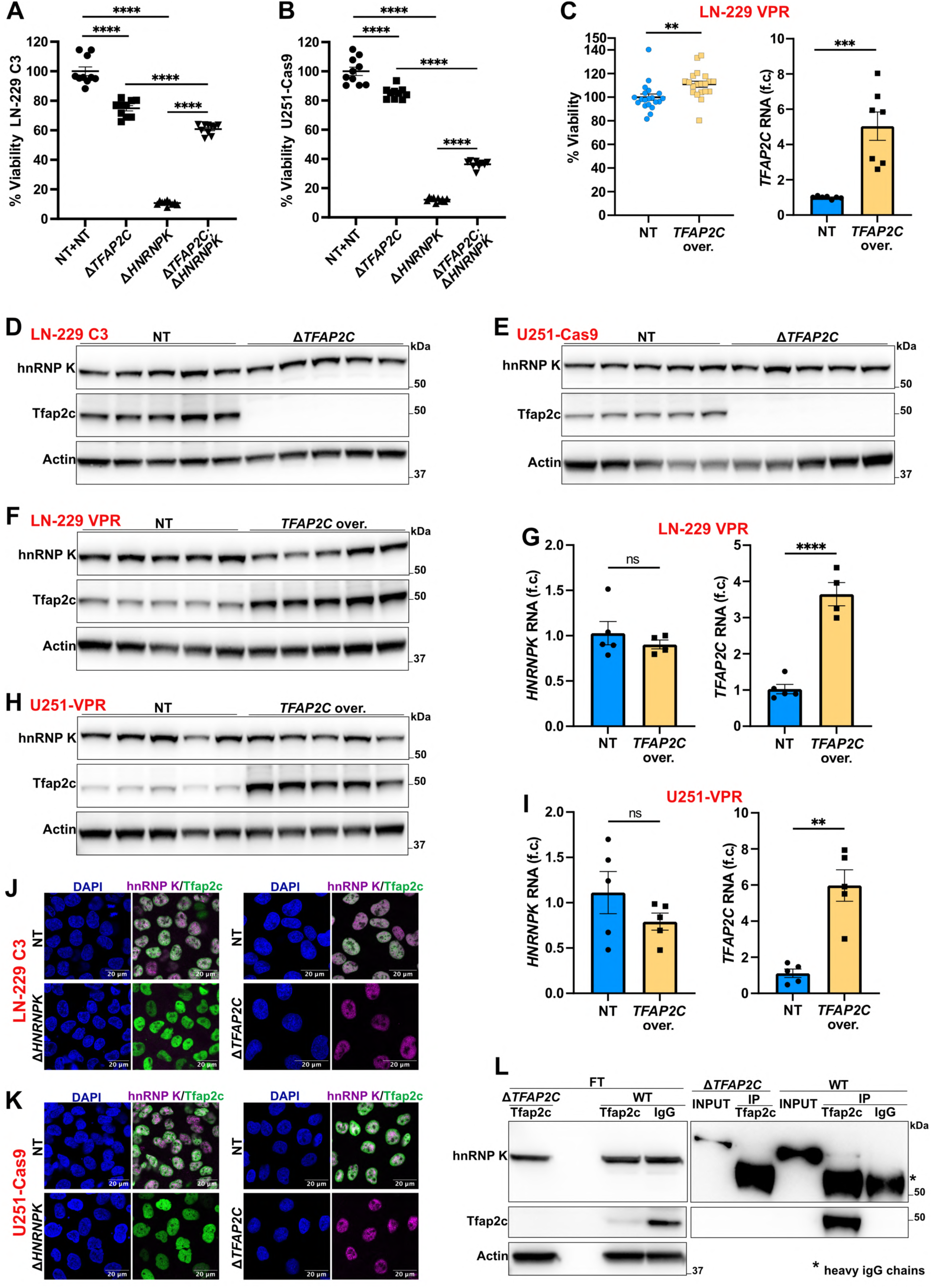
**A-B.** Viability of Δ*TFAP2C* or NT-unmodified cells 7 (A) and 10 days (B) upon delivering *HNRNPK* or NT qgRNAs (CellTiter-Glo assay). Results are normalized against the seeded cell density before *HNRNPK* ablation. *n =* 10. **C.** Viability of LN-229 dCas9-VPR cells upon *TFAP2C* overexpression (CellTiter-Glo assay). *n* = 20. qRT-PCR: *n* = 7. **D-E.** hnRNP K protein upon *TFAP2C* ablation. *n* = 5. **F-I.** hnRNP K protein (F, H) and RNA (G, I) after *TFAP2C* overexpression in dCas9-VPR cells. WB: *n* = 5. qRT-PCR: *n* ≥ 4. **J-K.** Confocal images showing hnRNP K and Tfap2c proteins in Δ*TFAP2C* and NT-unmodified cells. hnRNP K and Tfap2c proteins were also imaged in cells transduced with *HNRNPK* or NT qgRNAs for 4 (J) or 6 (K) days. **L.** Co-immunoprecipitation of Tfap2c and hnRNP K in Δ*TFAP2C* and WT LN-229 C3 cells. IP: Immunoprecipitated Protein; FT: Flow Through after immunoprecipitation. **Data information:** qRT-PCR results are normalized against *GAPDH* expression. *n* represents independent cultures. f.c.: fold change. Mean ± SEM. ns: *p* > 0.05, ^✱✱^: *p* < 0.01, ^✱✱✱^: *p* < 0.001, ^✱✱✱✱^: *p* < 0.0001 (Unpaired t-test in C, G, I. Two-way ANOVA Uncorrected Fisher’s LSD in A-B).

**Supplementary Figure 4.**
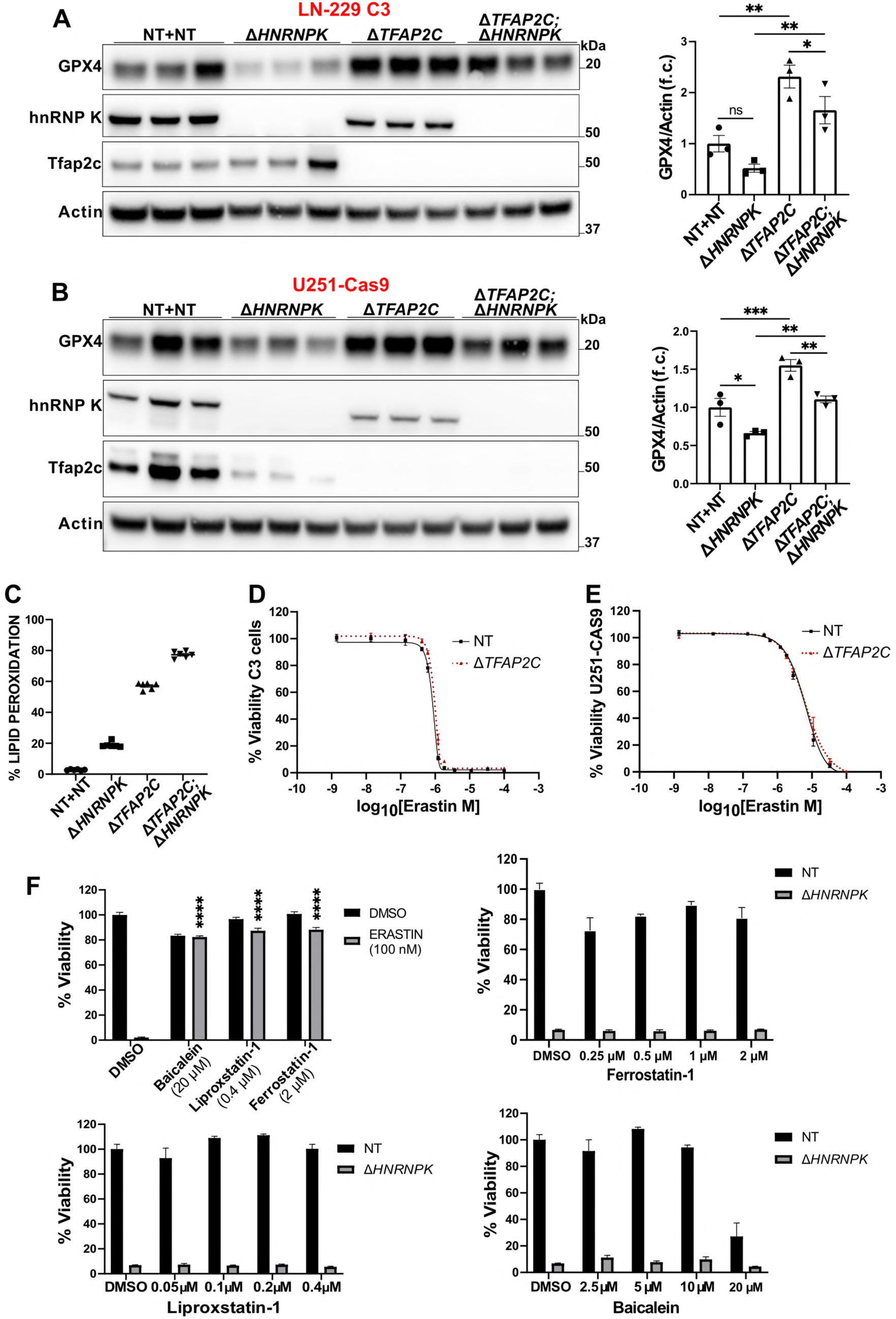
**A-B.** GPX4 protein after *HNRNPK* and *TFAP2C* ablation. *n* = 3. **C.** Percentage of Δ*TFAP2C* or NT-unmodified LN-229 C3 cells showing lipid peroxidation 4 days after delivering *HNRNPK* and NT qgRNAs (Liperfluo staining). *n* = 6. **D-E.** Viability of Δ*TFAP2C* treated with erastin or DMSO (CellTiter-Glo assay). Results are normalized against the DMSO-treated cells. *n* = 4. **F.** Viability of LN-229 C3 cells treated with erastin as a control (top left) or transduced with *HNRNPK* or NT qgRNAs and supplemented with anti-ferroptosis drugs: Ferrostatin-1 (top right), Liproxstatin-1 (bottom left), Baicalein (bottom right) (CellTiter-Glo assay). Results are normalized on the DMSO/NT condition. *n* ≥ 3. **Data information:** *n* represents independent cultures. f.c.: fold change. Mean ± SEM. ^✱^: *p* < 0.05, ^✱✱^: *p* < 0.01, ^✱✱✱^: *p* < 0.001, ^✱✱✱✱^: *p* < 0.001 (Two-way ANOVA Uncorrected Fisher’s LSD in A-B and Dunnett’s test in F).

**Supplementary Figure 5.**
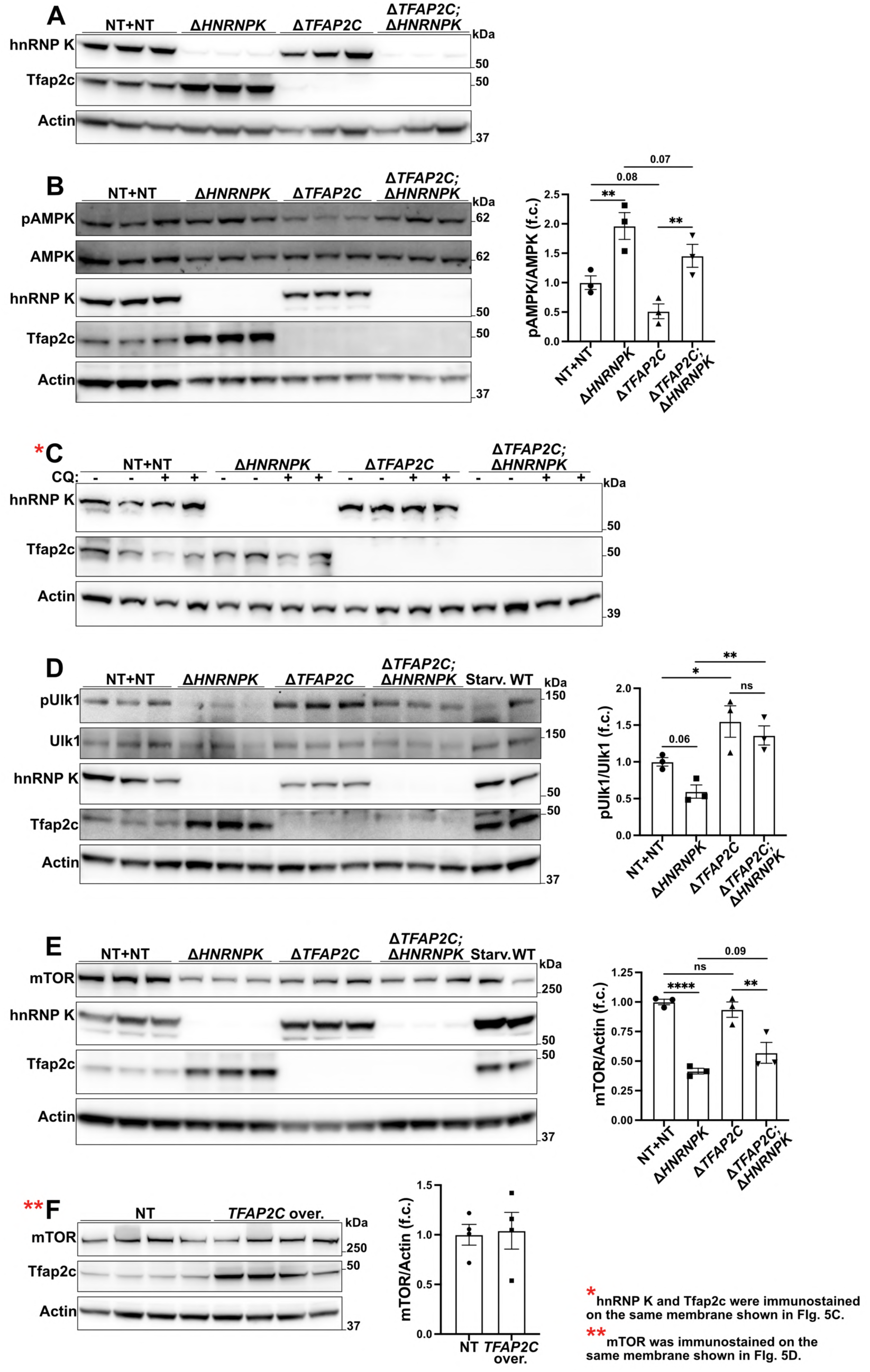
**A, C.** Ablations of *HNRNPK* and *TFAP2C* for RNA-seq samples (A) and from western blot shown in Fig. 5B (C), respectively. *hnRNP K and Tfap2c in C were immunostained on the same membrane shown in Fig. 5B. **B, D-E.** mTOR protein (E) and AMPK (B) and Ulk1 (D) phosphorylation ratio upon deletion of *HNRNPK* and *TFAP2C* in LN-229 C3 cells. 6h HBSS-starvation (starv.) was used as a positive control *n* = 3. **F.** mTOR protein upon *TFAP2C* overexpression in LN-229 dCas9-VPR cells. **mTOR was immunostained on the same membrane shown in Fig. 5D. **Data information:** *n* represents independent cultures. f.c.: fold change. Mean ± SEM. ^✱^: *p* < 0.05, ^✱✱^: *p* < 0.01, ^✱✱✱✱^: *p* < 0.001 (Two-way ANOVA Uncorrected Fisher’s LSD).

**Supplementary Figure 6.**
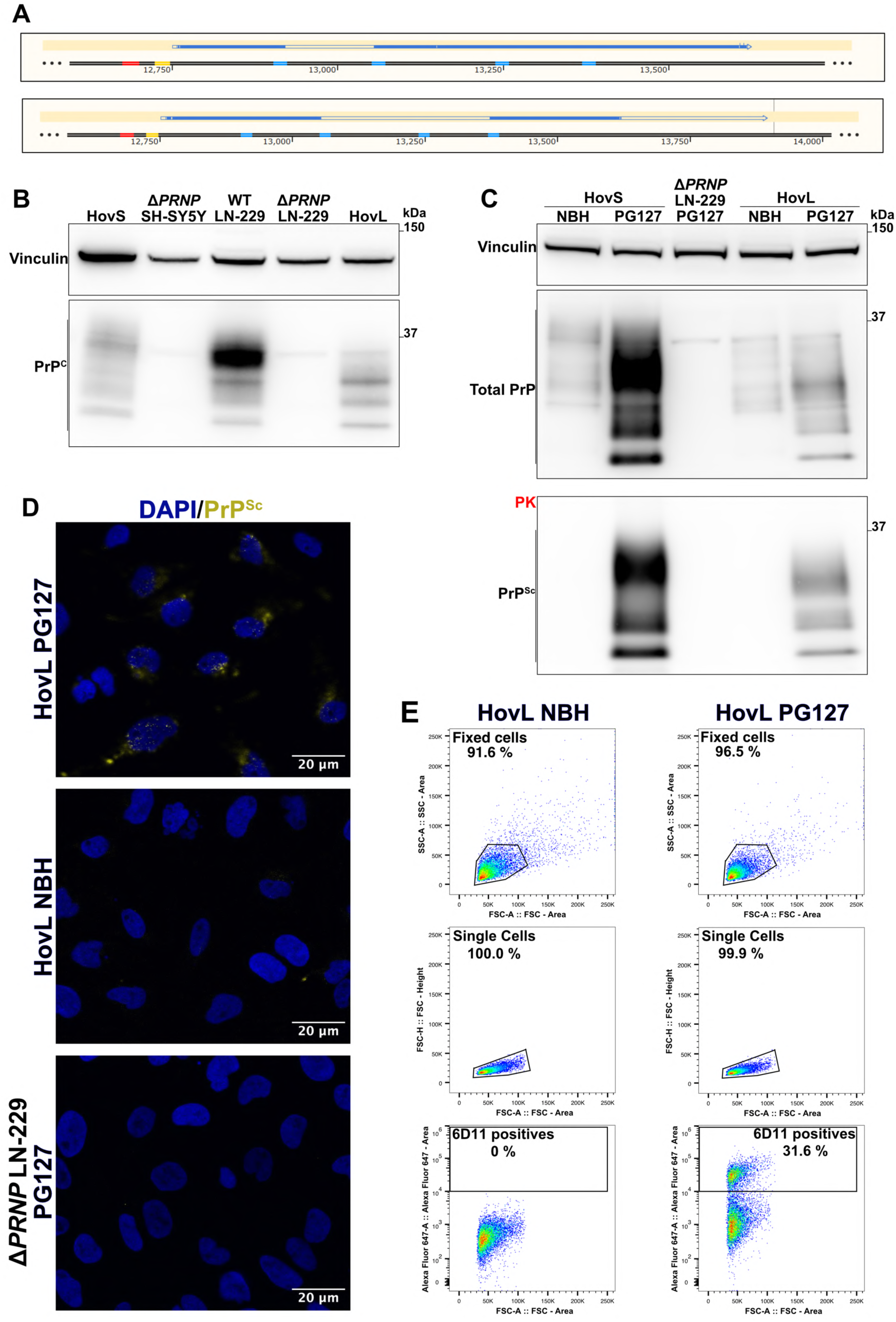
**A.** qgRNAs (blue segments in reference sequence) and Cas9 transiently transfected in LN-229 cells promoted two major *PRNP* deletions from position 12922 to 13055 and from 13055 to 13372. **B.** Western blot showing the lack of the human PrP^C^ protein in the LN-229^Δ*PRNP*^ from A and the expression of the ovine PrP^C^ in the resulting HovL cells. HovS and SH-SY5Y^Δ*PRNP*^ cells were used as controls for the “ovinization” and *PRNP* ablation, respectively. **C.** Proteinase K (PK) digested (bottom) and undigested (top) western blots showing respectively PrP^Sc^ and the total PrP in LN-229^Δ*PRNP*^ and HovL cells inoculated either with PG127 prion-infected Brain Homogenate (PG127) or with Not-infectious Brain Homogenate (NBH). PG127-infected and NBH mock-infected HovS cells were used as positive and negative controls, respectively. **D-E.** Imaging (D) and flow cytometry analysis (E) of anti-PrP^Sc^ 6D11 antibody signal in PG127-infected HovL or LN-229^Δ*PRNP*^ and in NBH HovL cells.

**Supplementary Figure 7.**
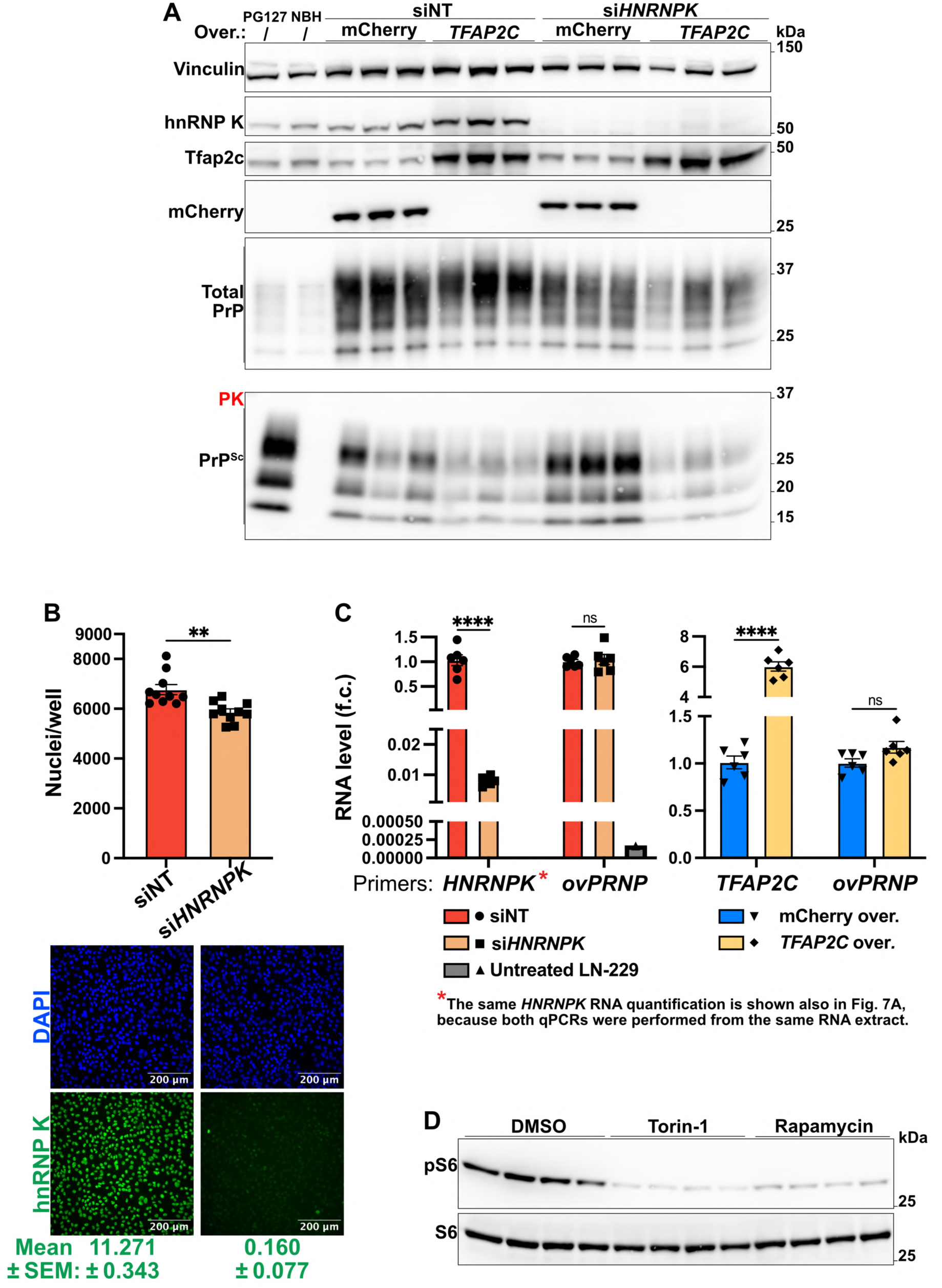
**A.** Proteinase K (PK) digested (bottom) and undigested (top) western blots showing, respectively, PrP^Sc^ and total PrP after *HNRNPK* silencing (96 hours) and *TFAP2C* overexpression (192 hours) in PG127-infected HovL cells. *n* = 3. **B.** Cell density (nuclei per well) (top) after *HNRNPK* silencing (96 hours) in PG127-infected HovL cells. Representative image and quantification of hnRNP K intensity per cell (mean ± SEM) (bottom). *n* = 10. **C.** qRT-PCR showing *HNRNPK* and *ovPRNP* RNA upon *HNRNPK* silencing (96 hours) (left), and *TFAP2C* and *ovPRNP* RNA upon *TFAP2C* overexpression (192 hours) (right) in PG127-infected HovL cells. LN-229 cells were used as a negative control for *ovPRNP* expression. *n* = 6. *The same RNA extract was used for the qPCR shown in Fig. 7A. **D.** S6 protein phosphorylation in PG127-infected HovL cells treated with 500 nM of Torin-1 or Rapamycin (72 hours) (Same samples used for Fig. 7C). **Data information:** Non-targeting siRNA (siNT) and mCherry overexpression were used as controls. qRT-PCR results are normalized against *GAPDH* expression. *n* represents independent cultures. f.c.: fold change. Mean ± SEM. ns: *p* > 0.05, ^✱✱^: *p* < 0.01, ^✱✱✱✱^: *p* < 0.001 (Unpaired t-test in B. Multiple Unpaired t-test Holm-Šídák method in C).

## Uncropped western blots

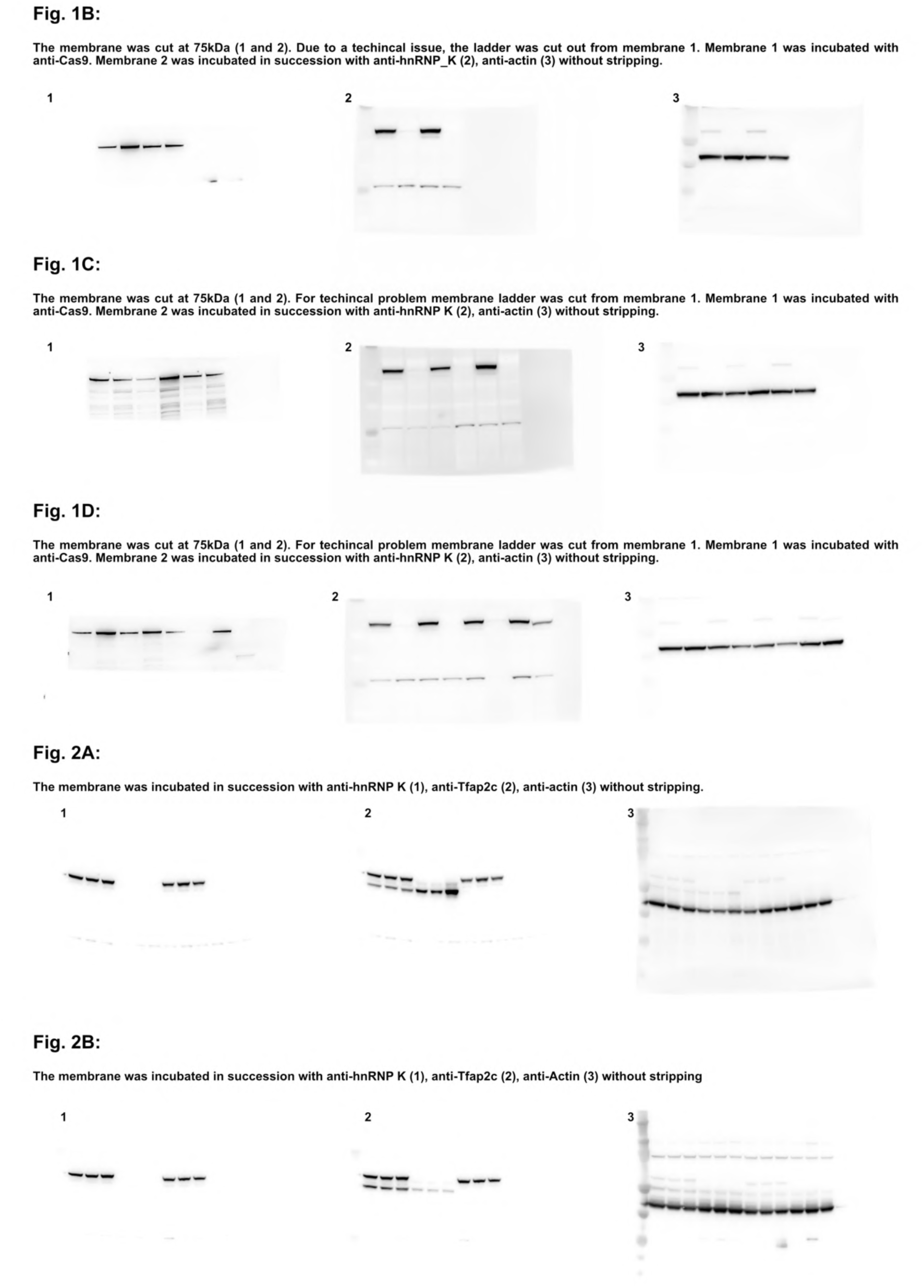
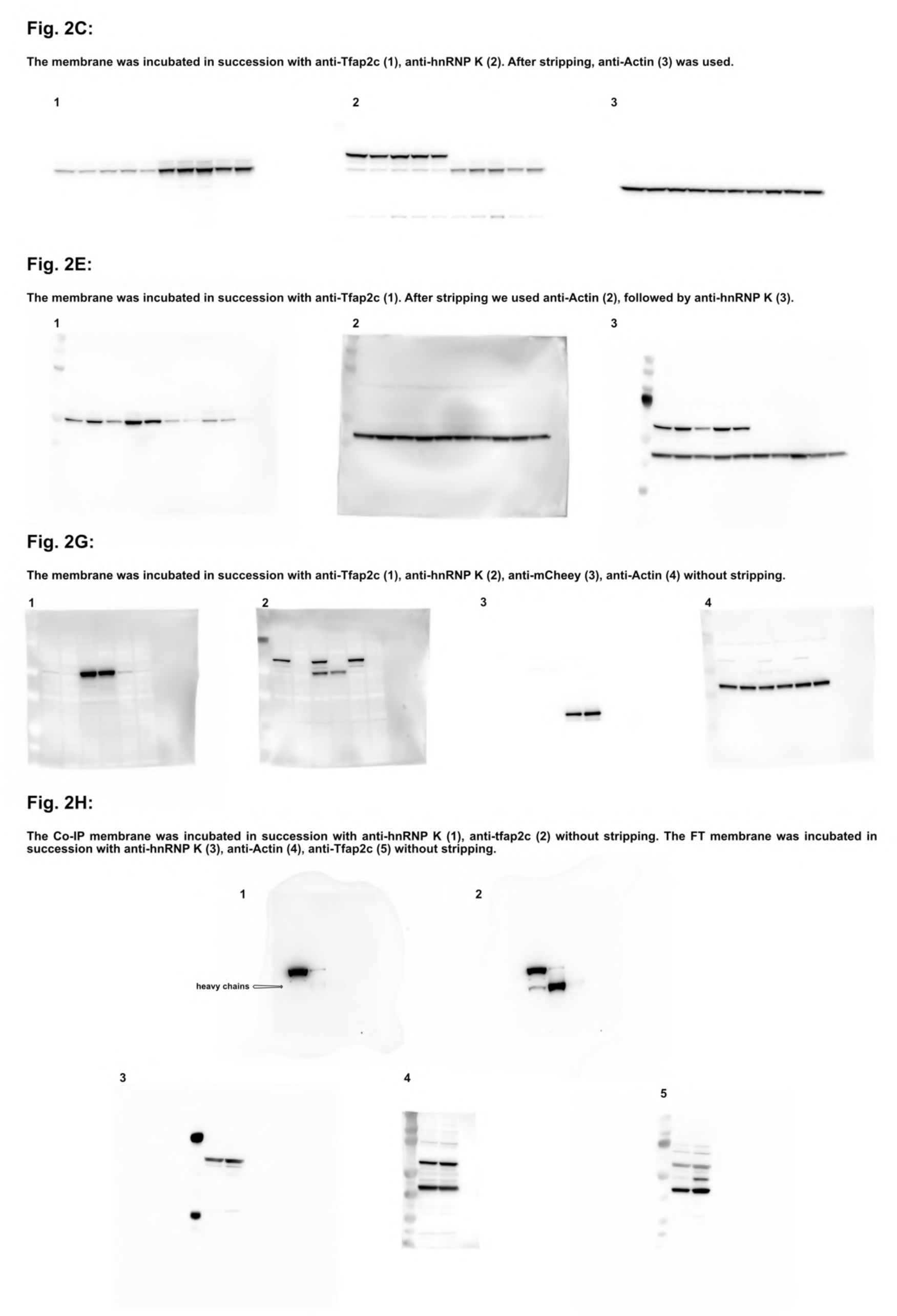
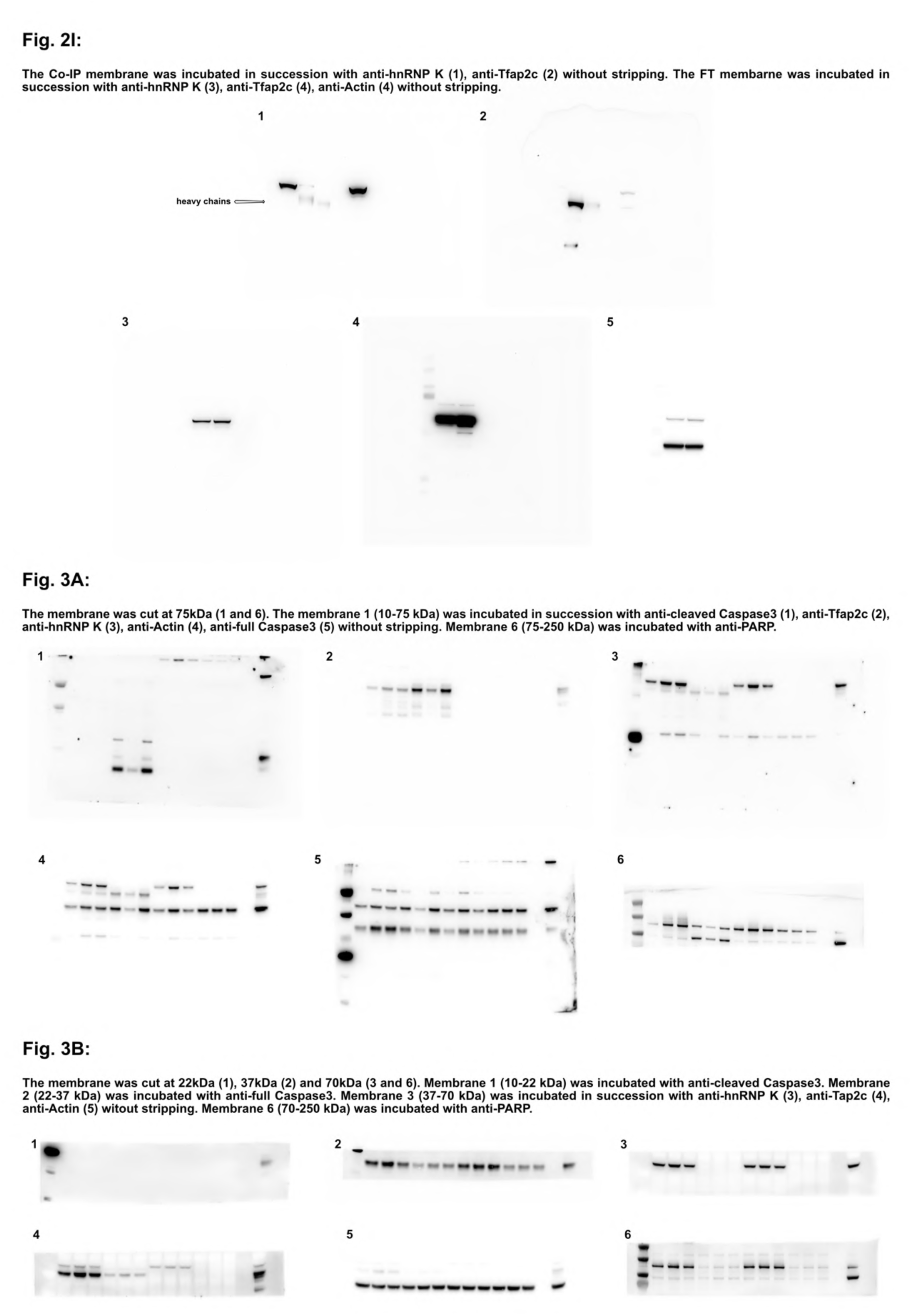
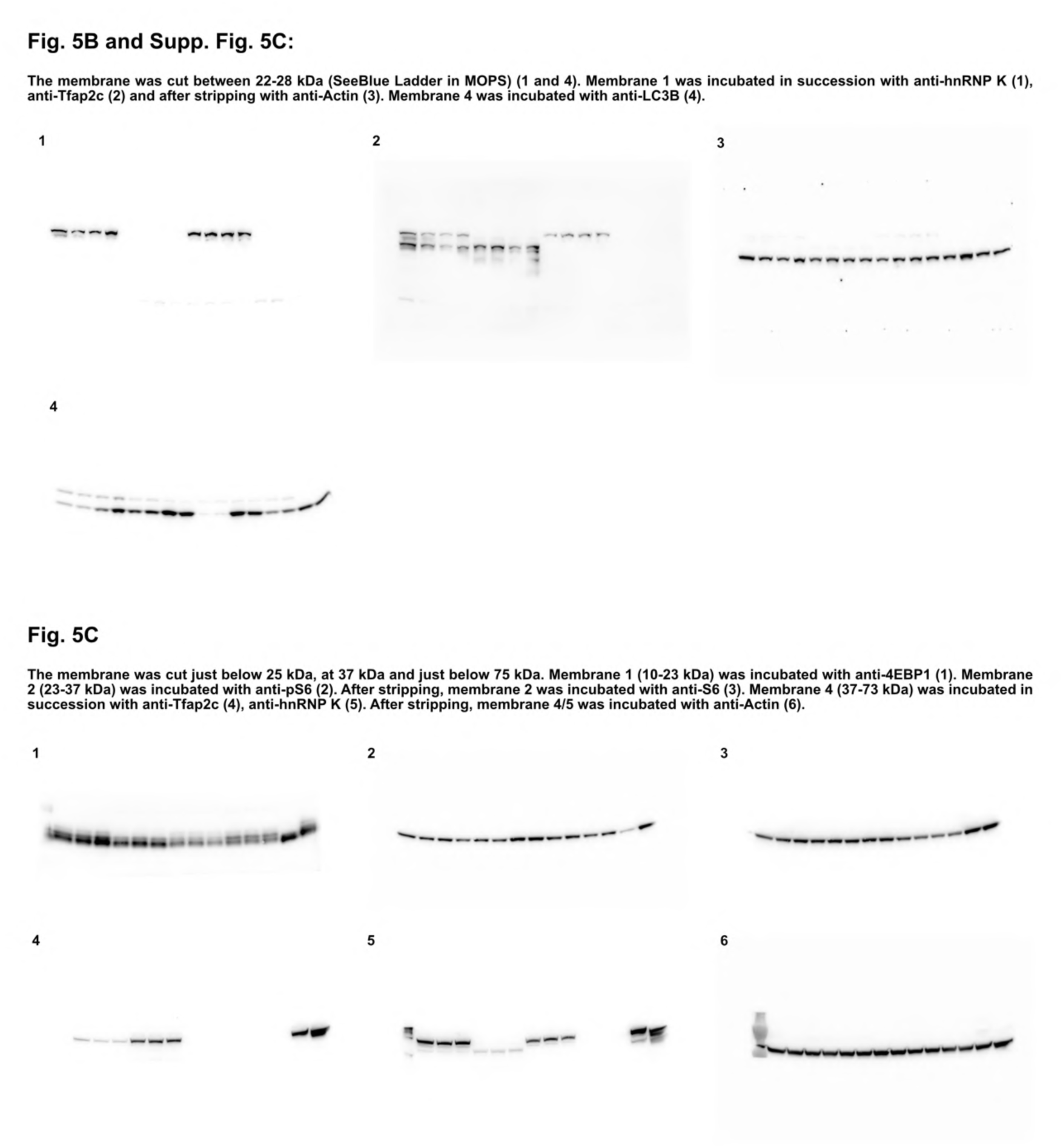
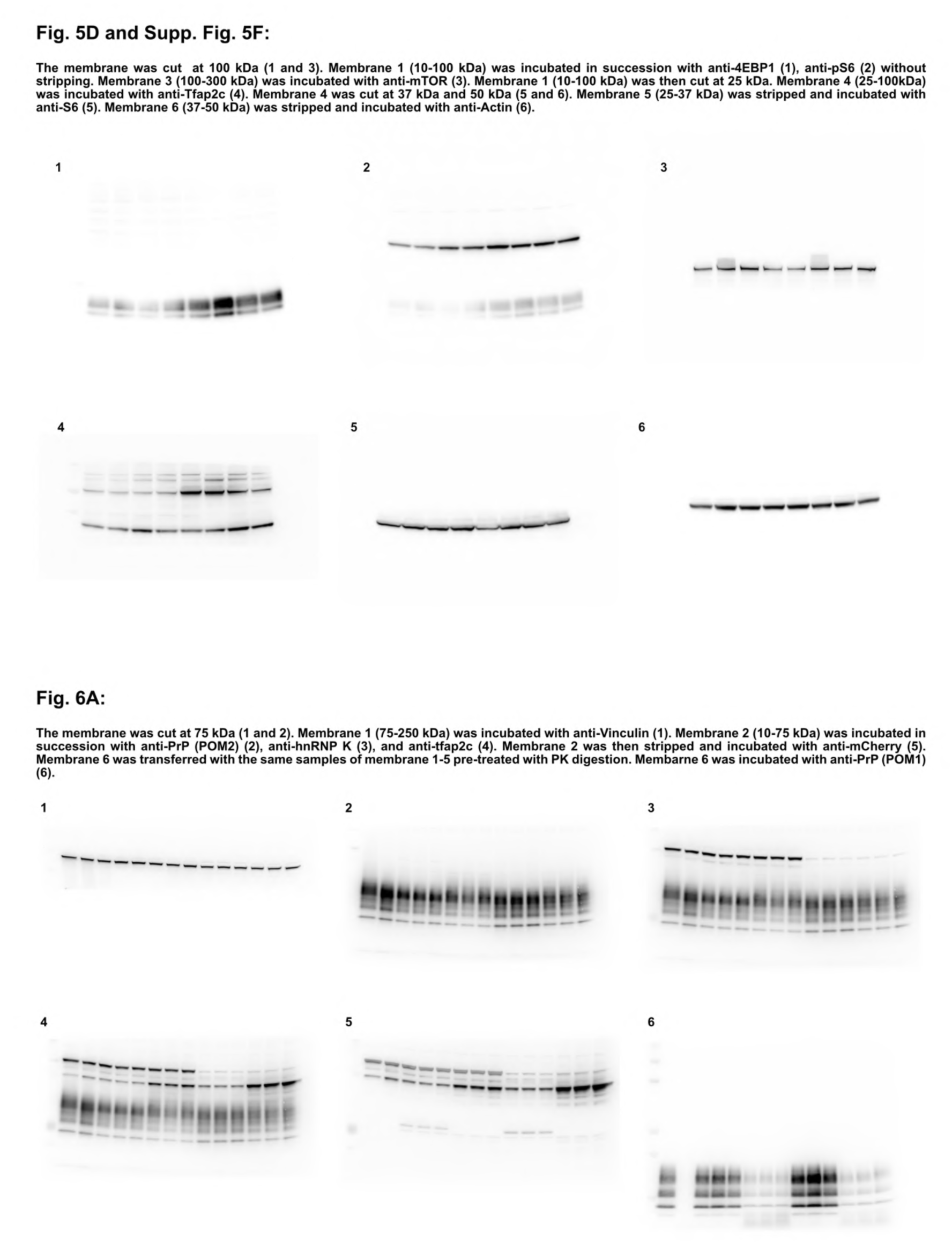
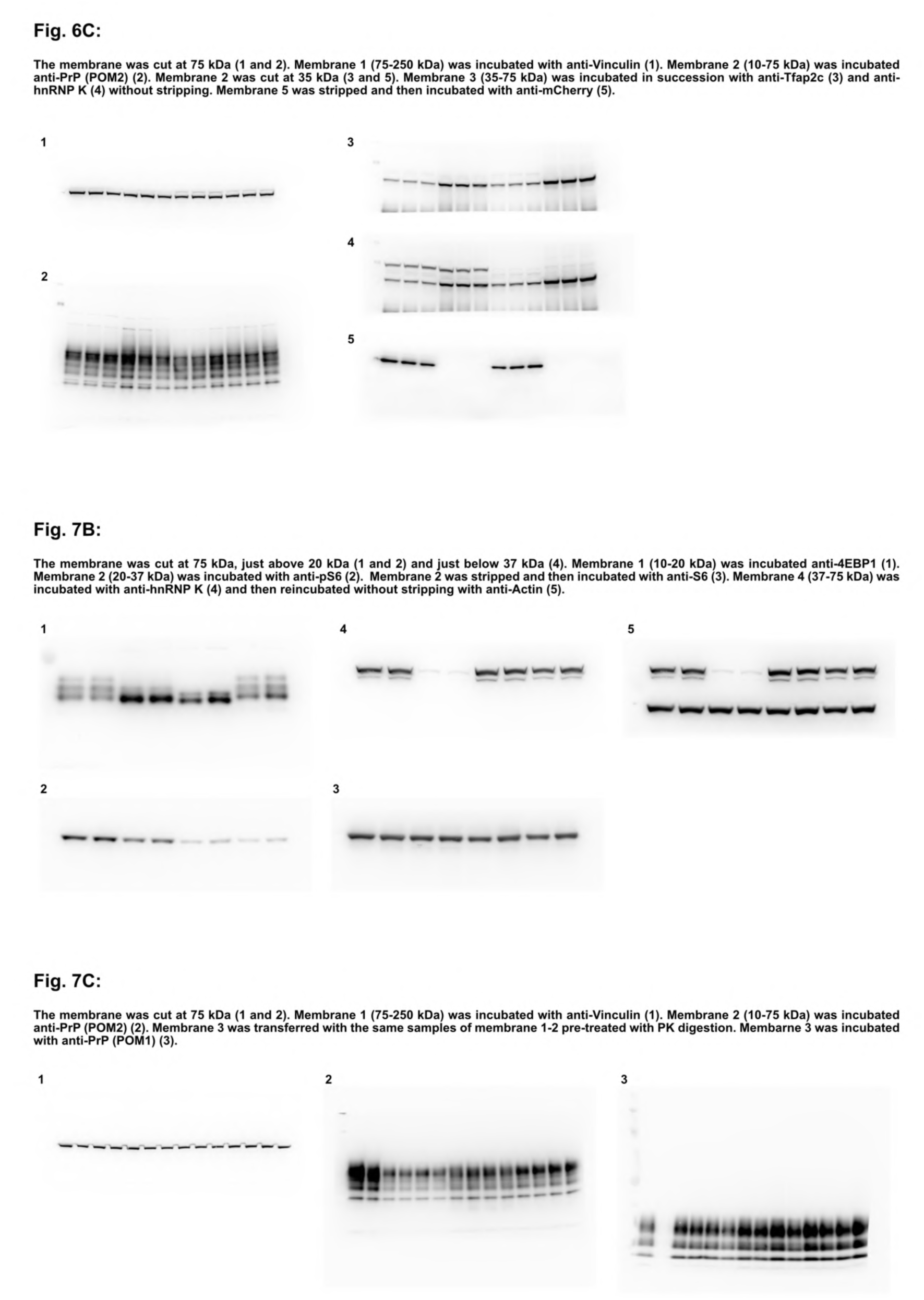
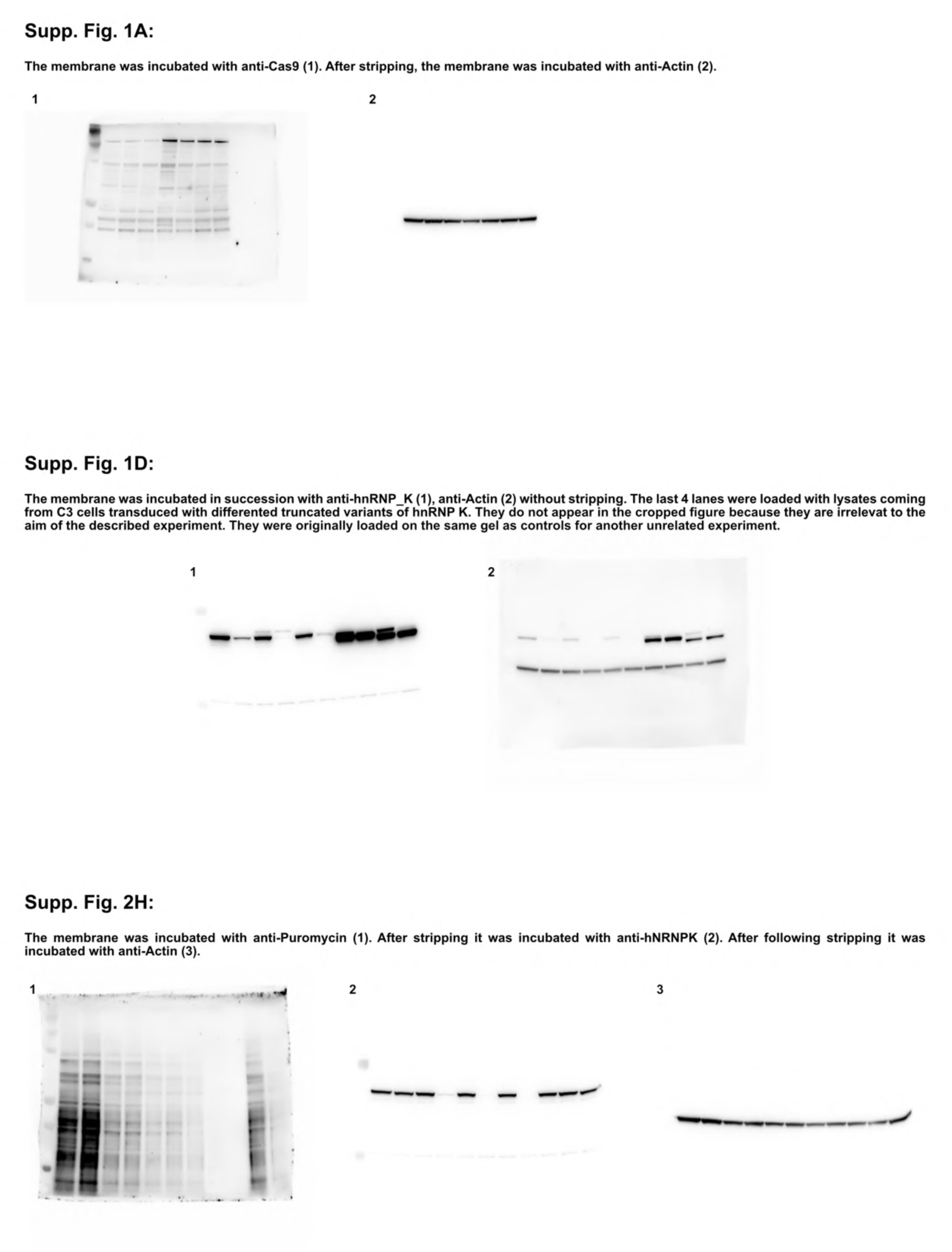
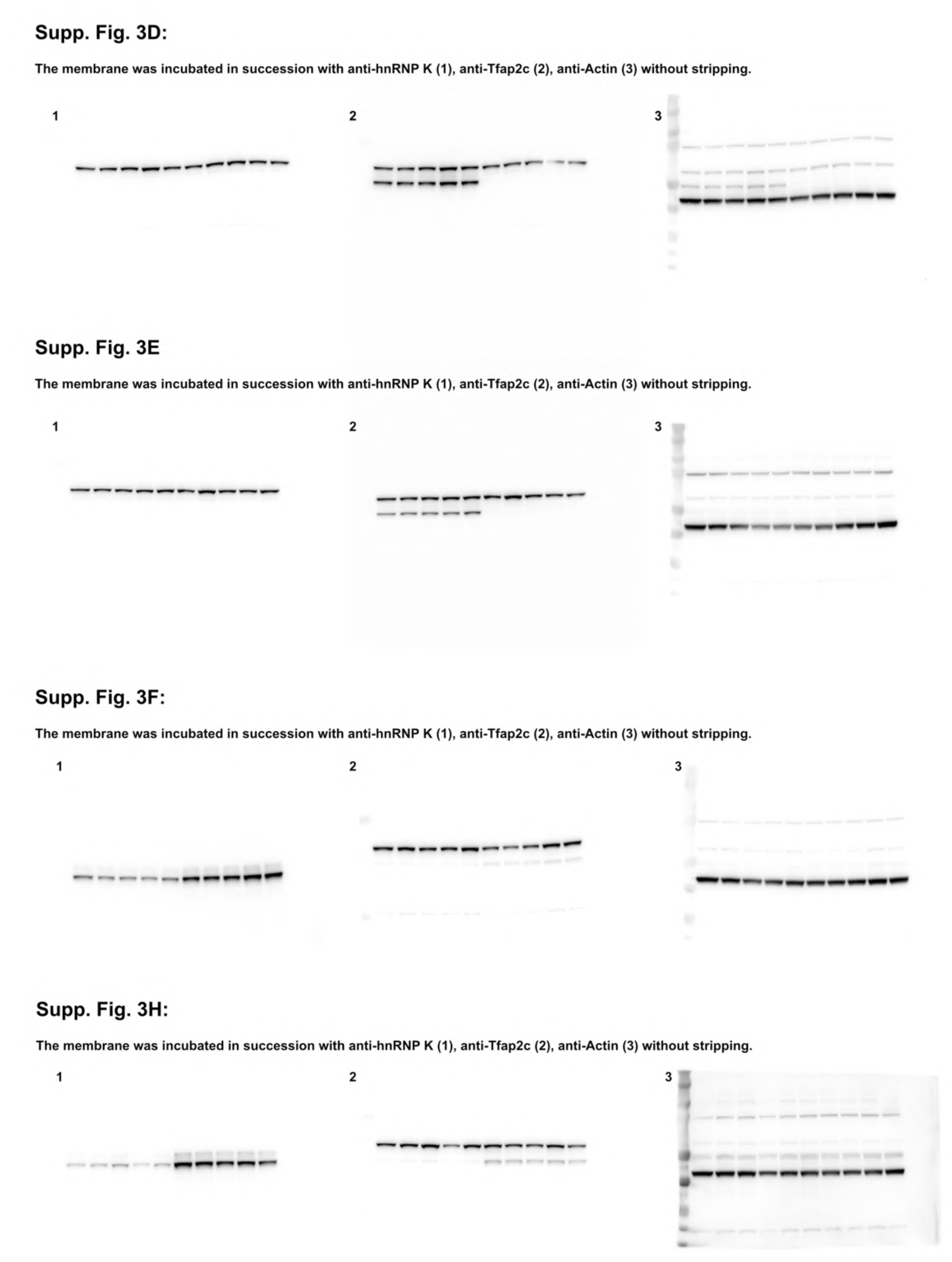
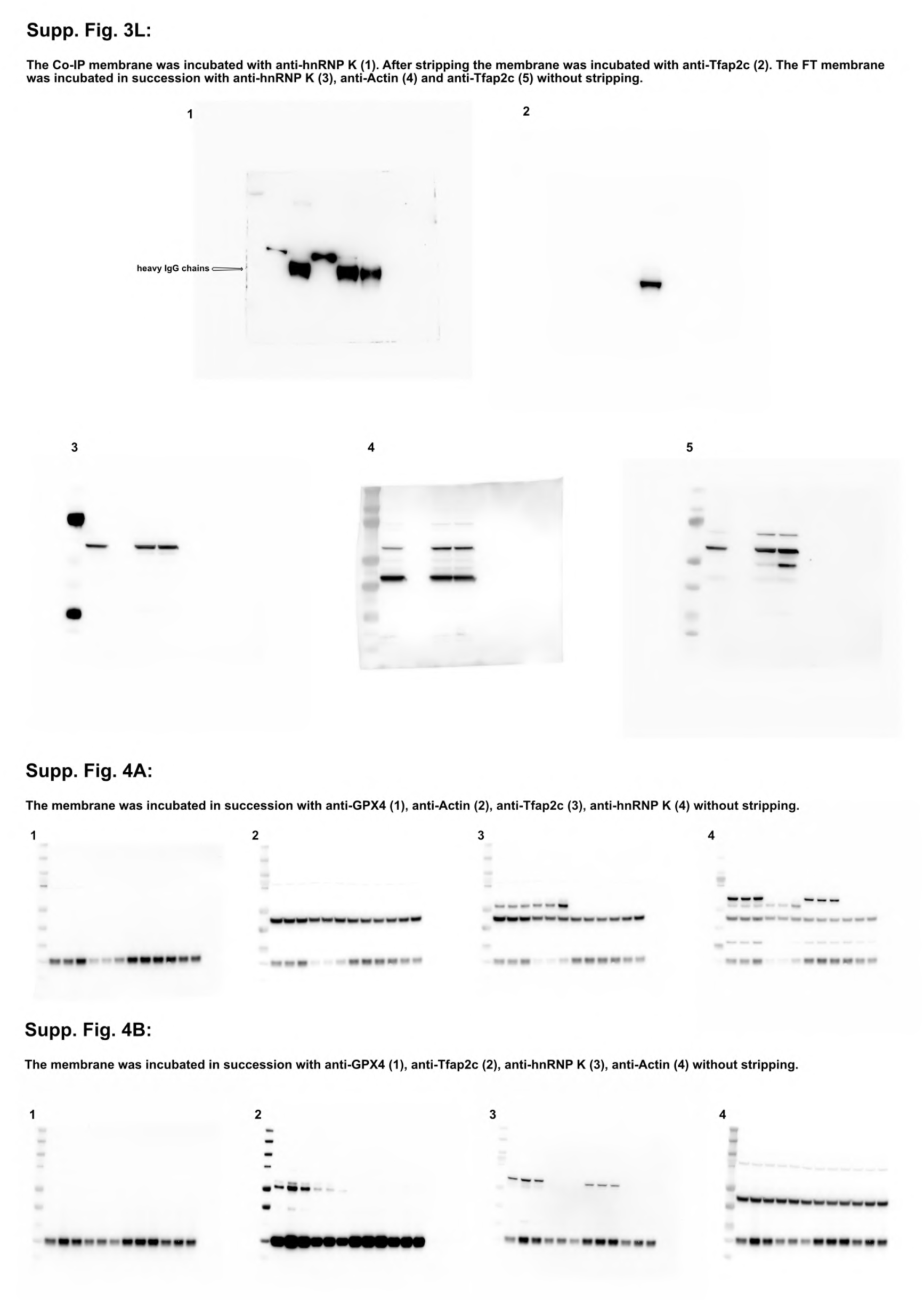
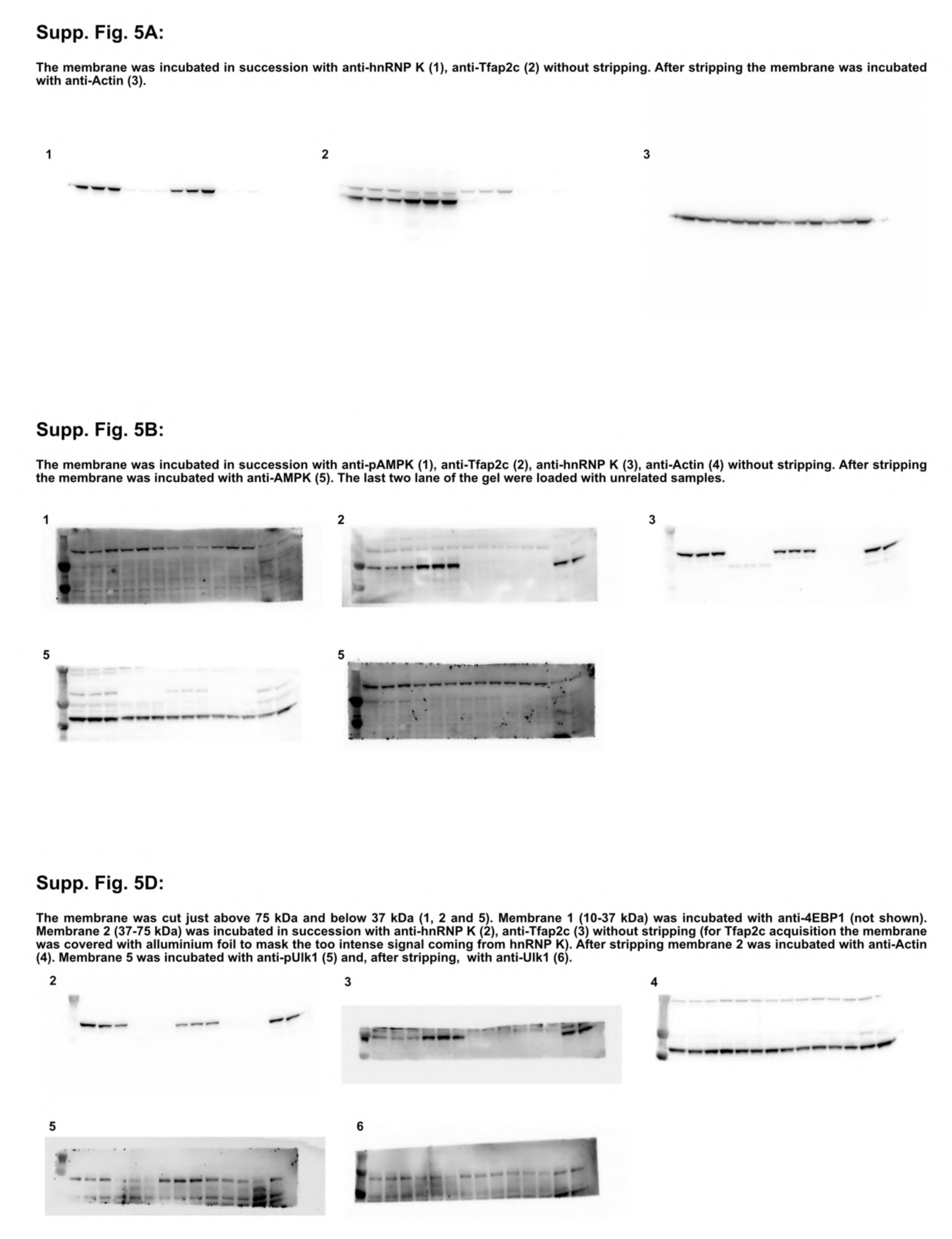
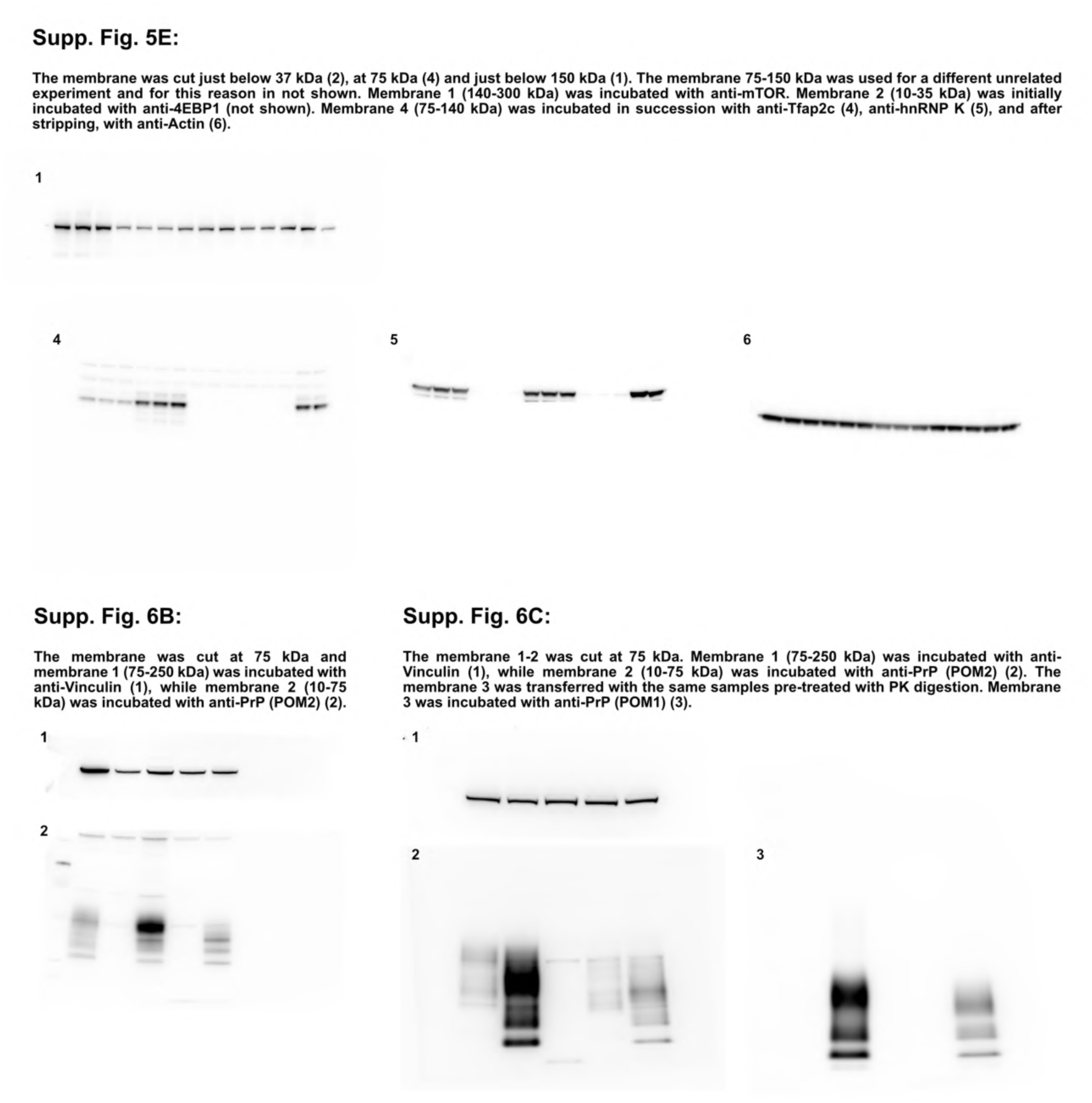
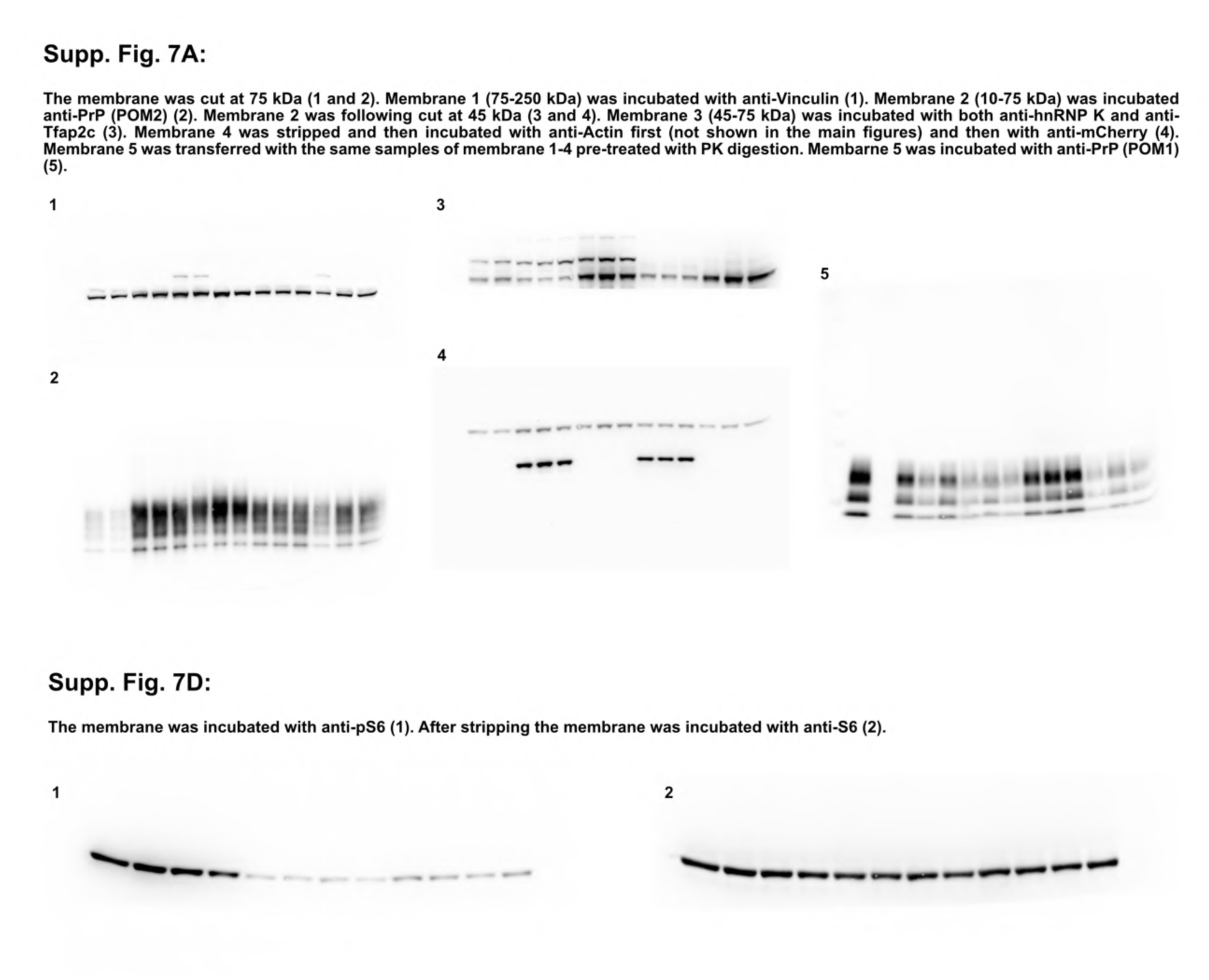

## Gating strategy

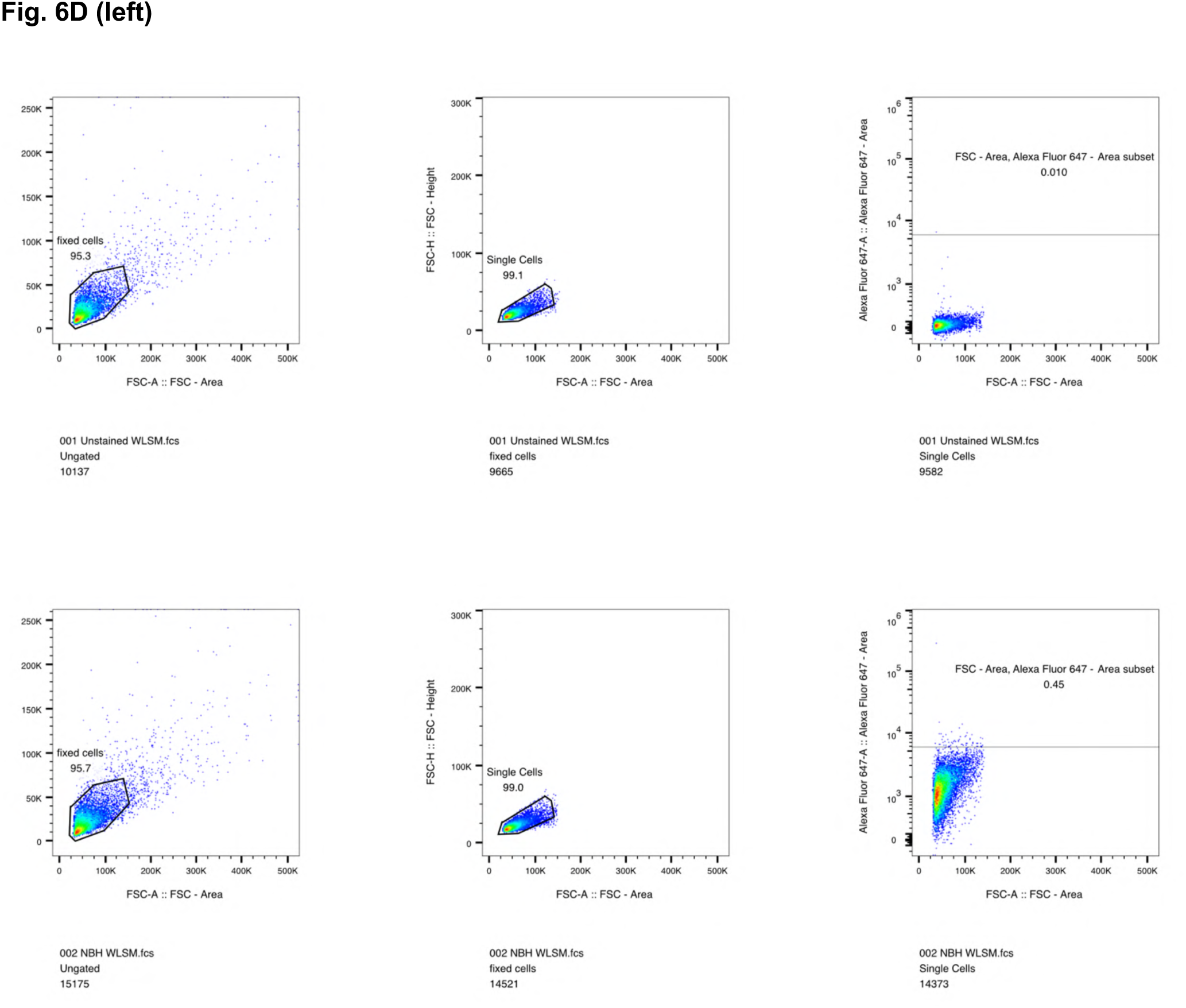
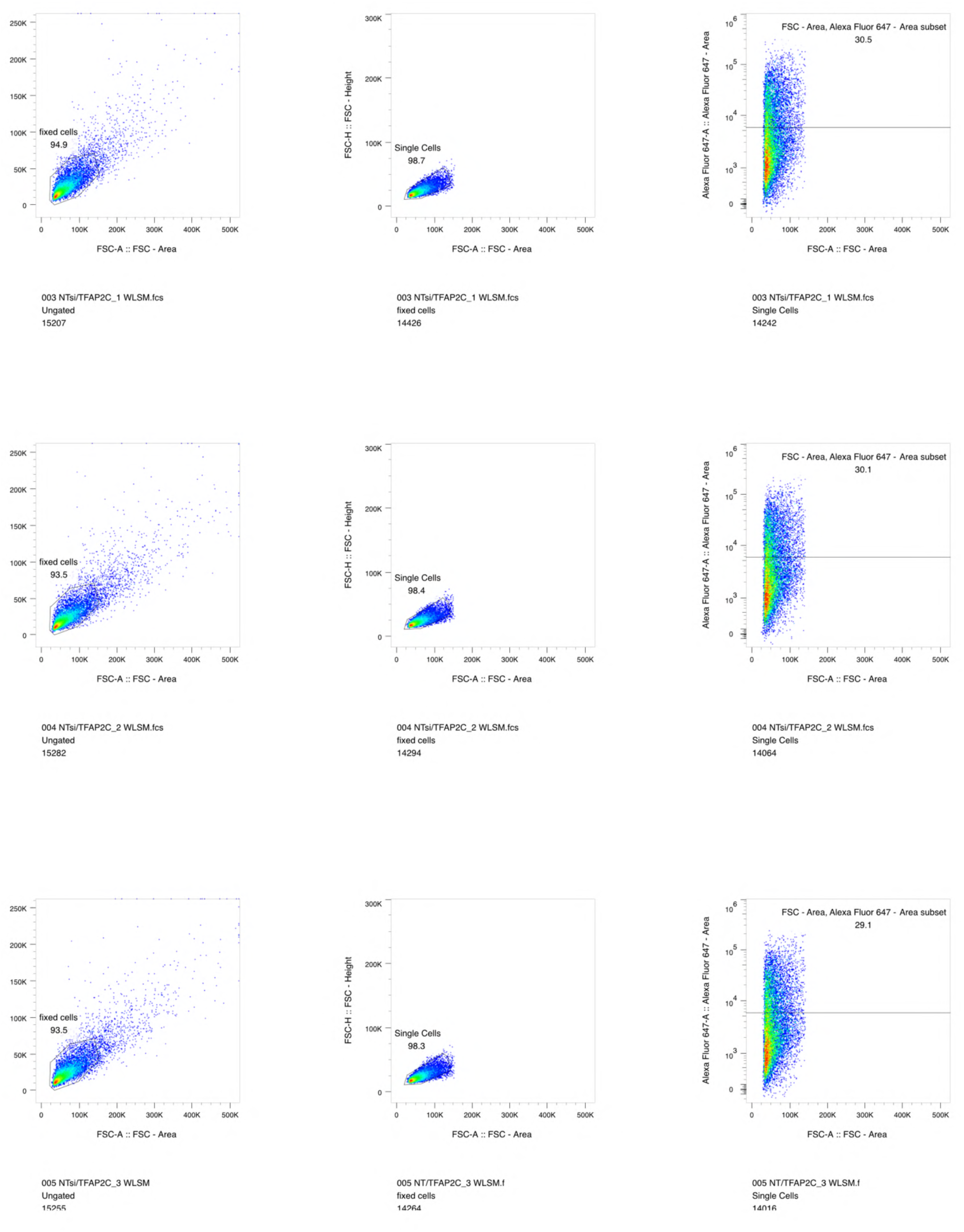
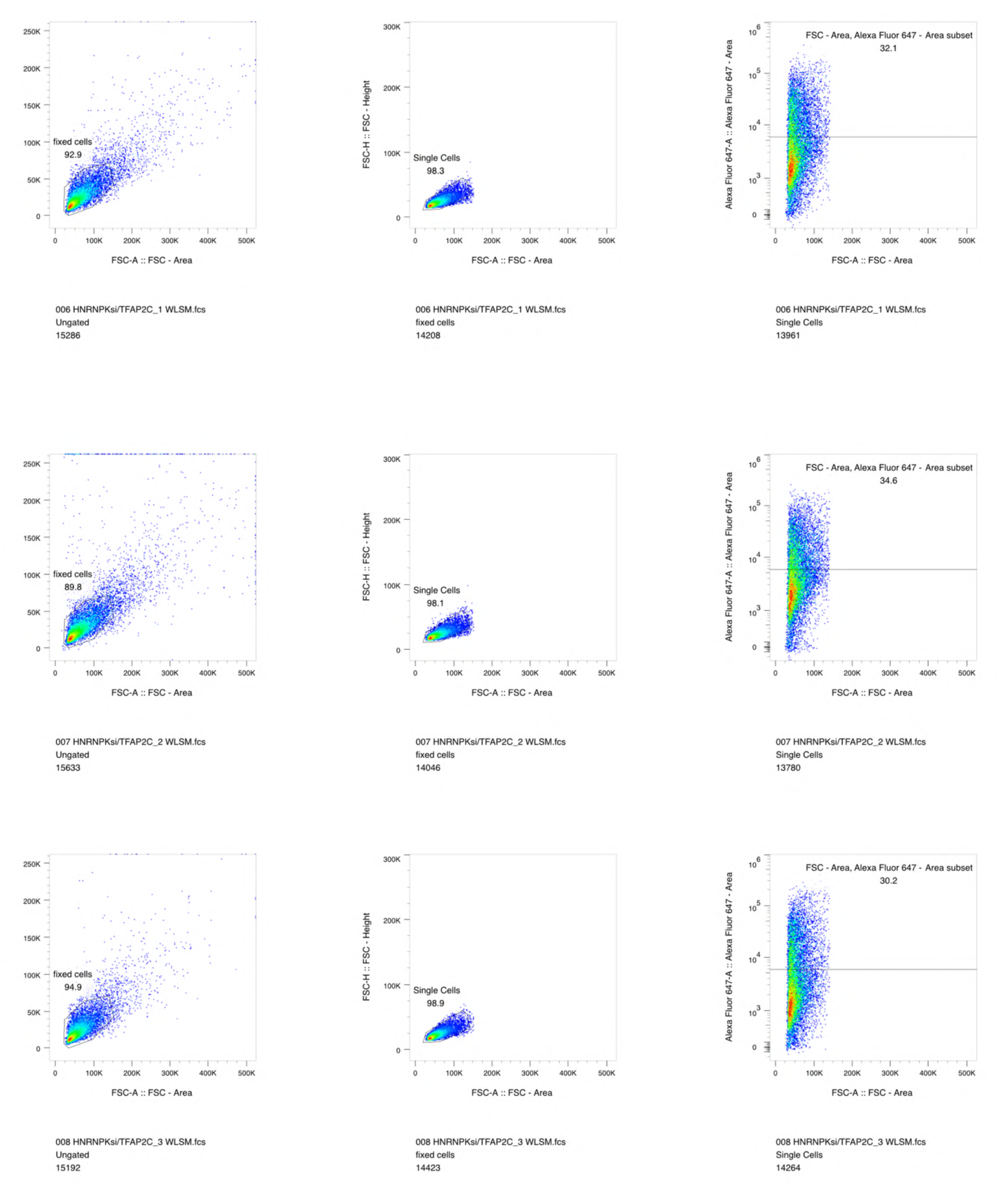
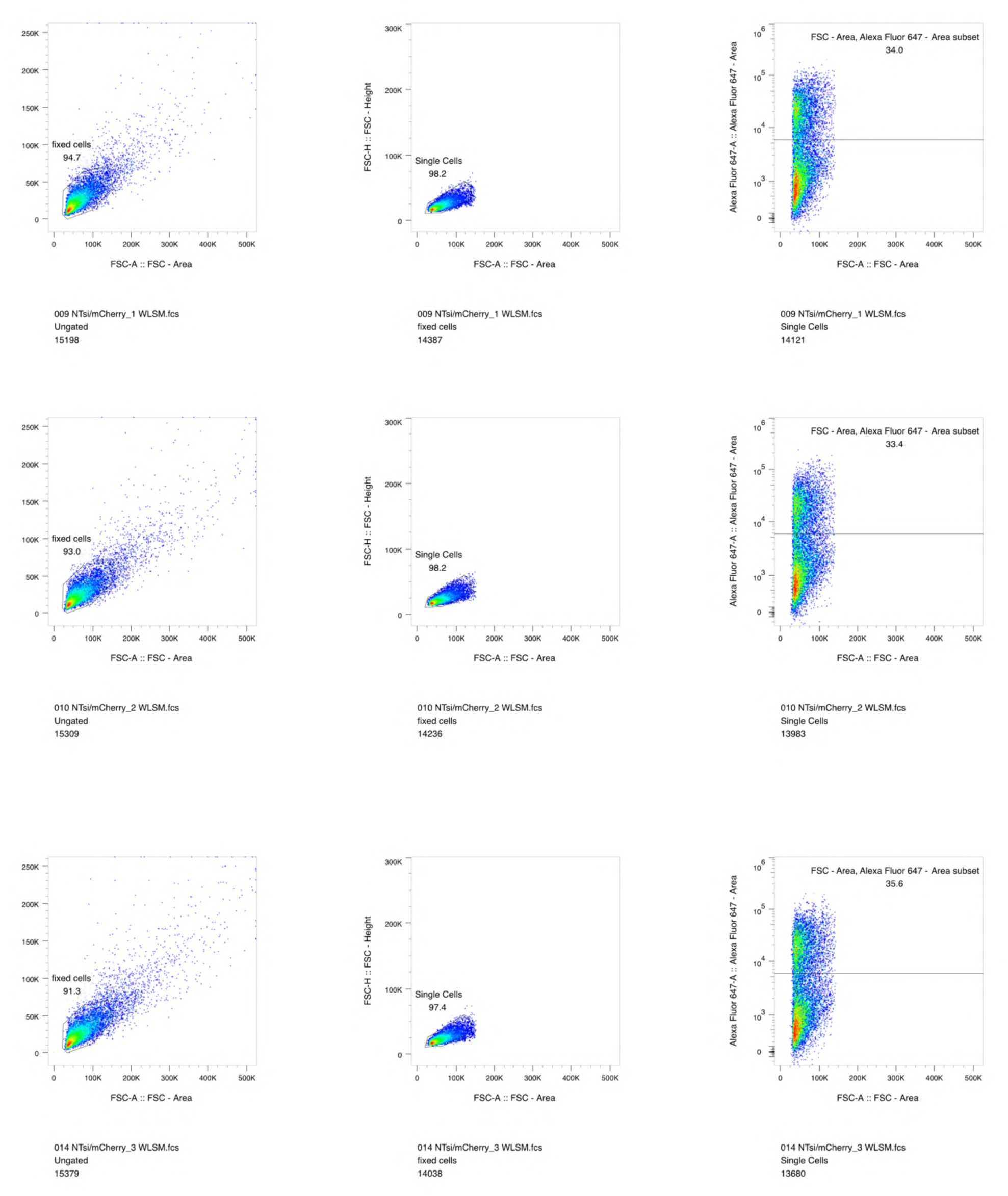
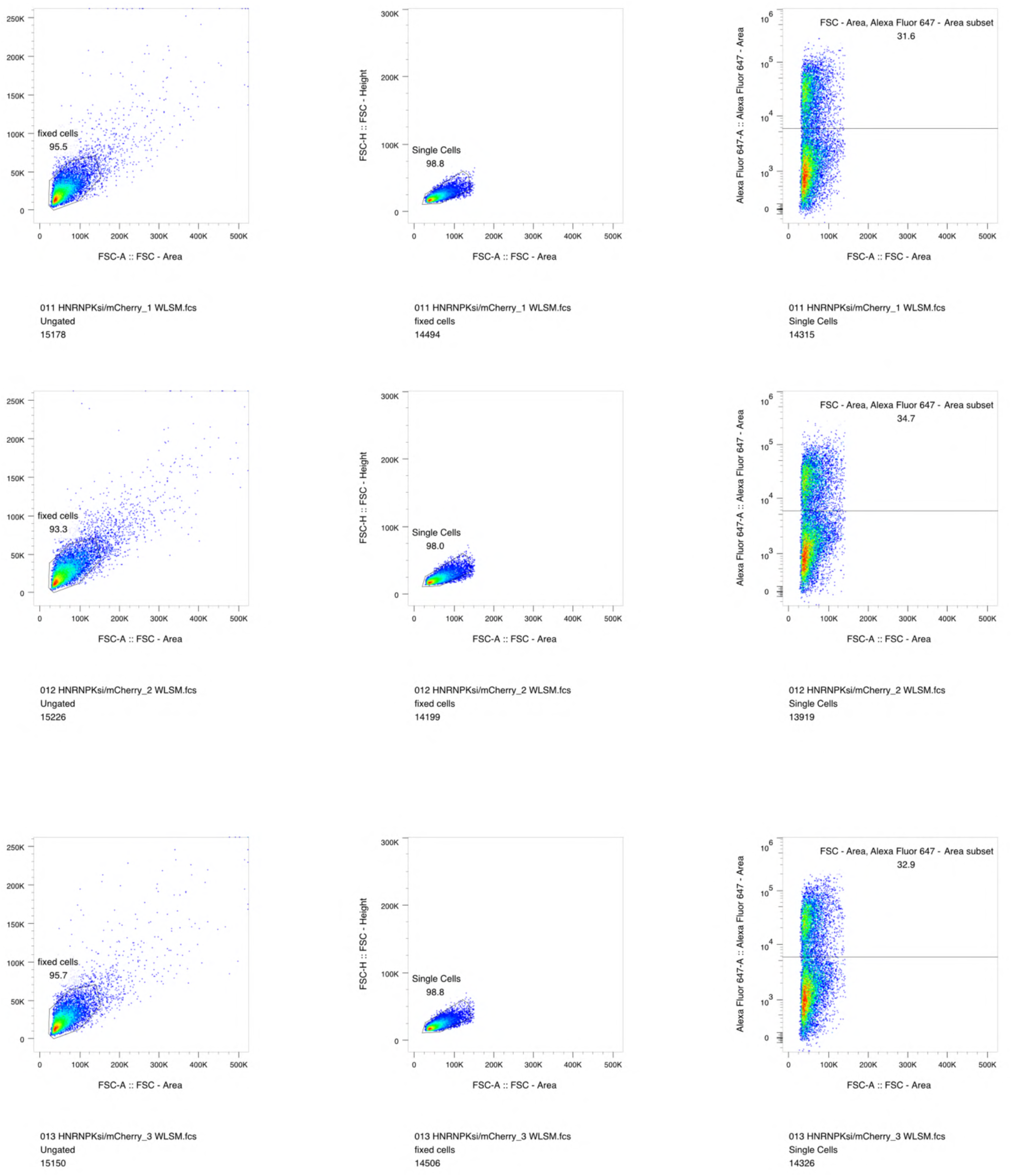
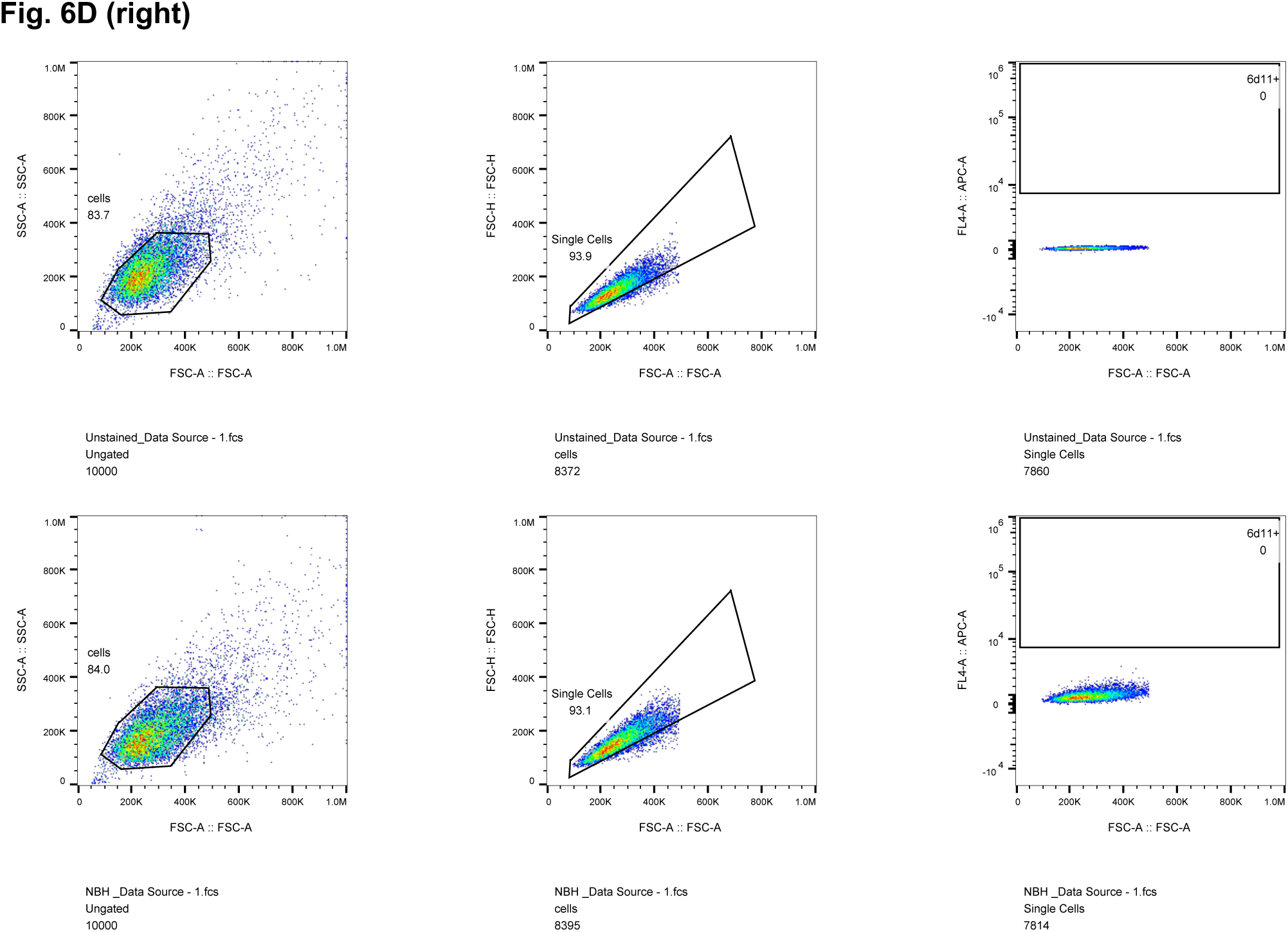
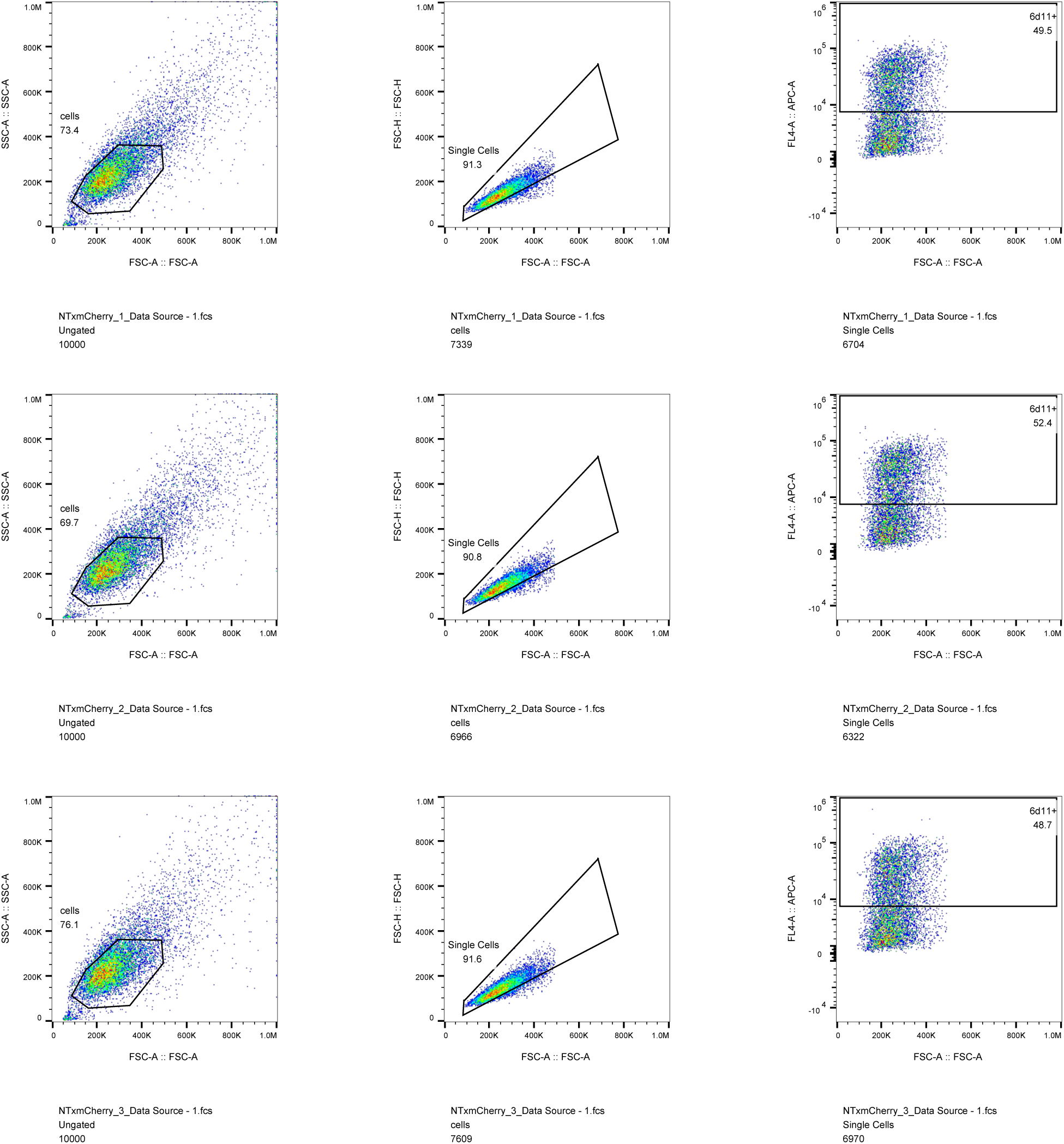
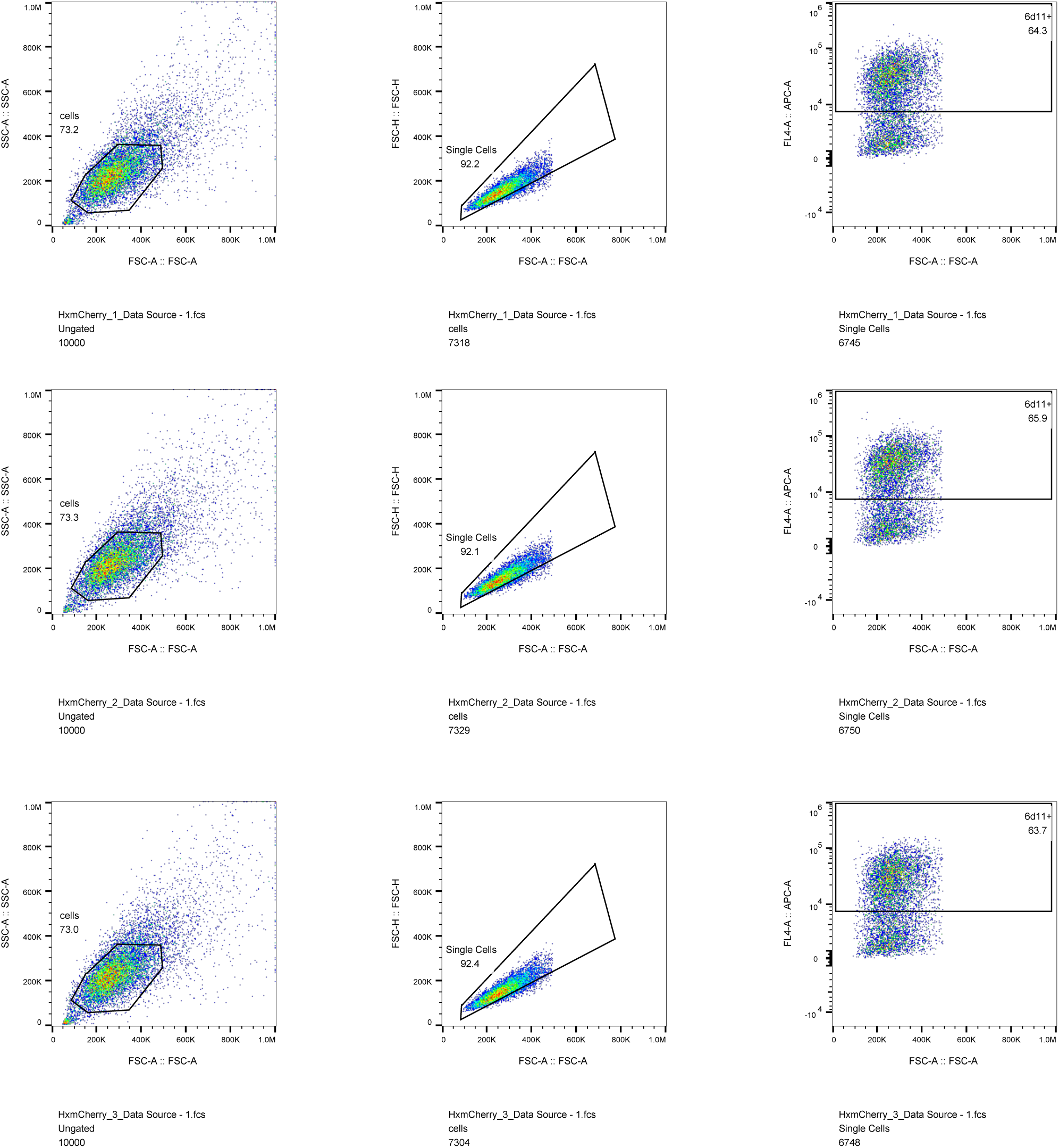
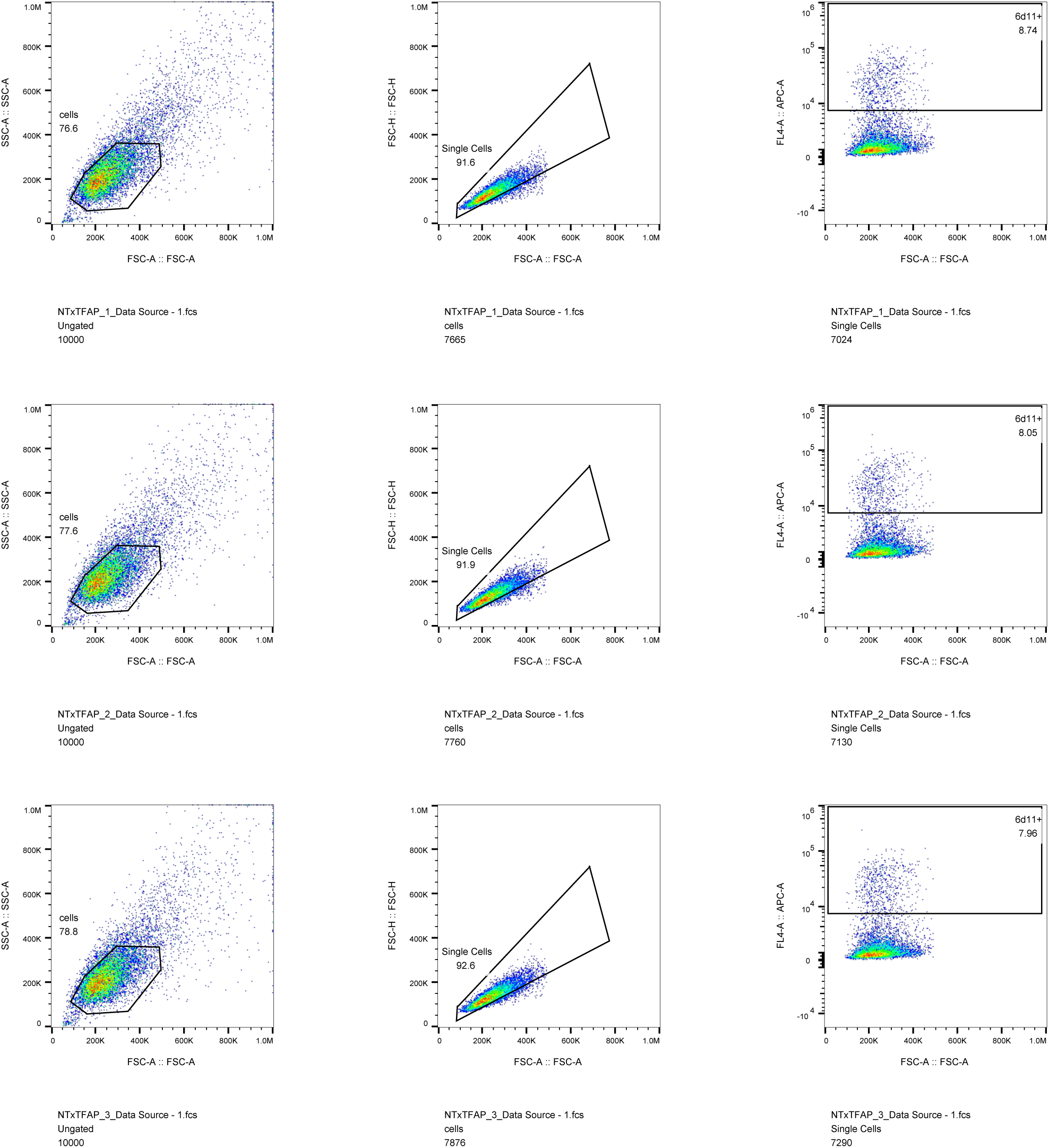
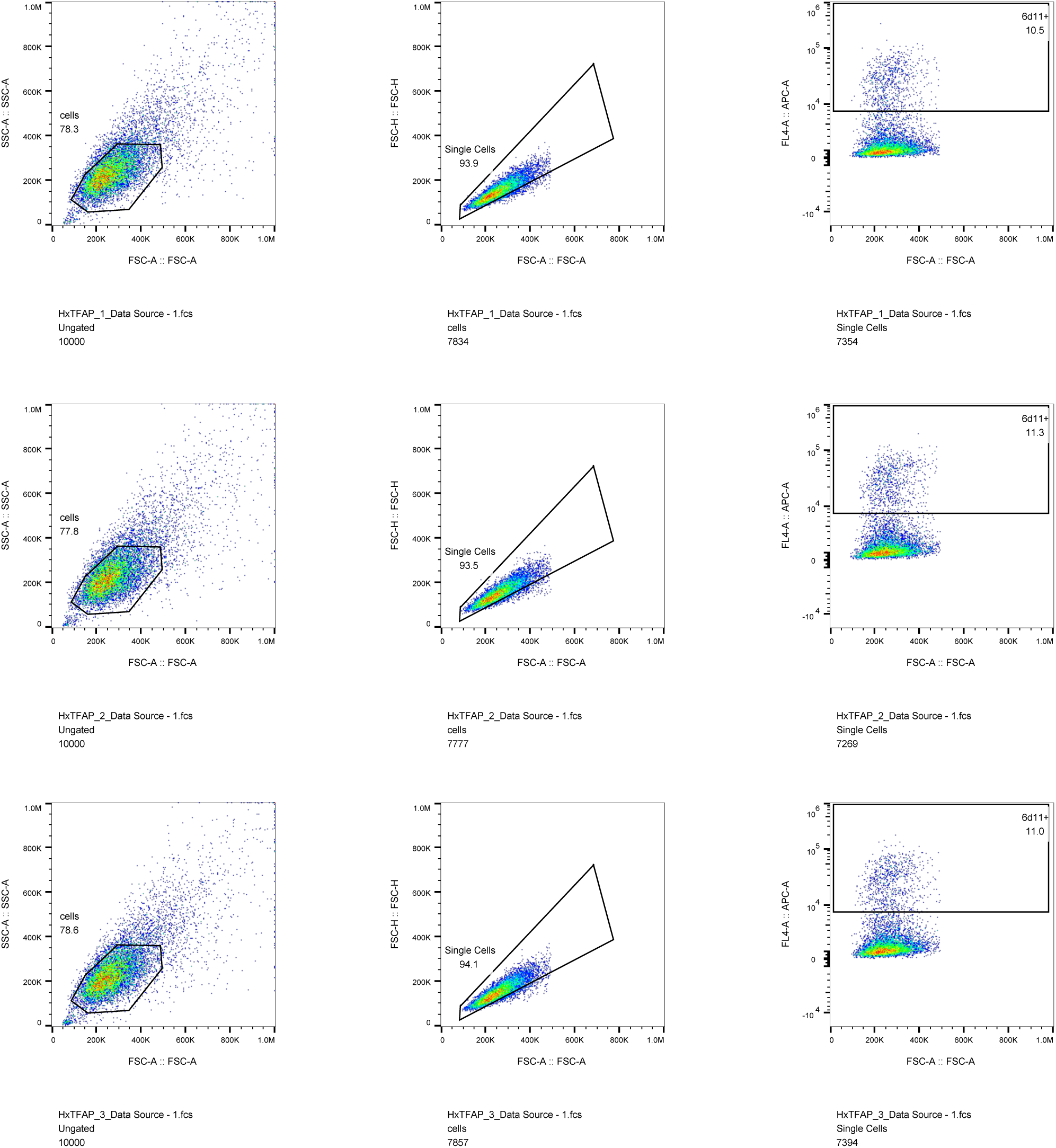
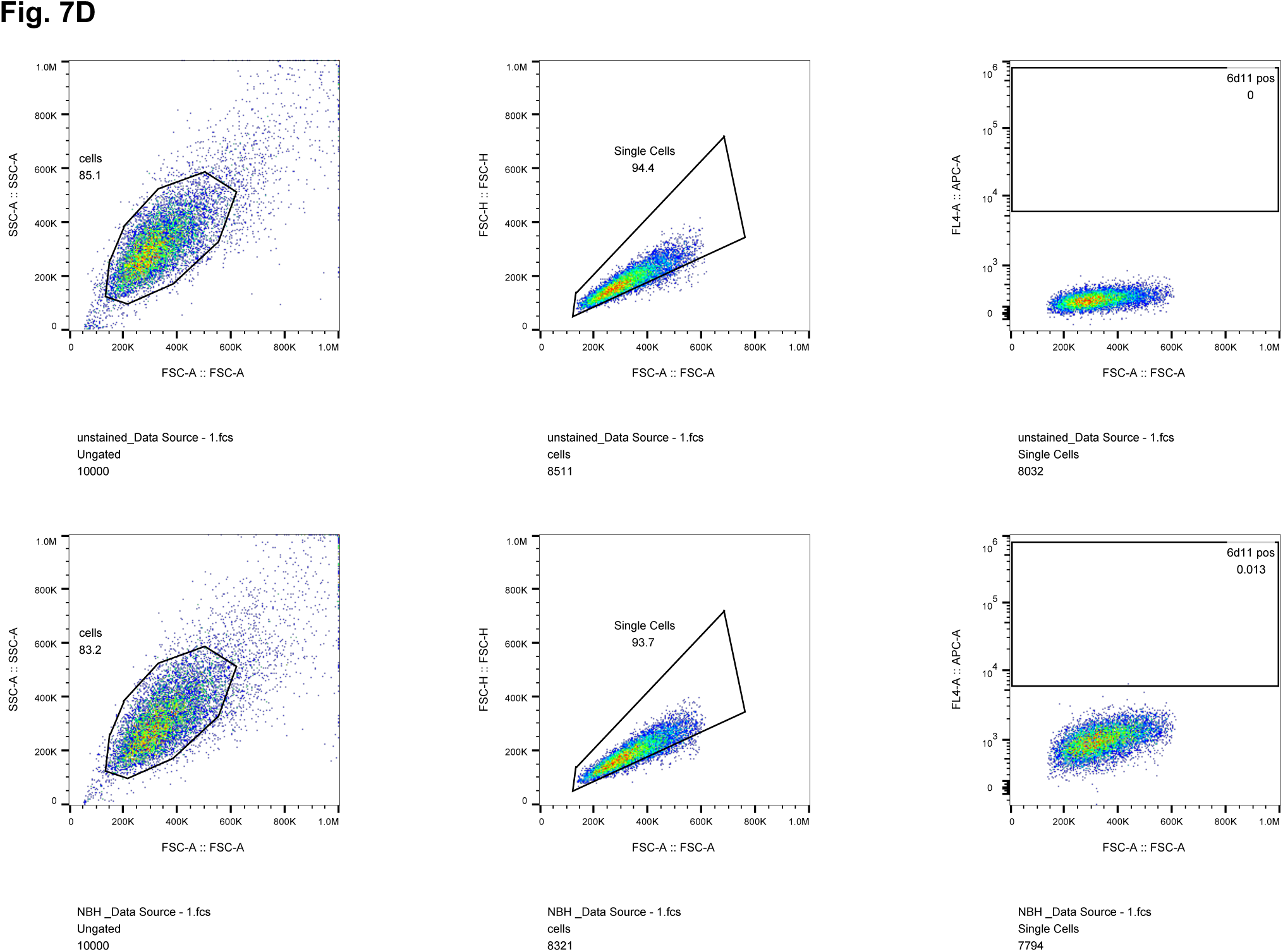
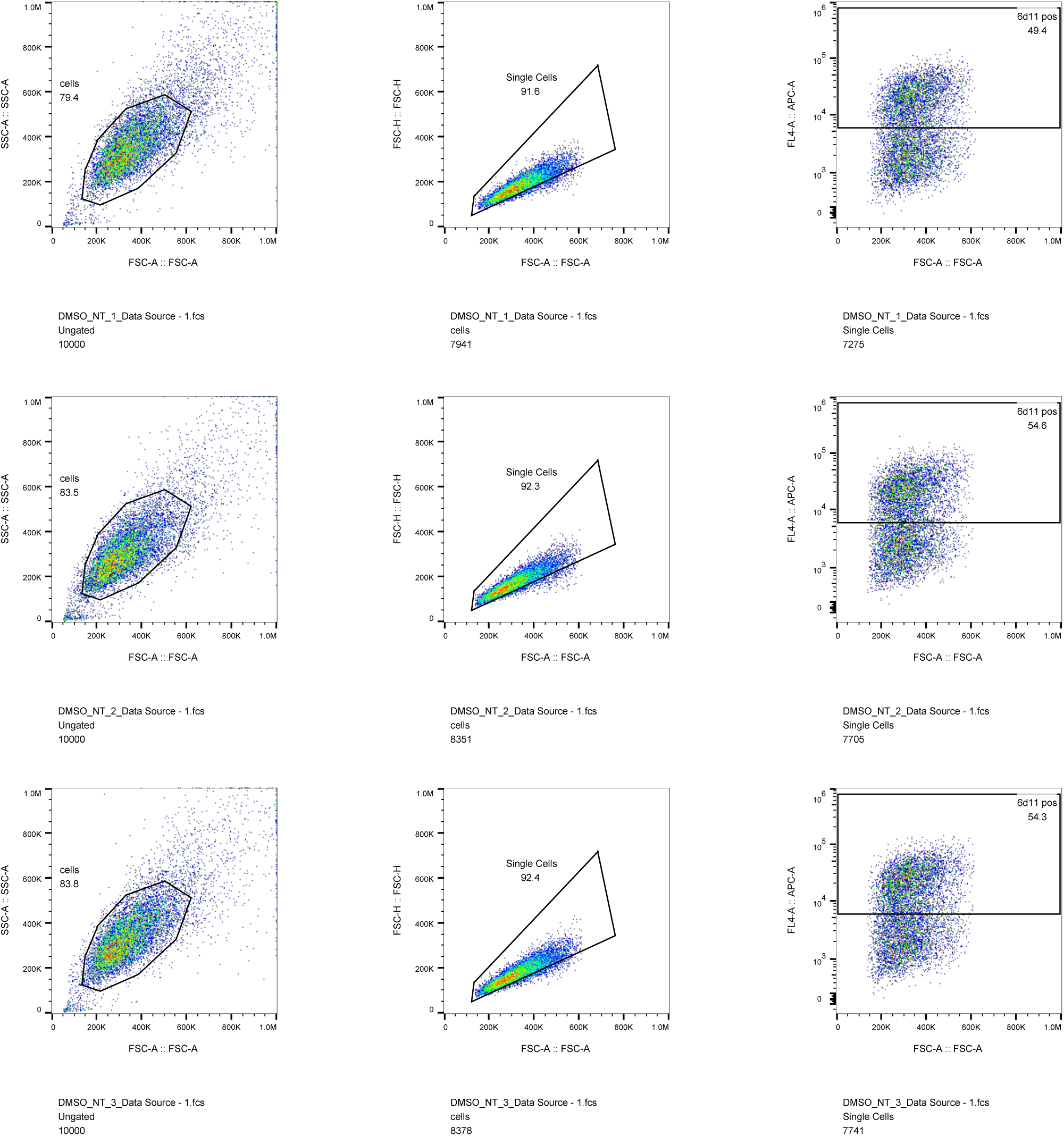
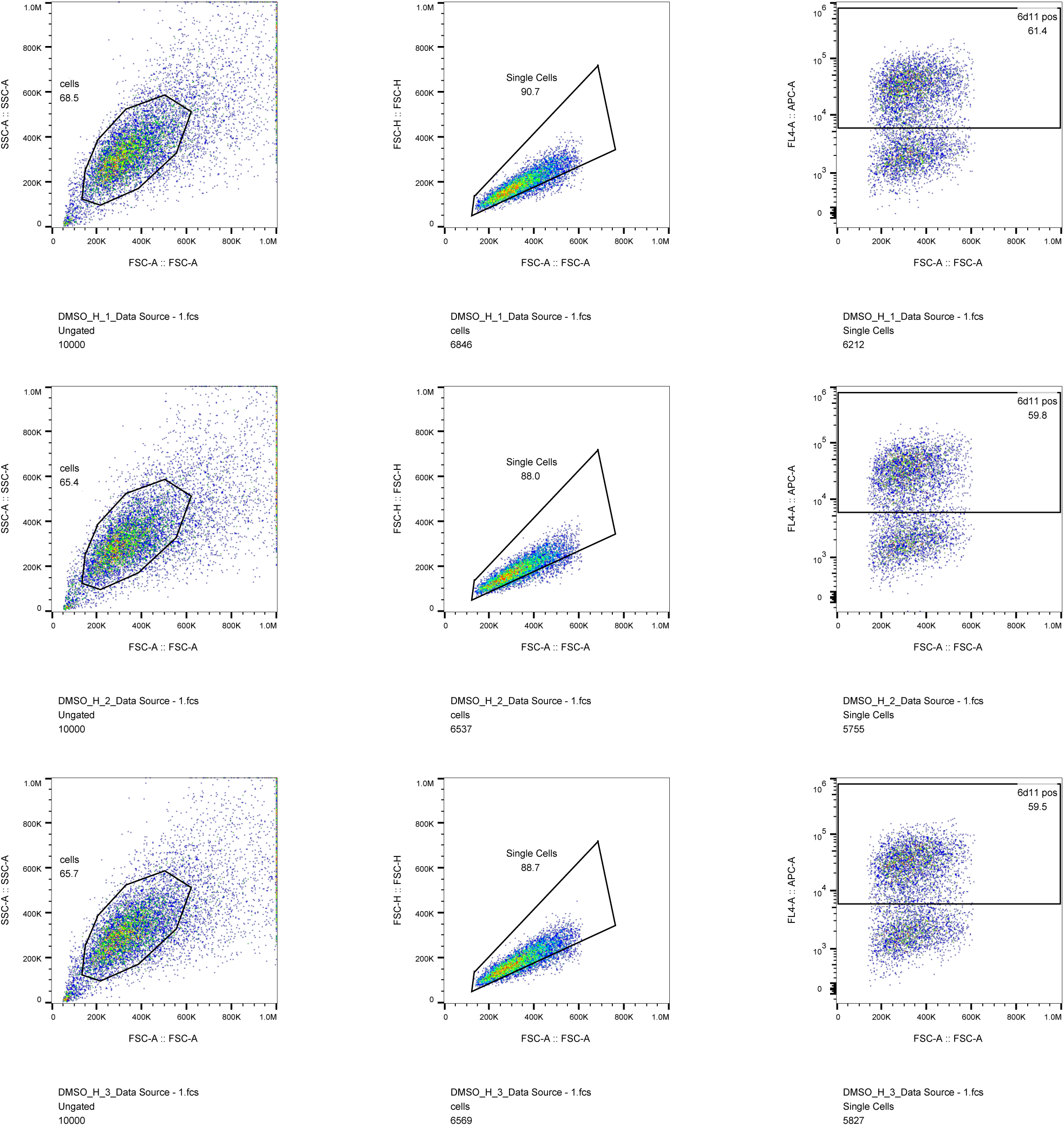
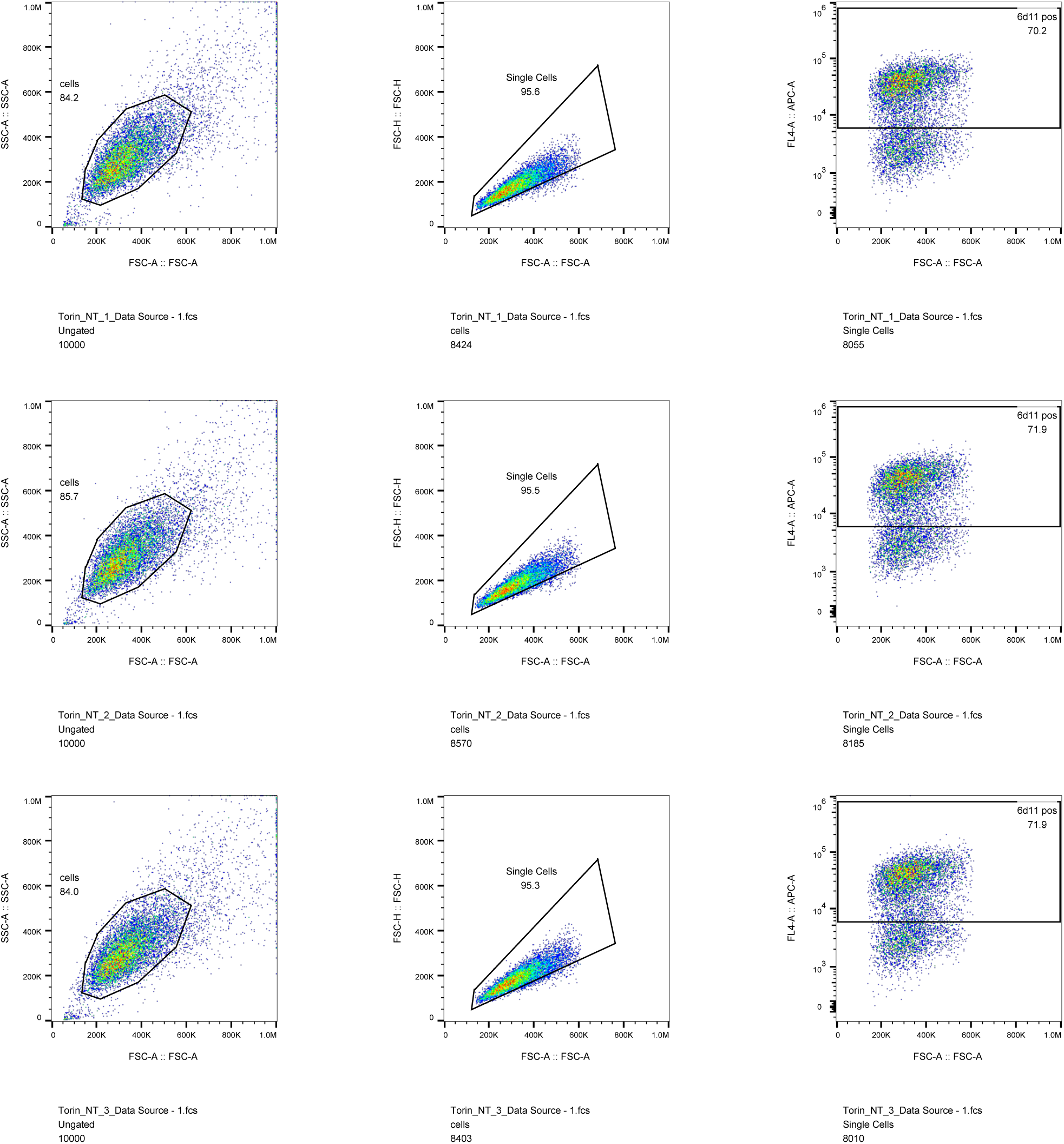
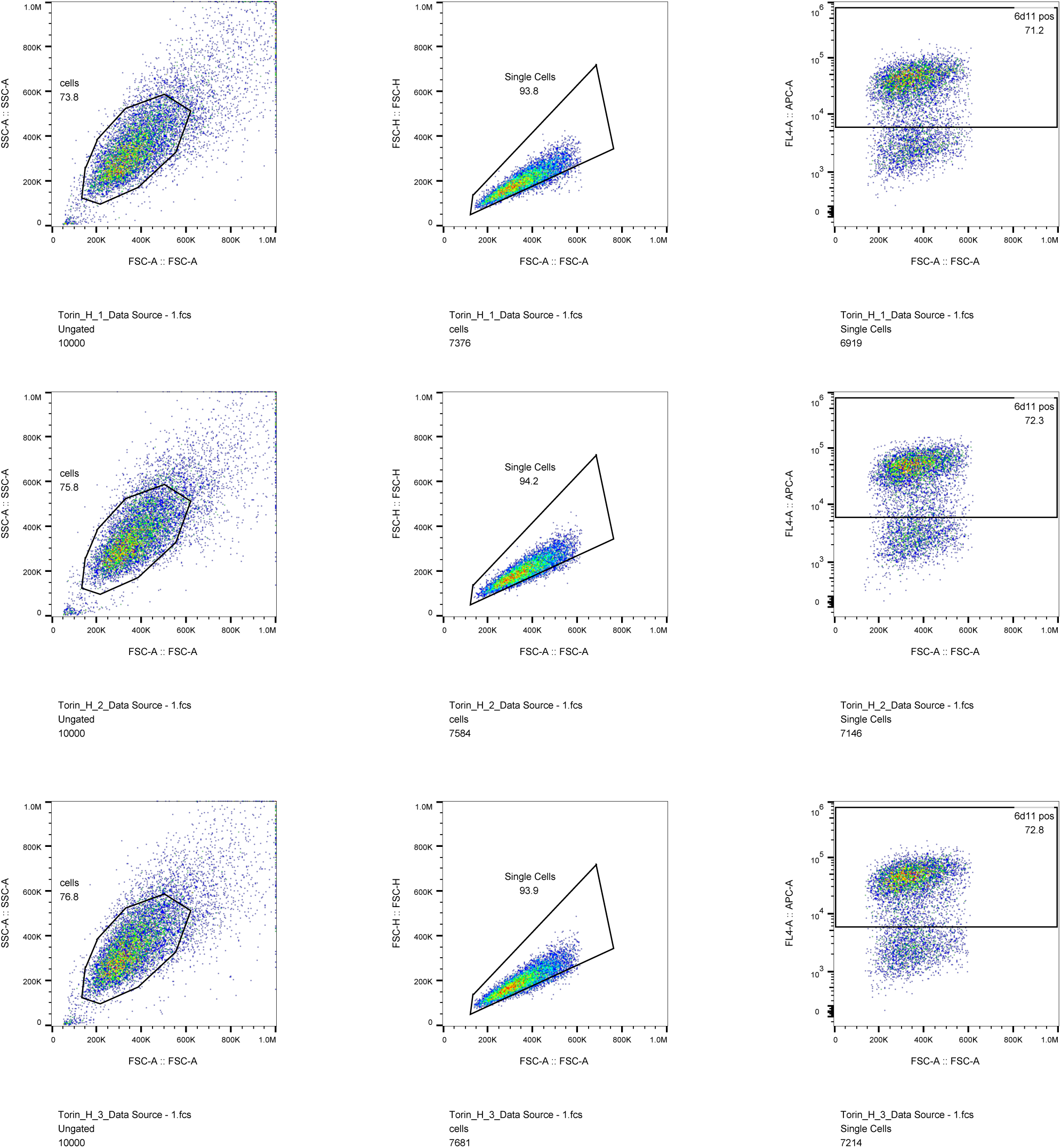
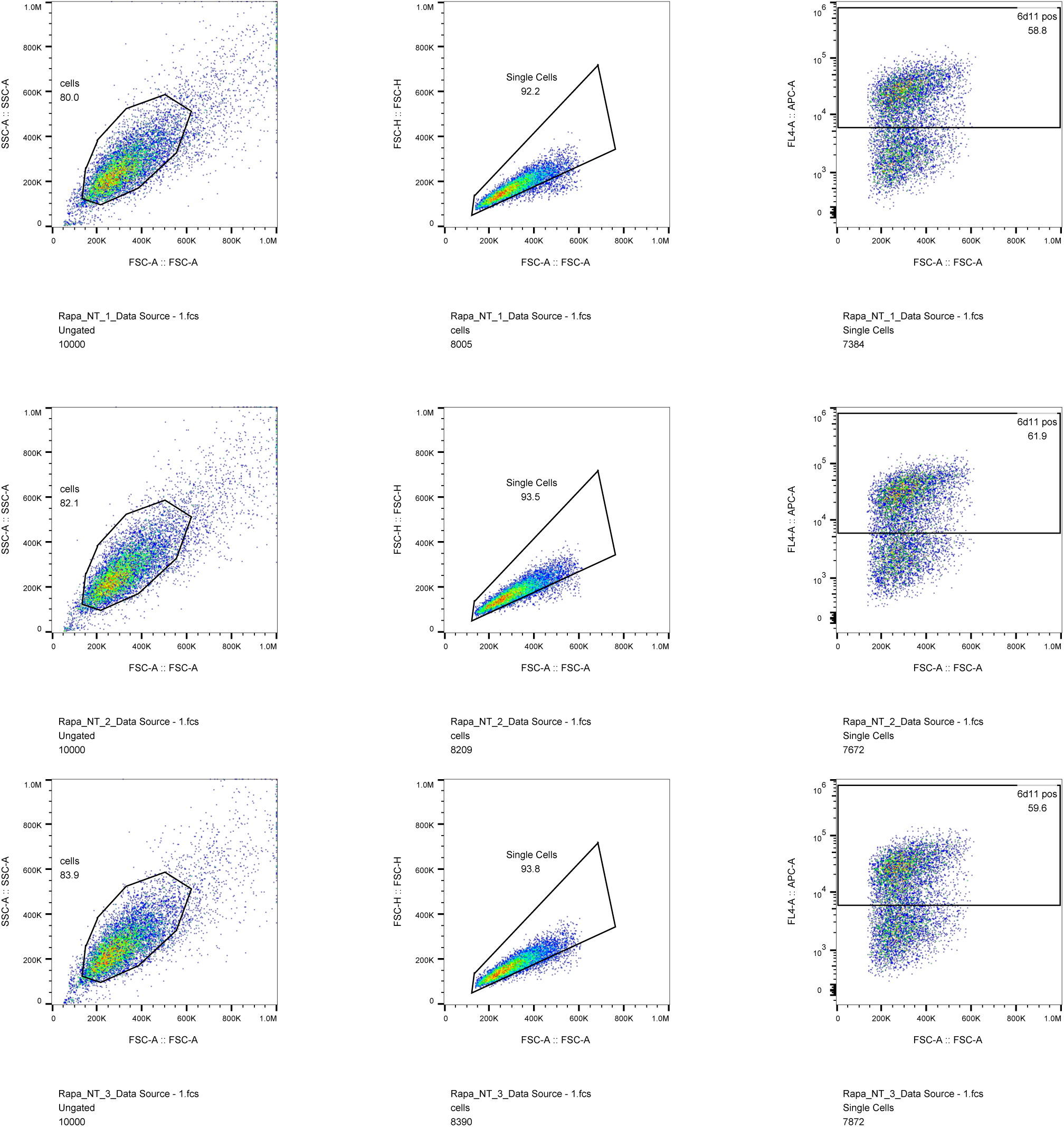
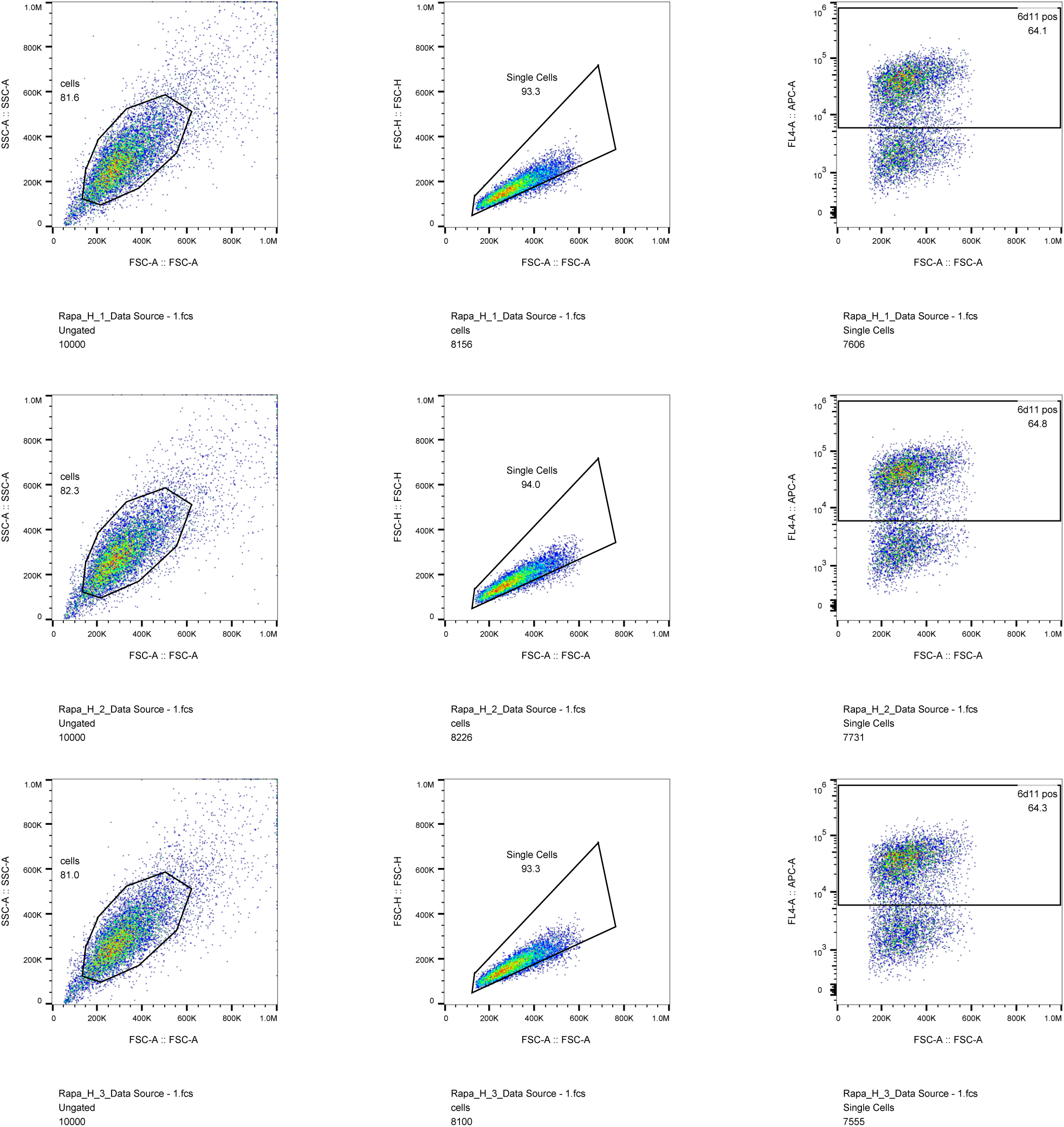
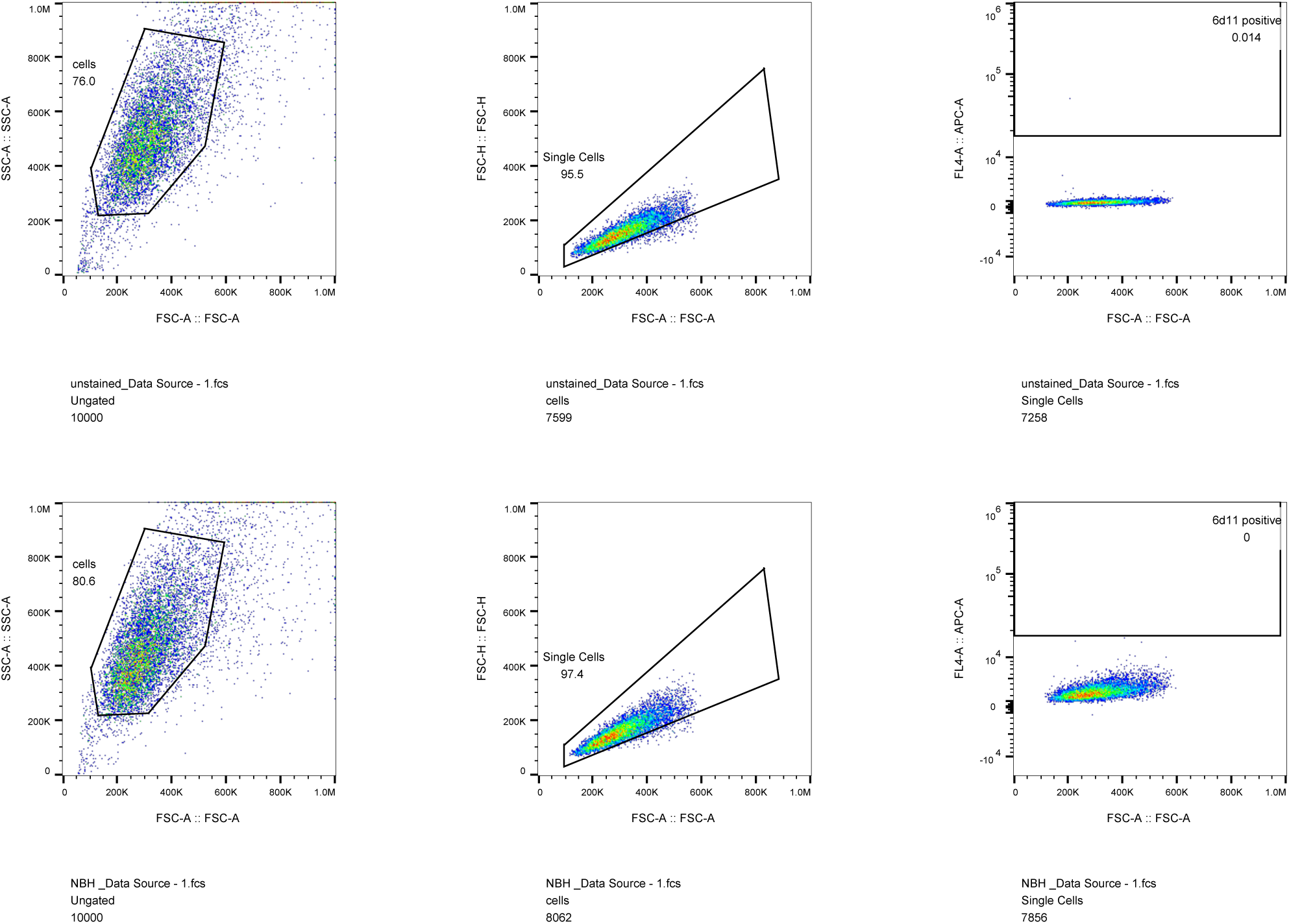
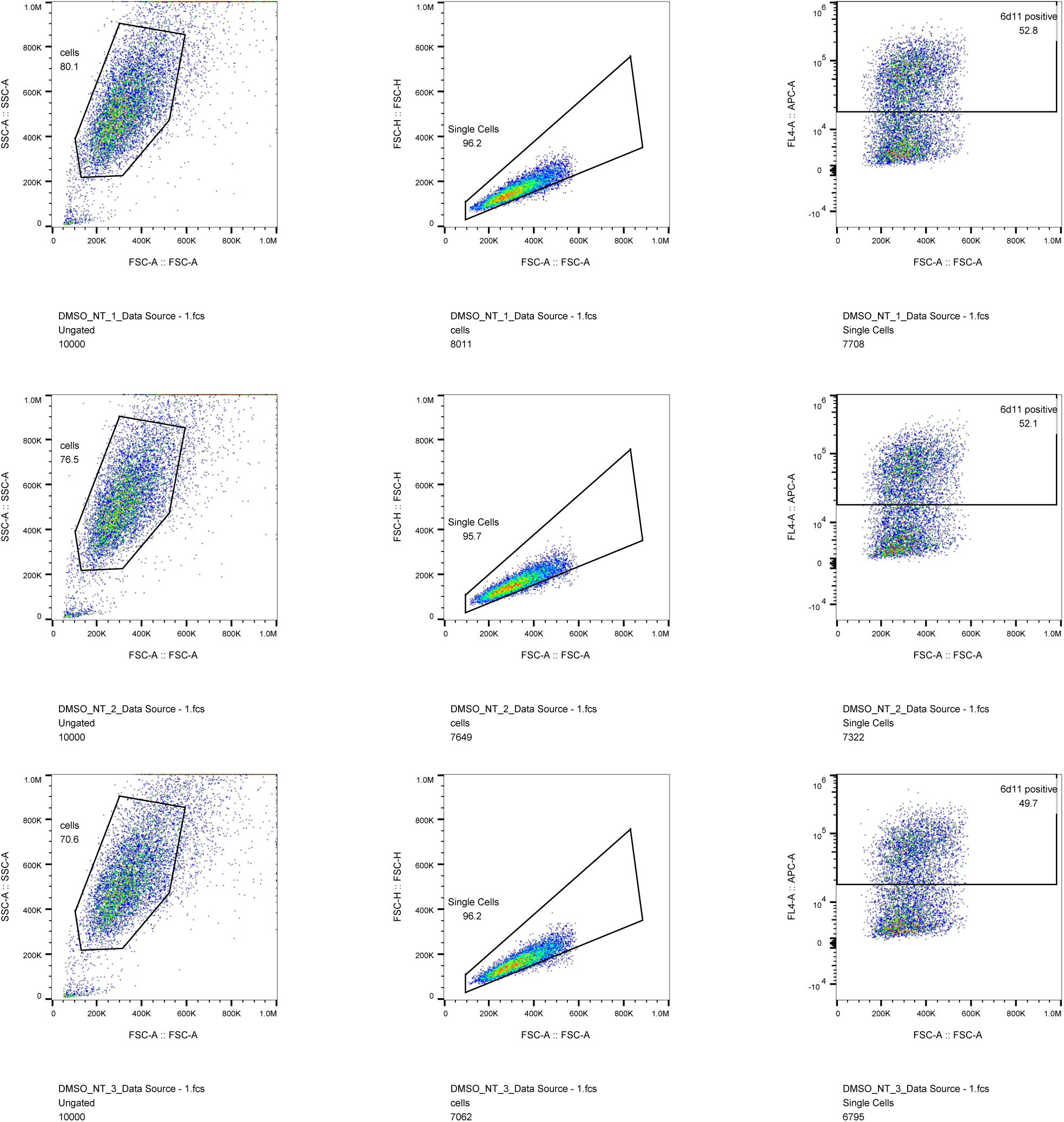
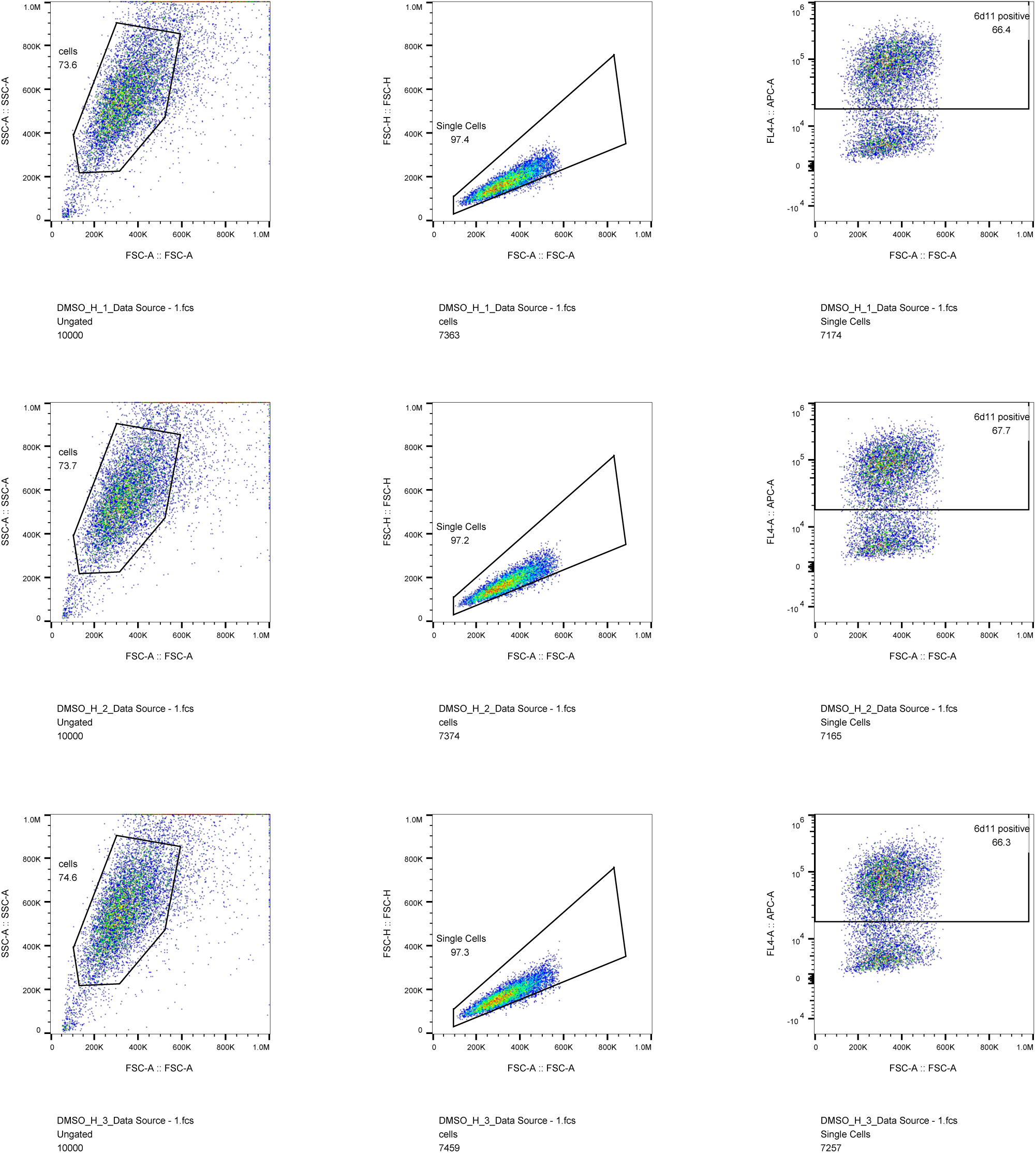
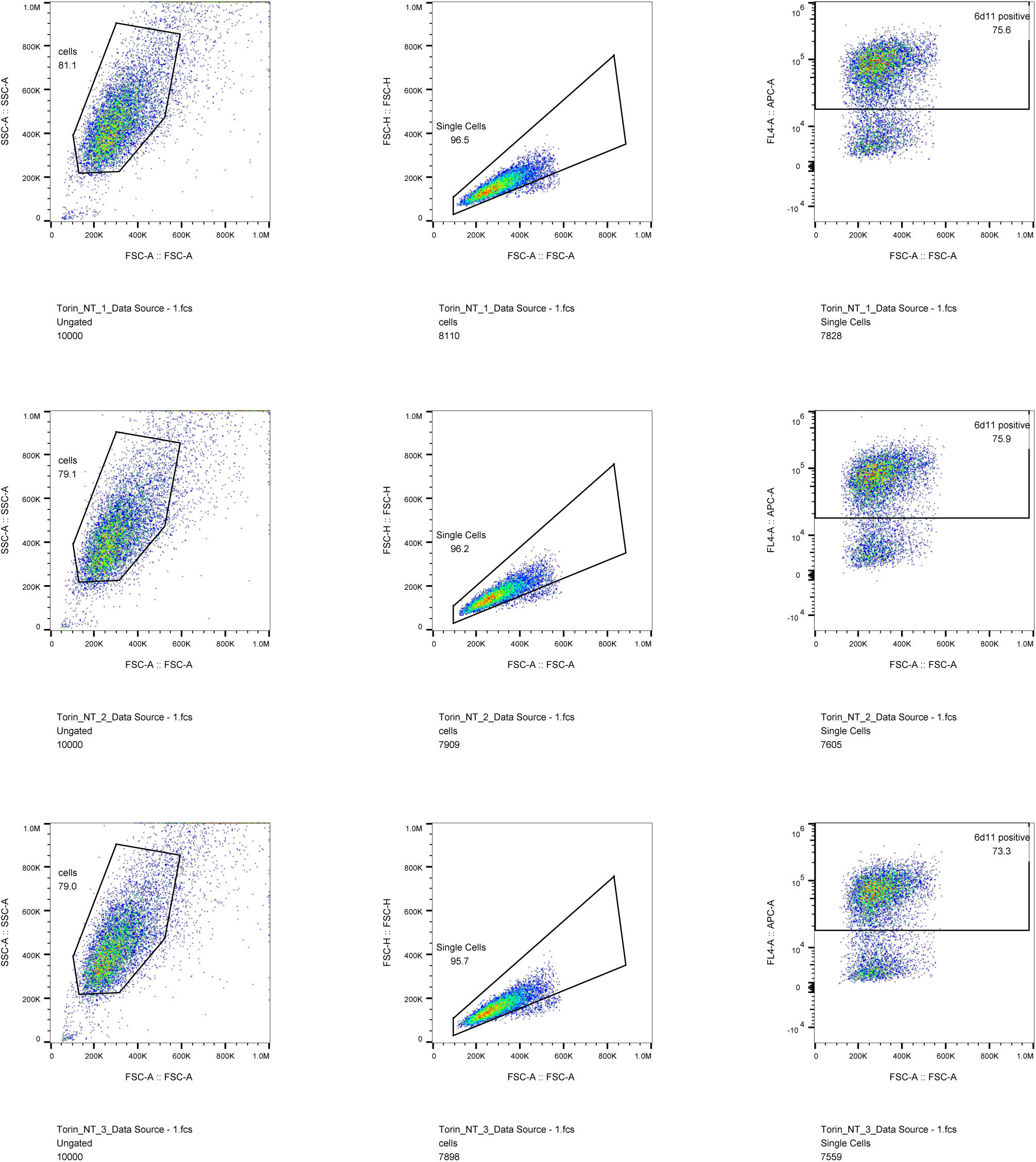
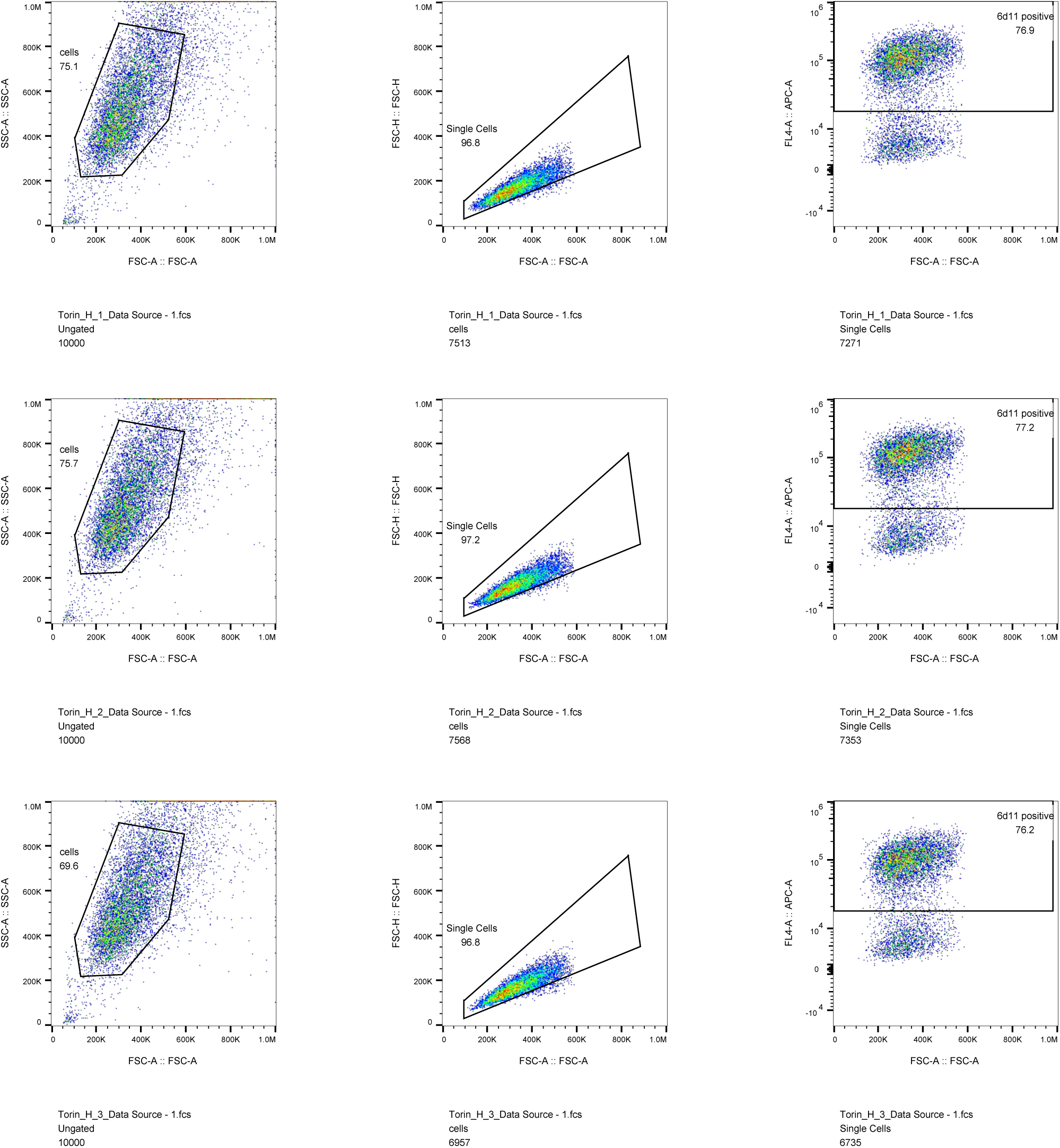
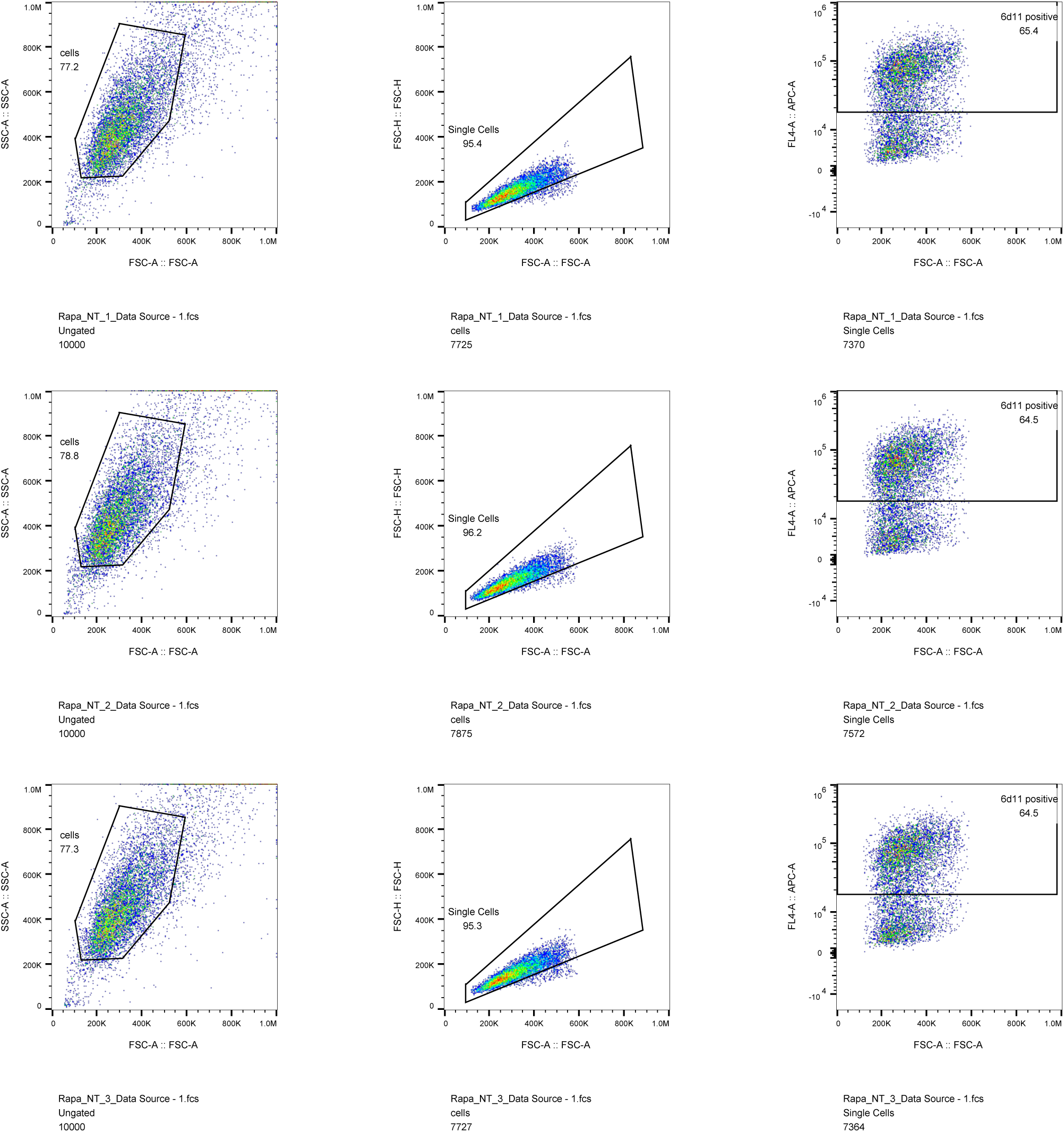
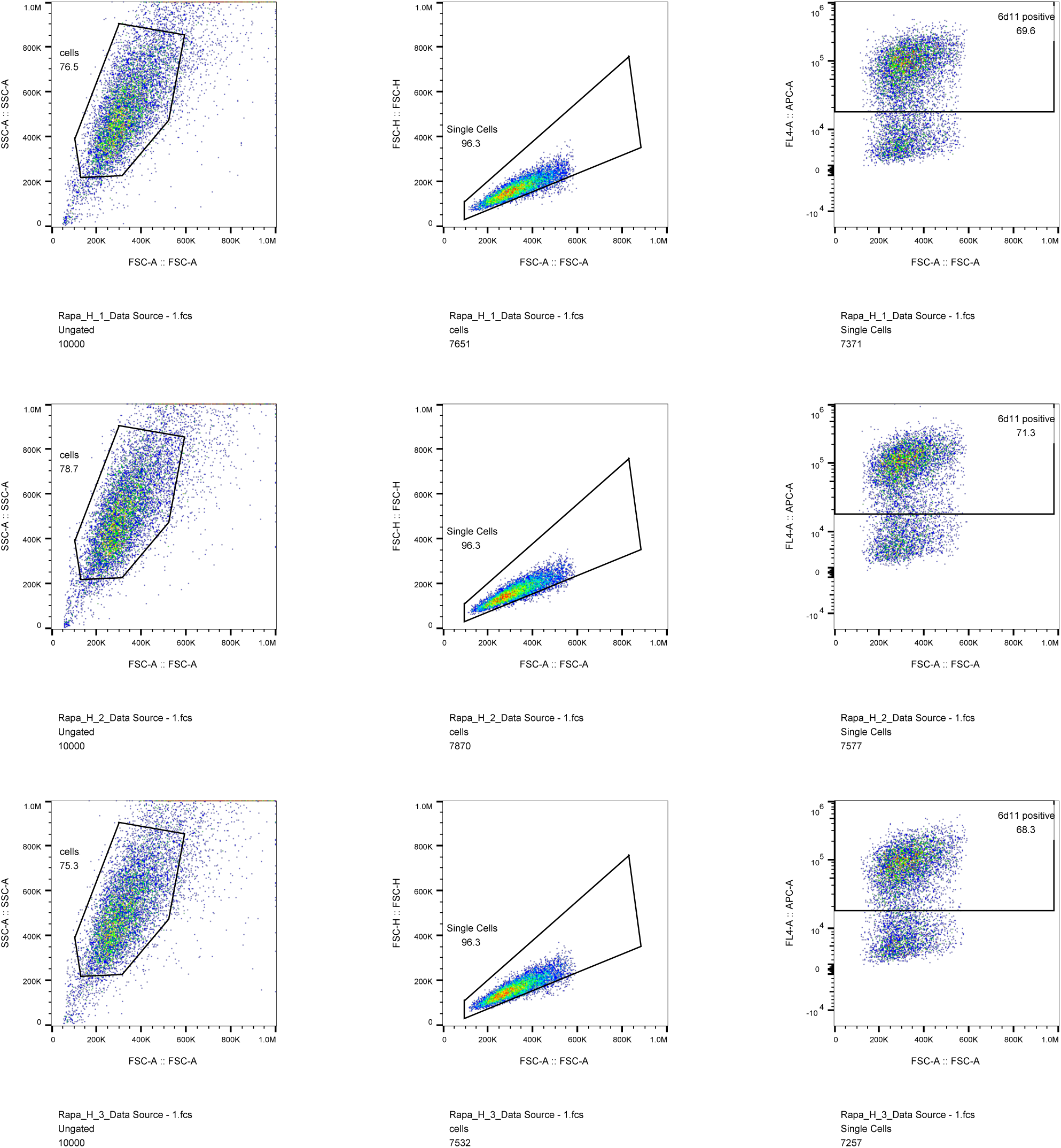
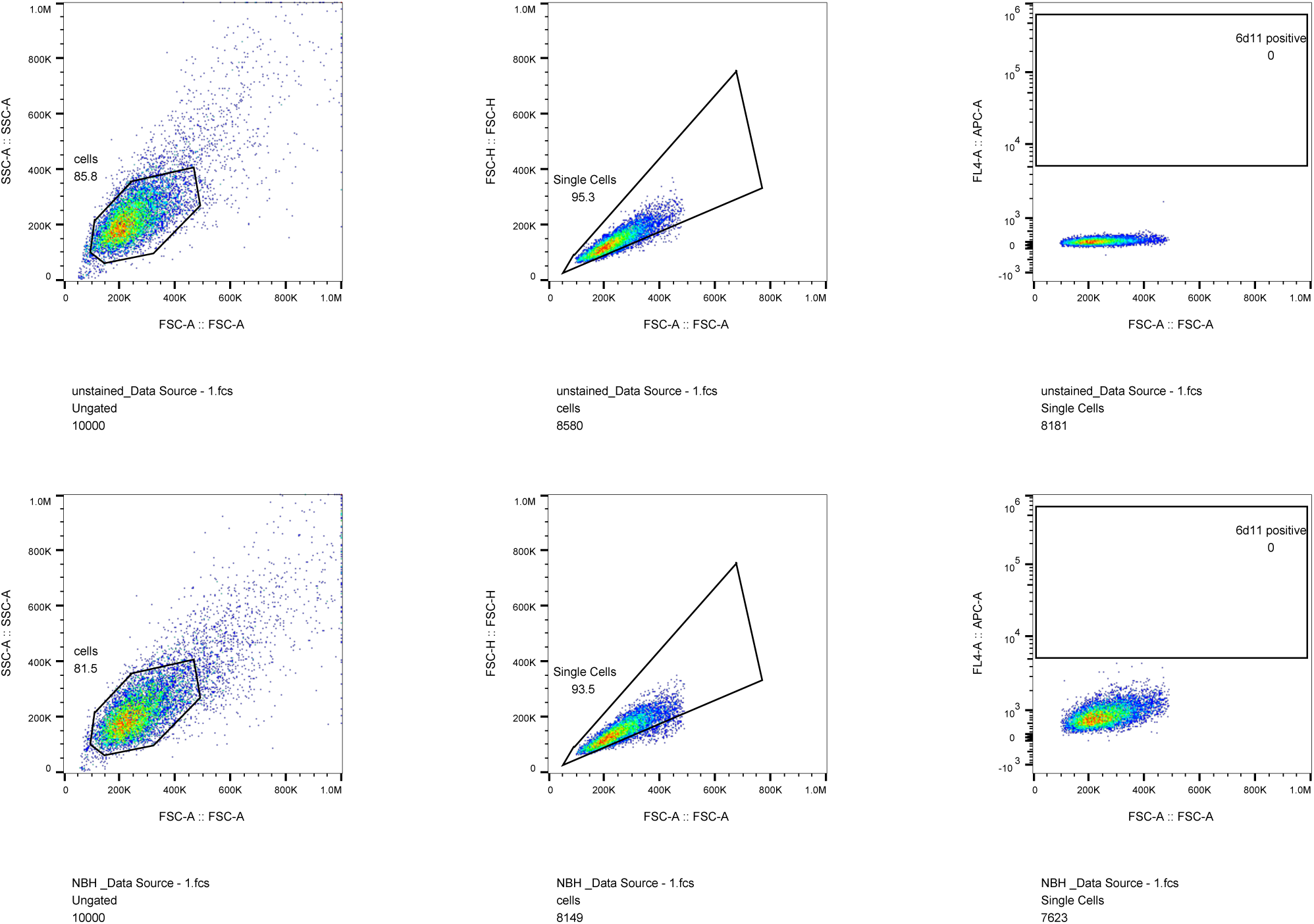
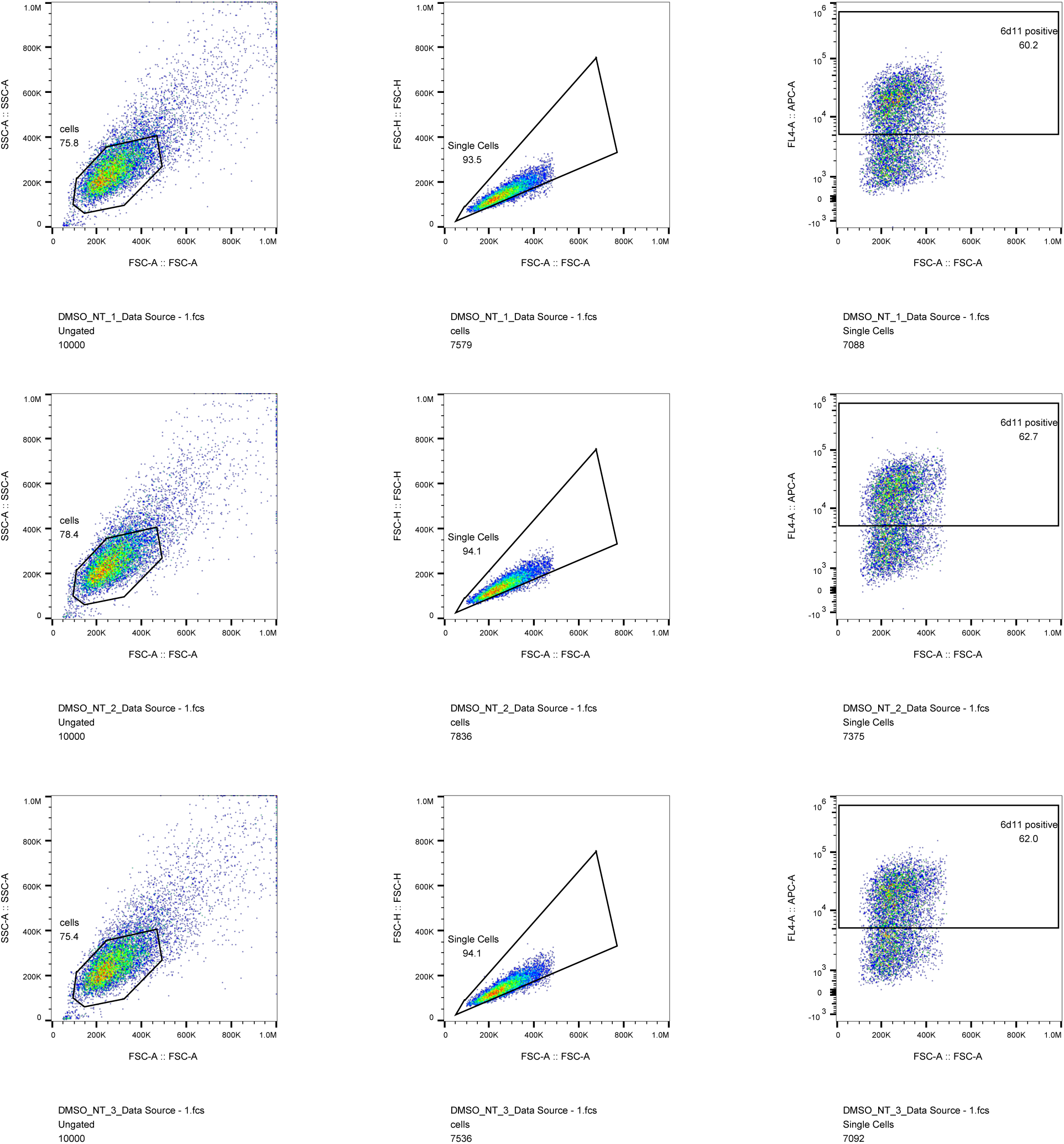
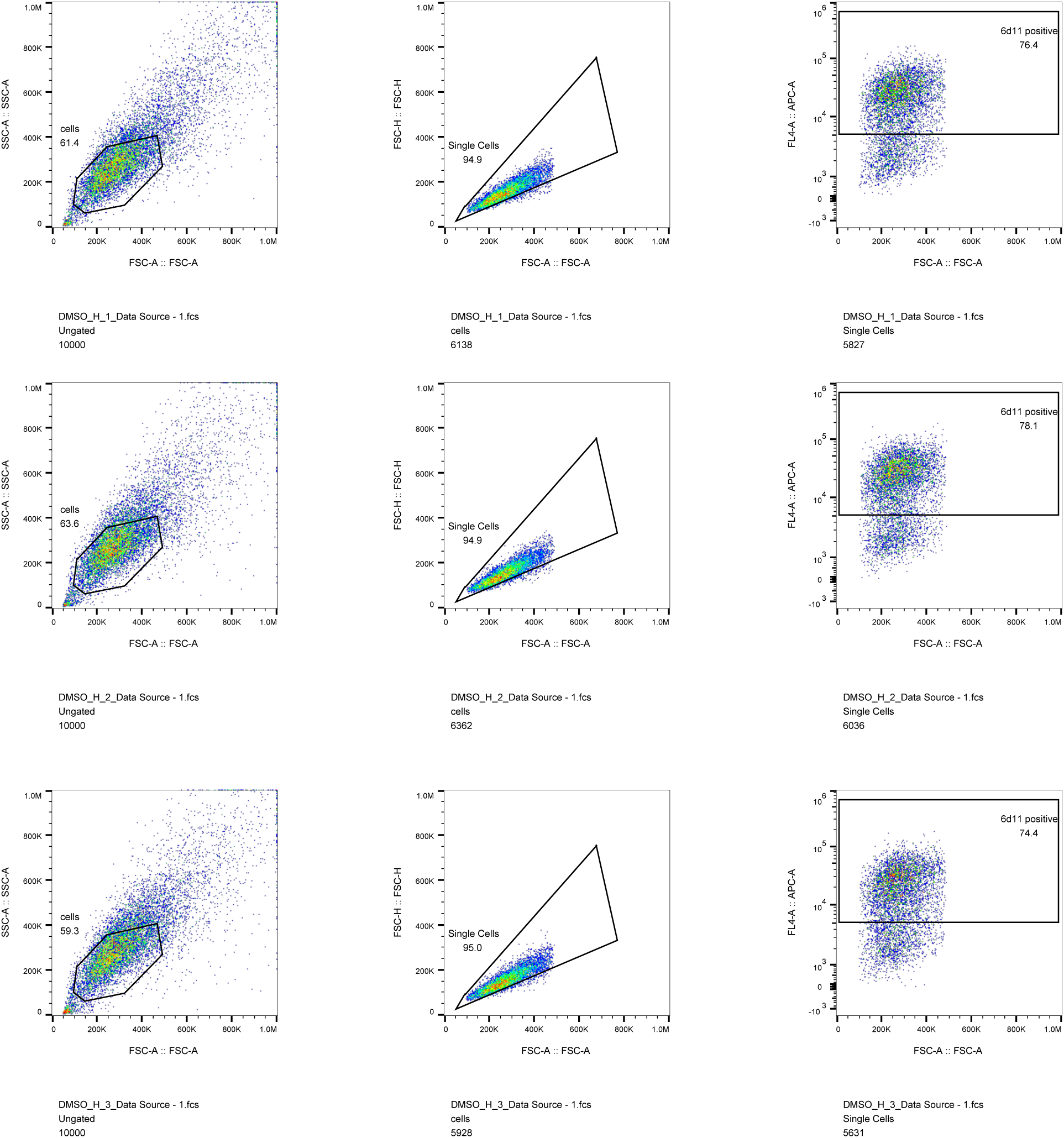
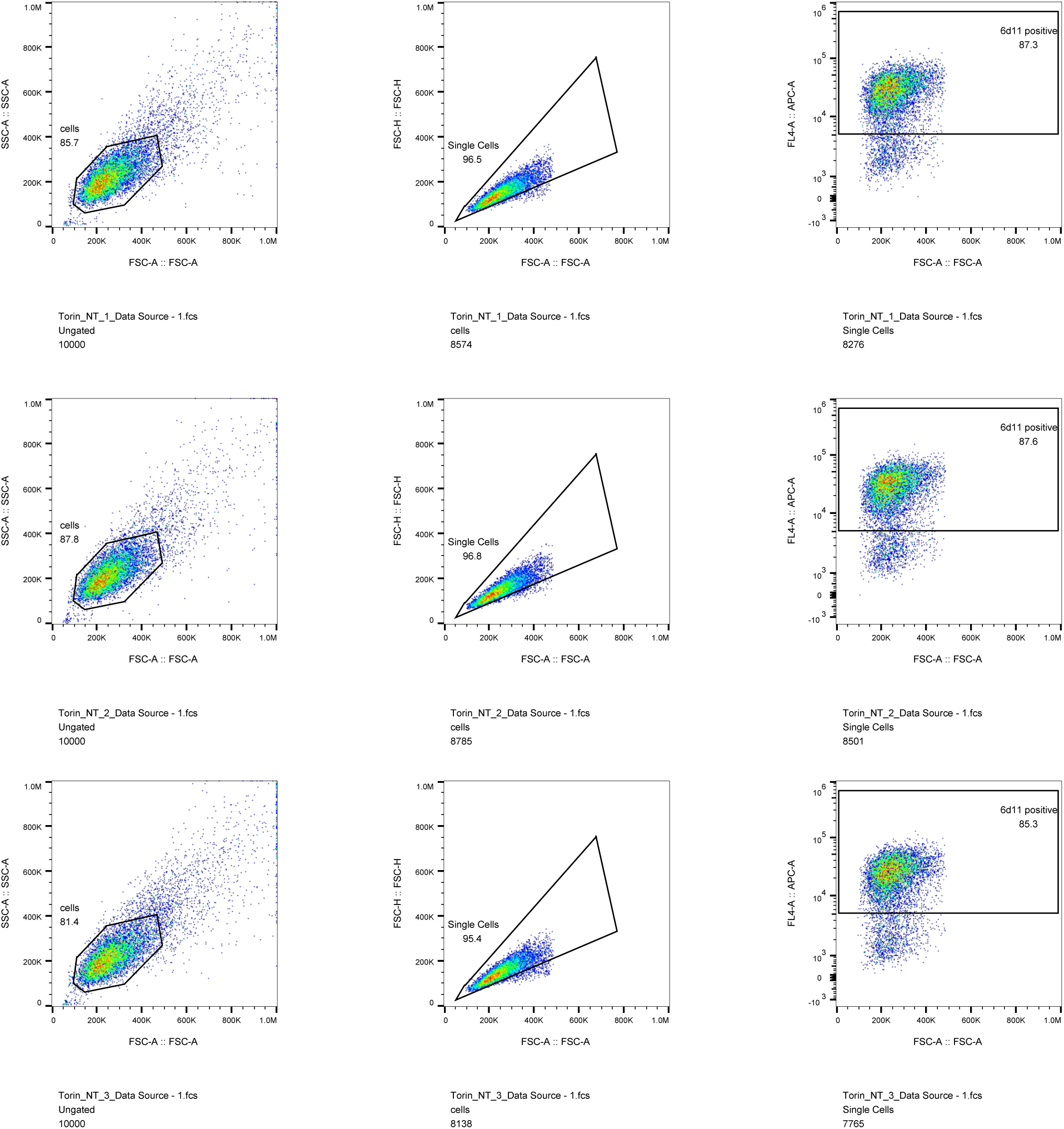
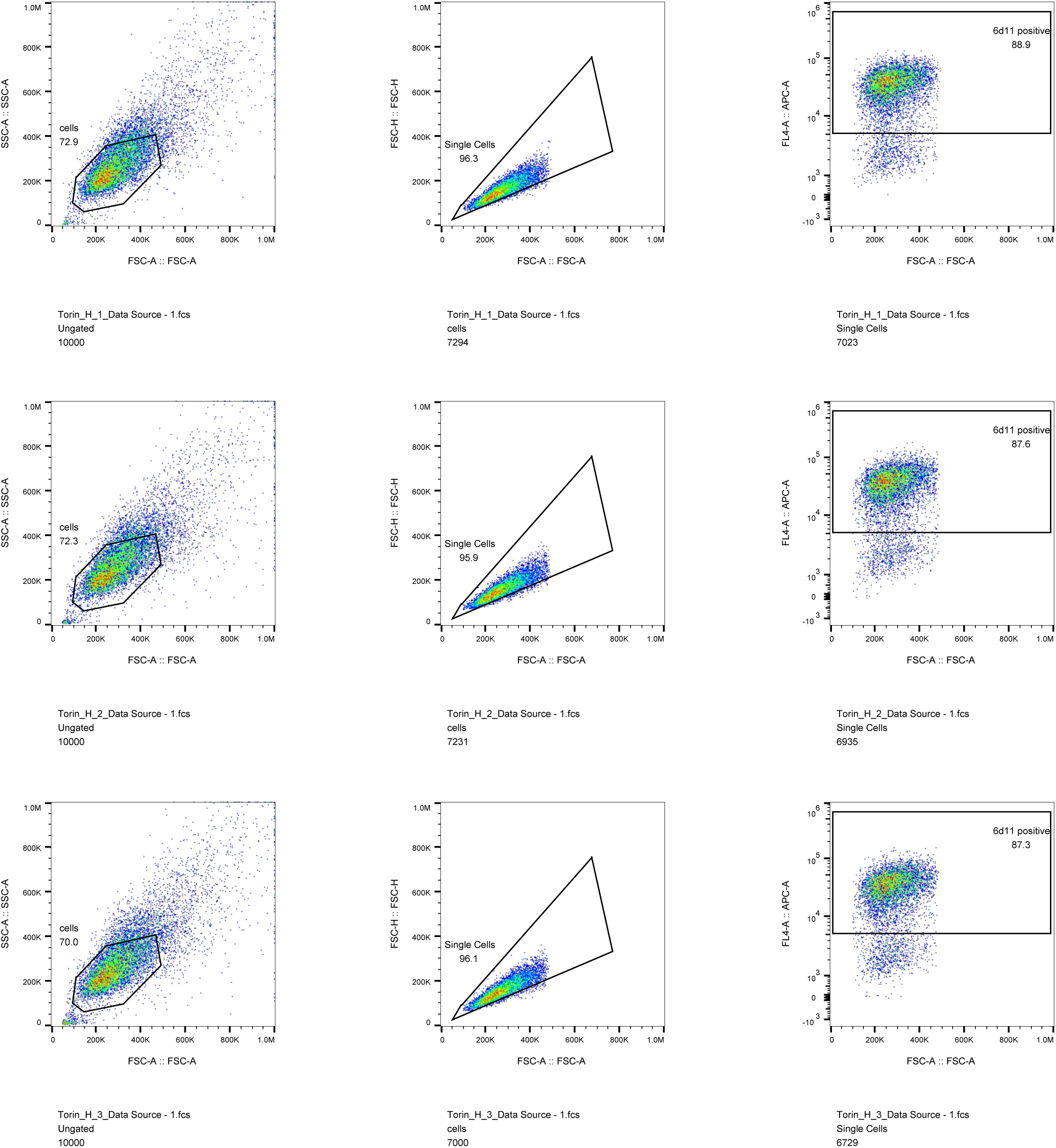
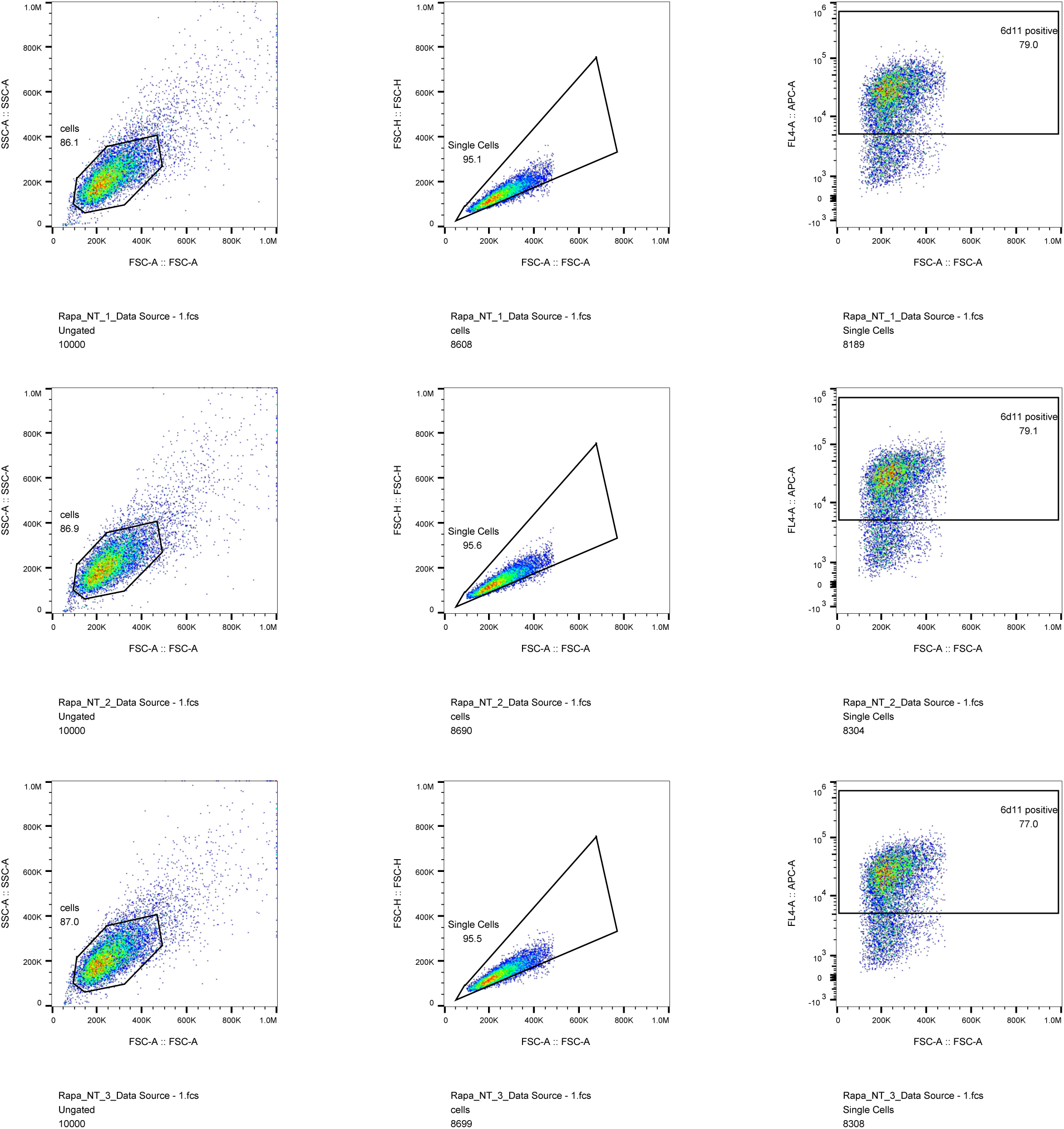
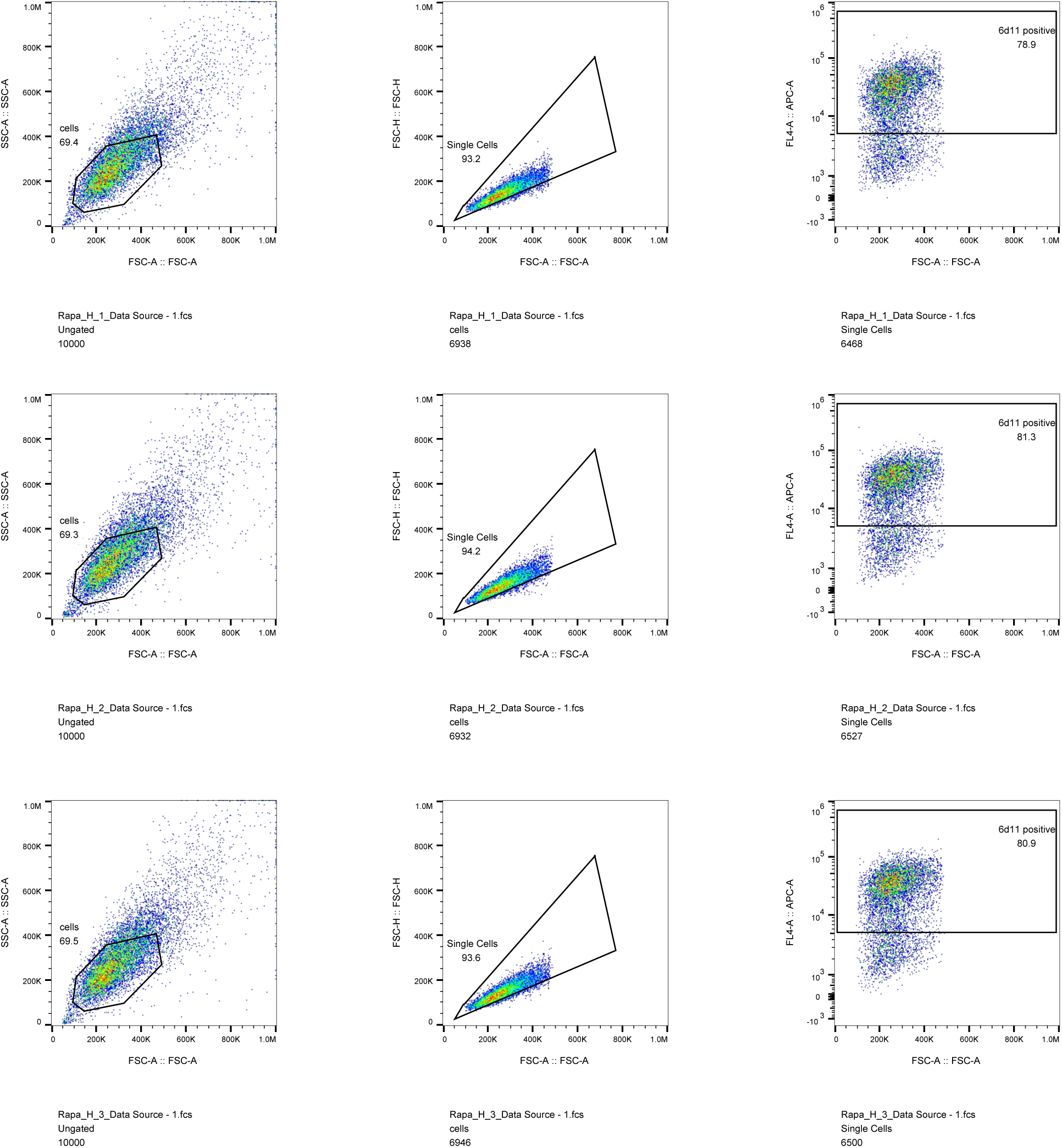
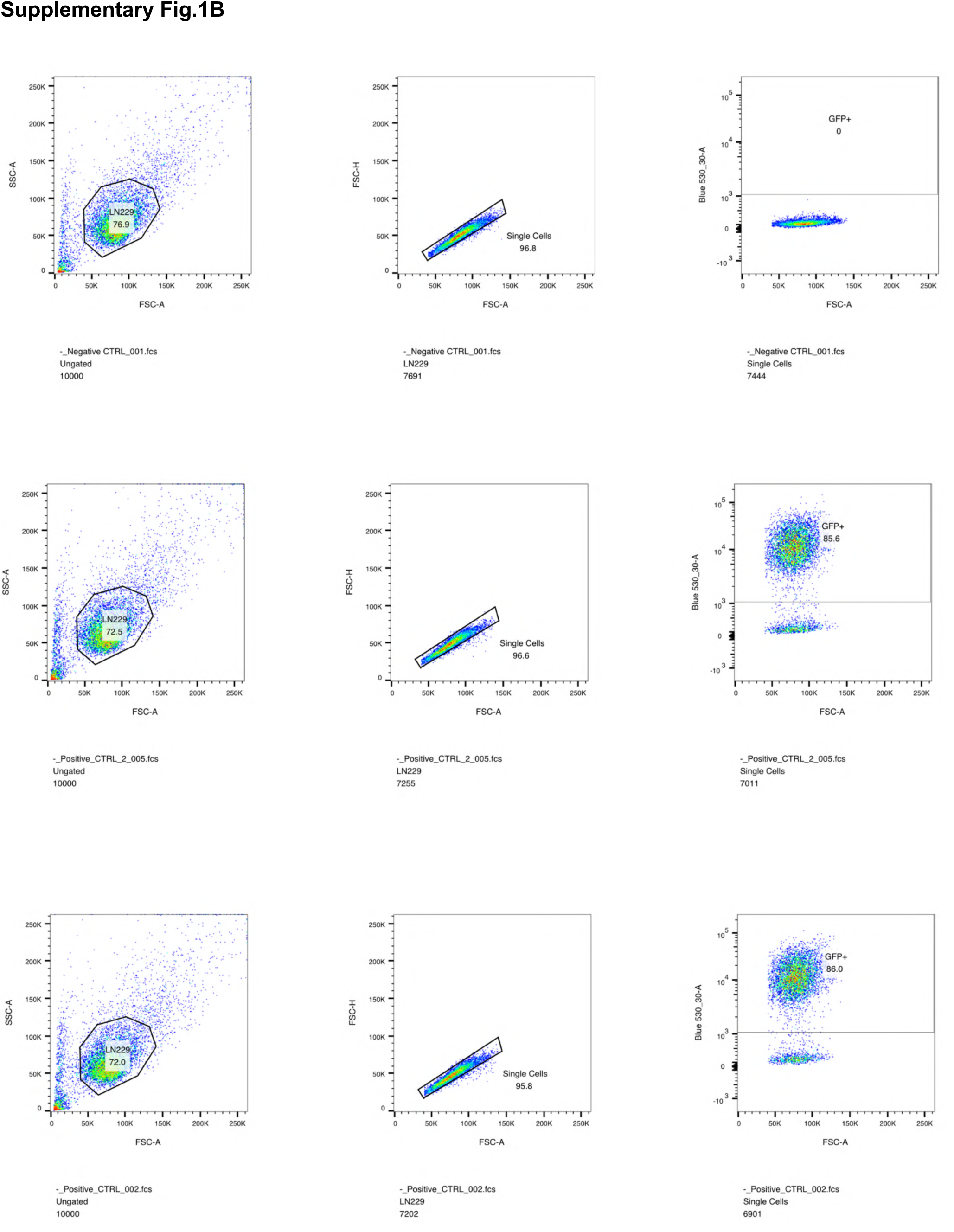
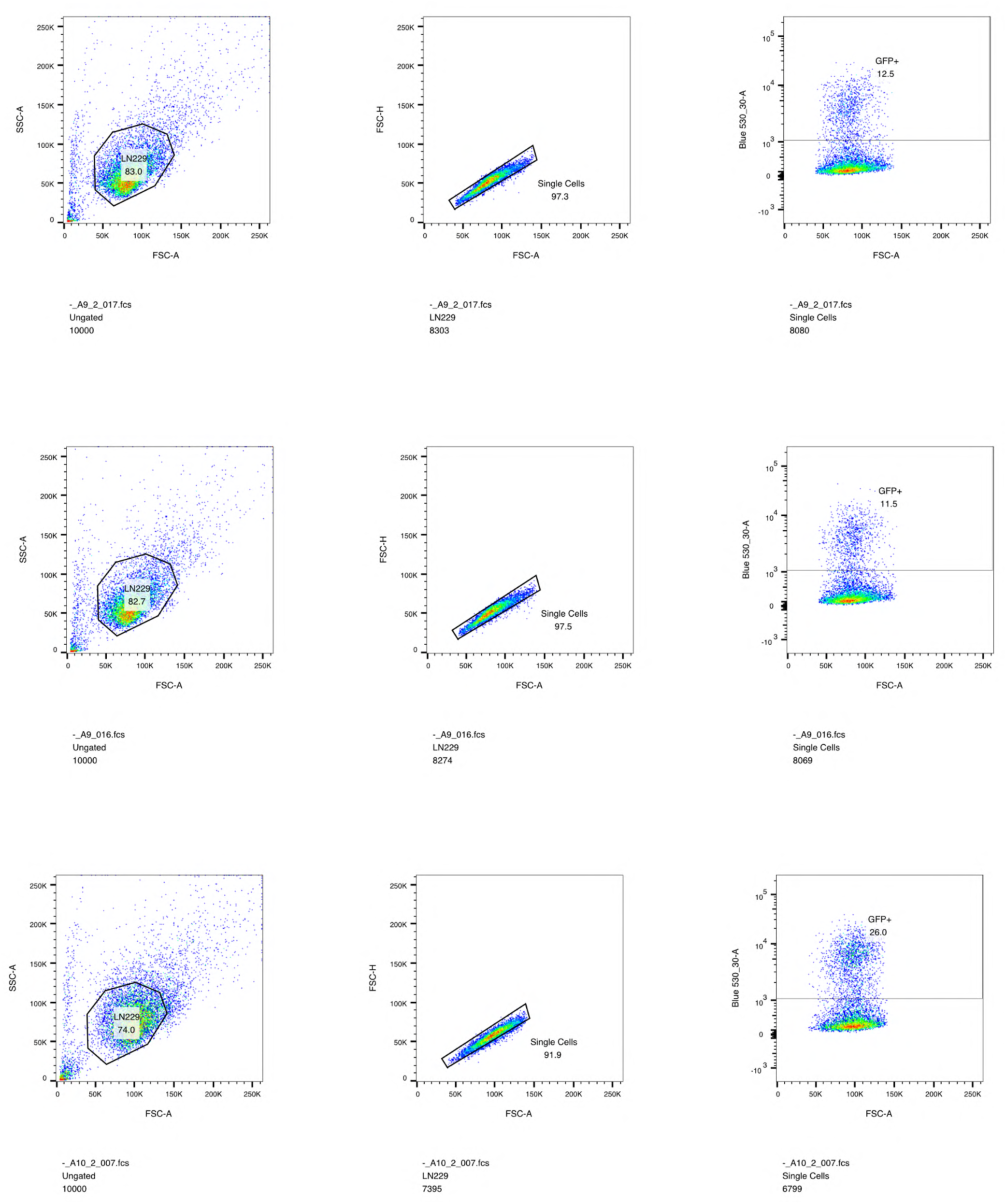
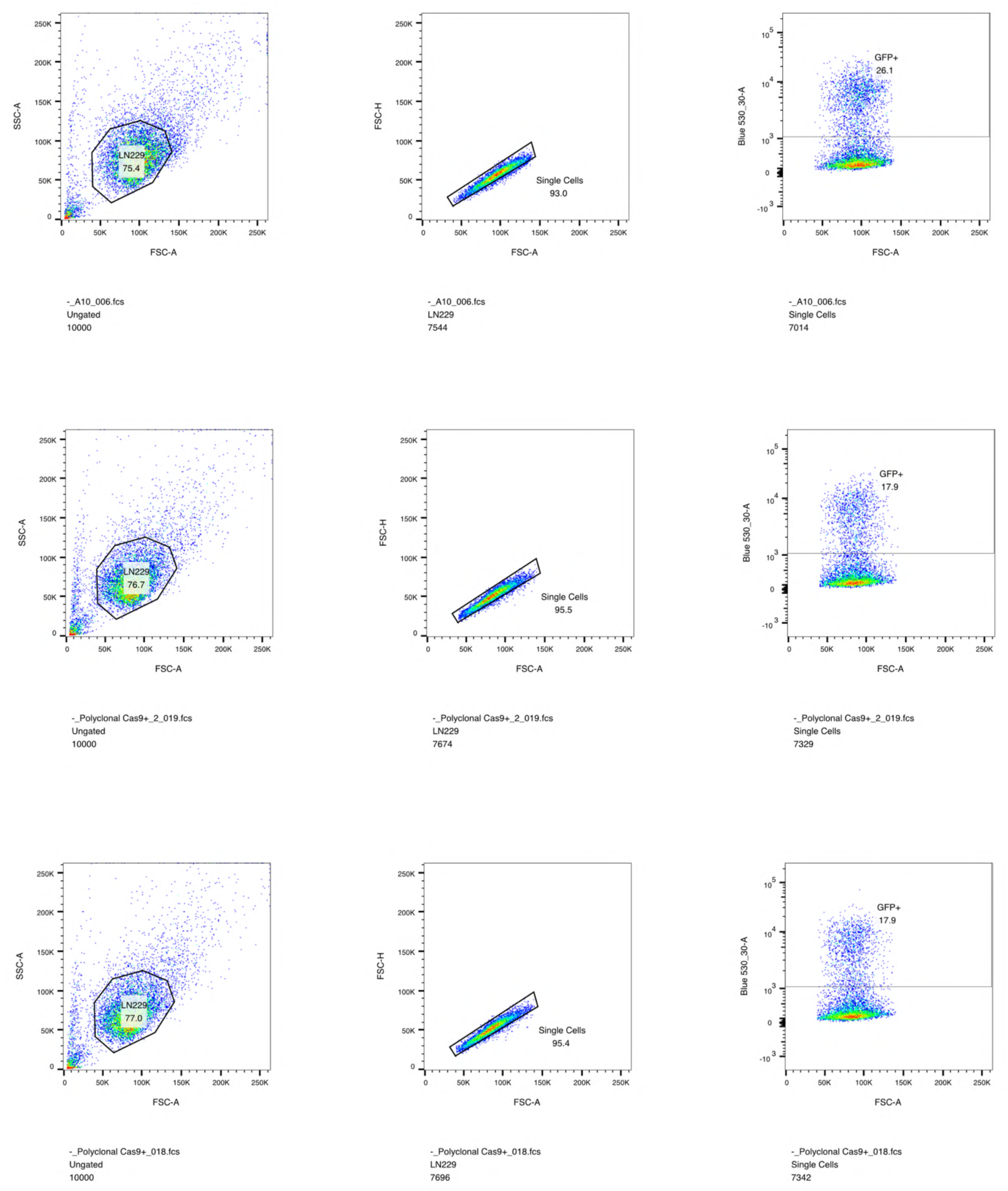
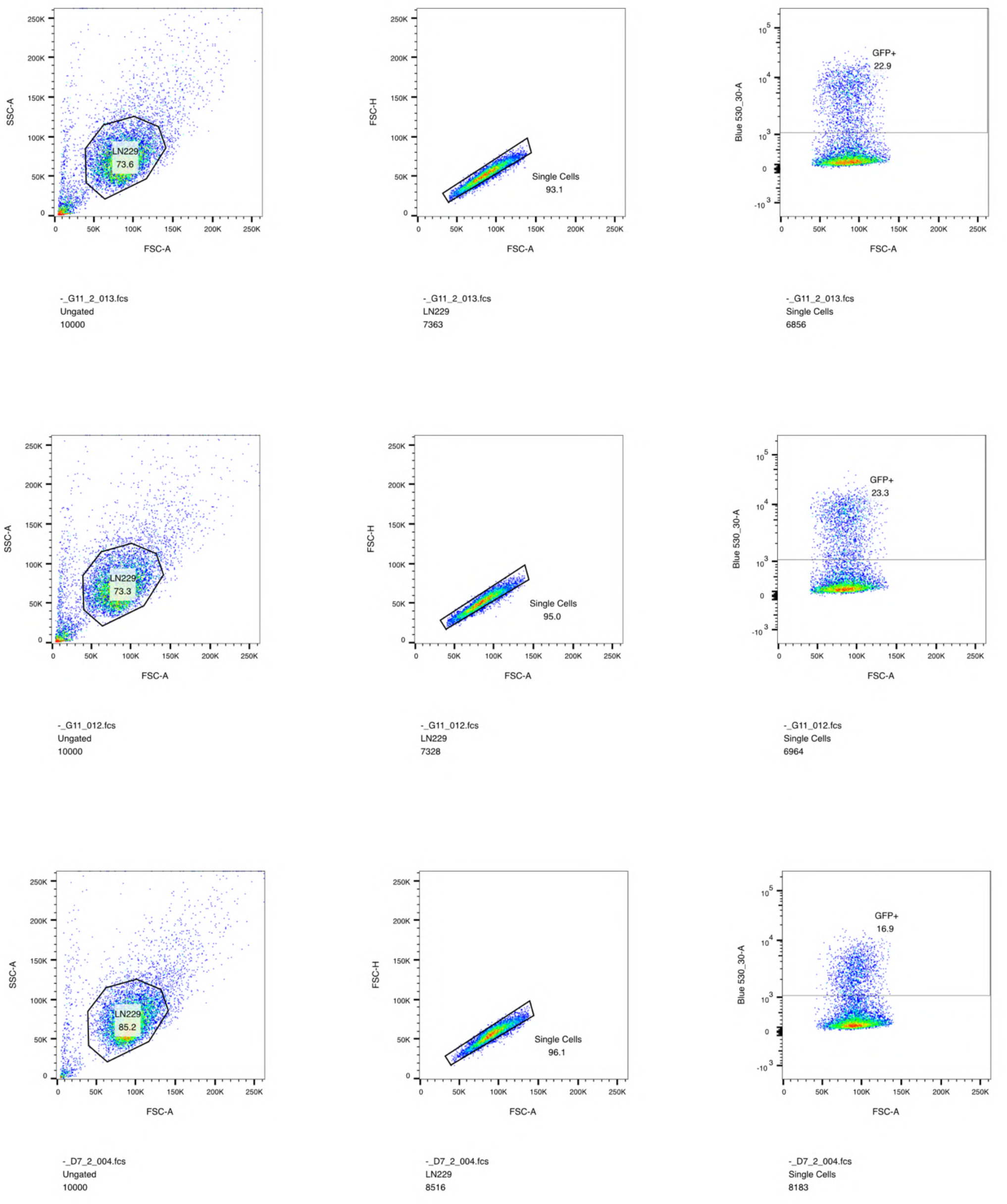
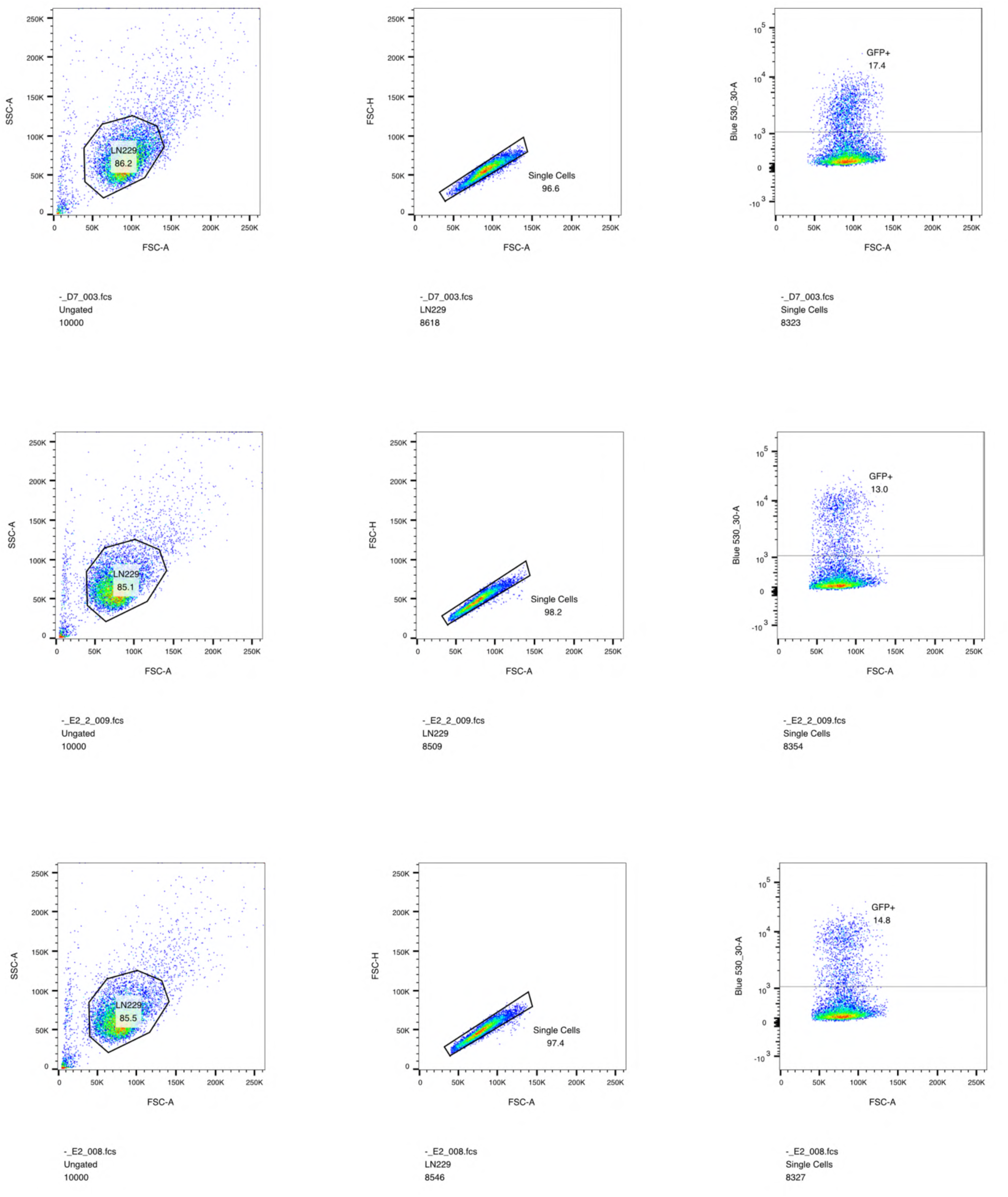
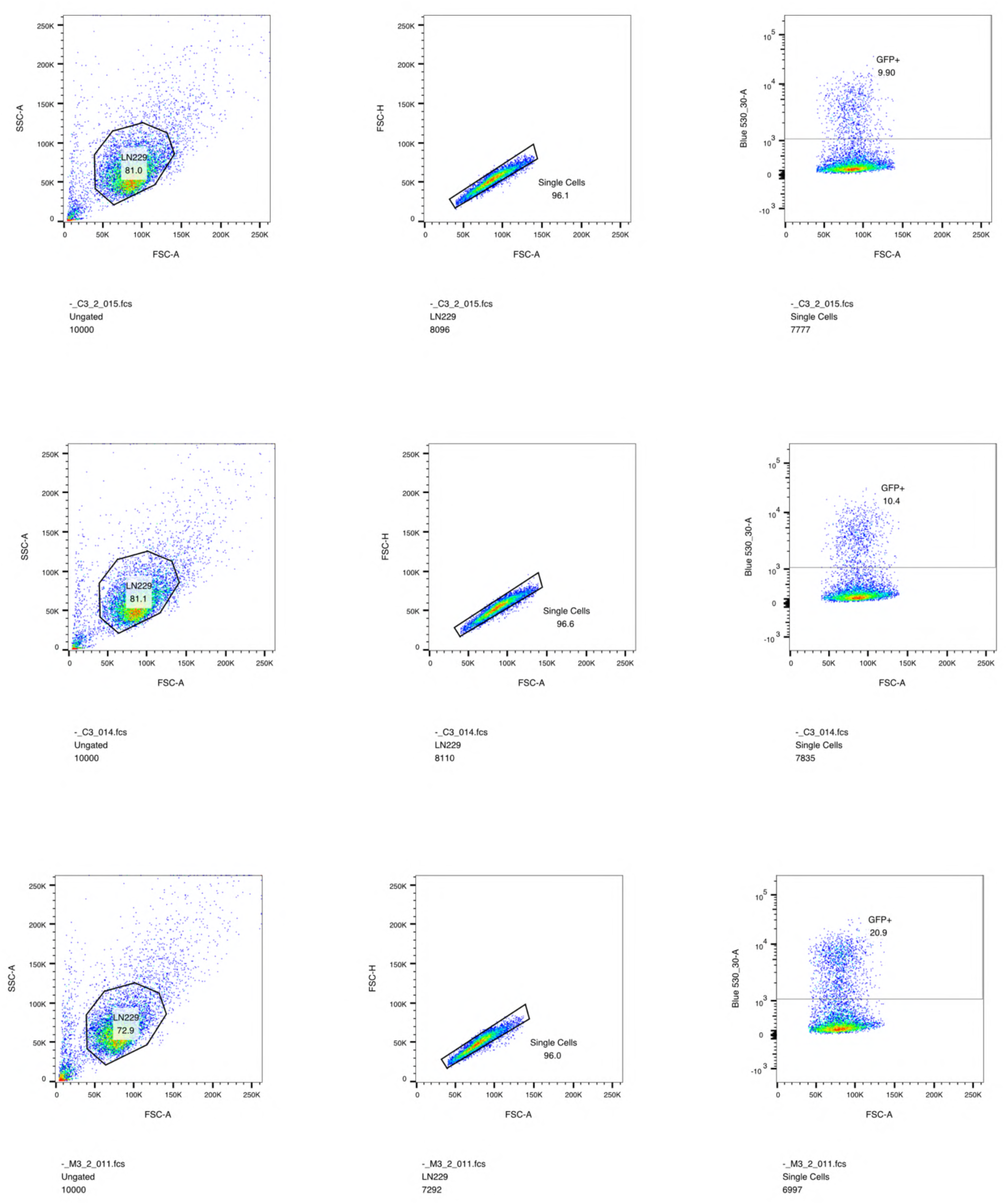
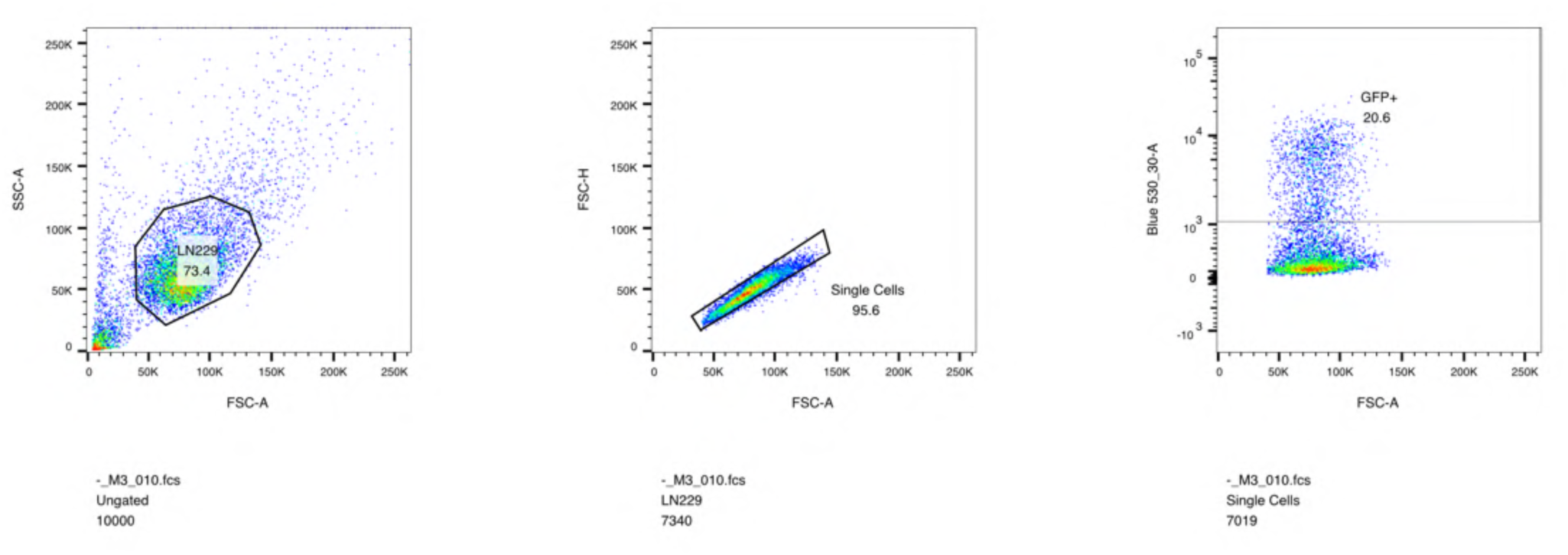
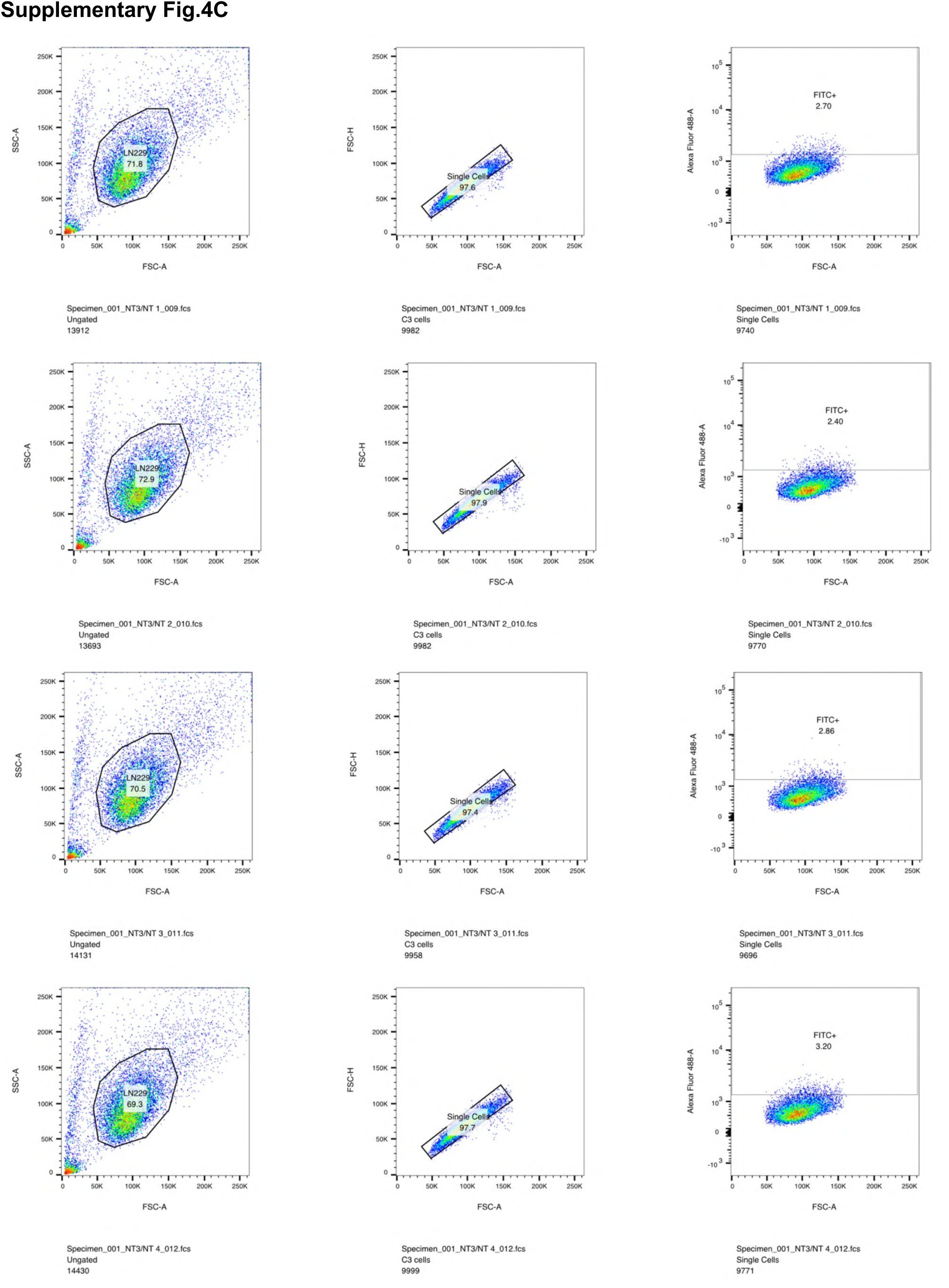
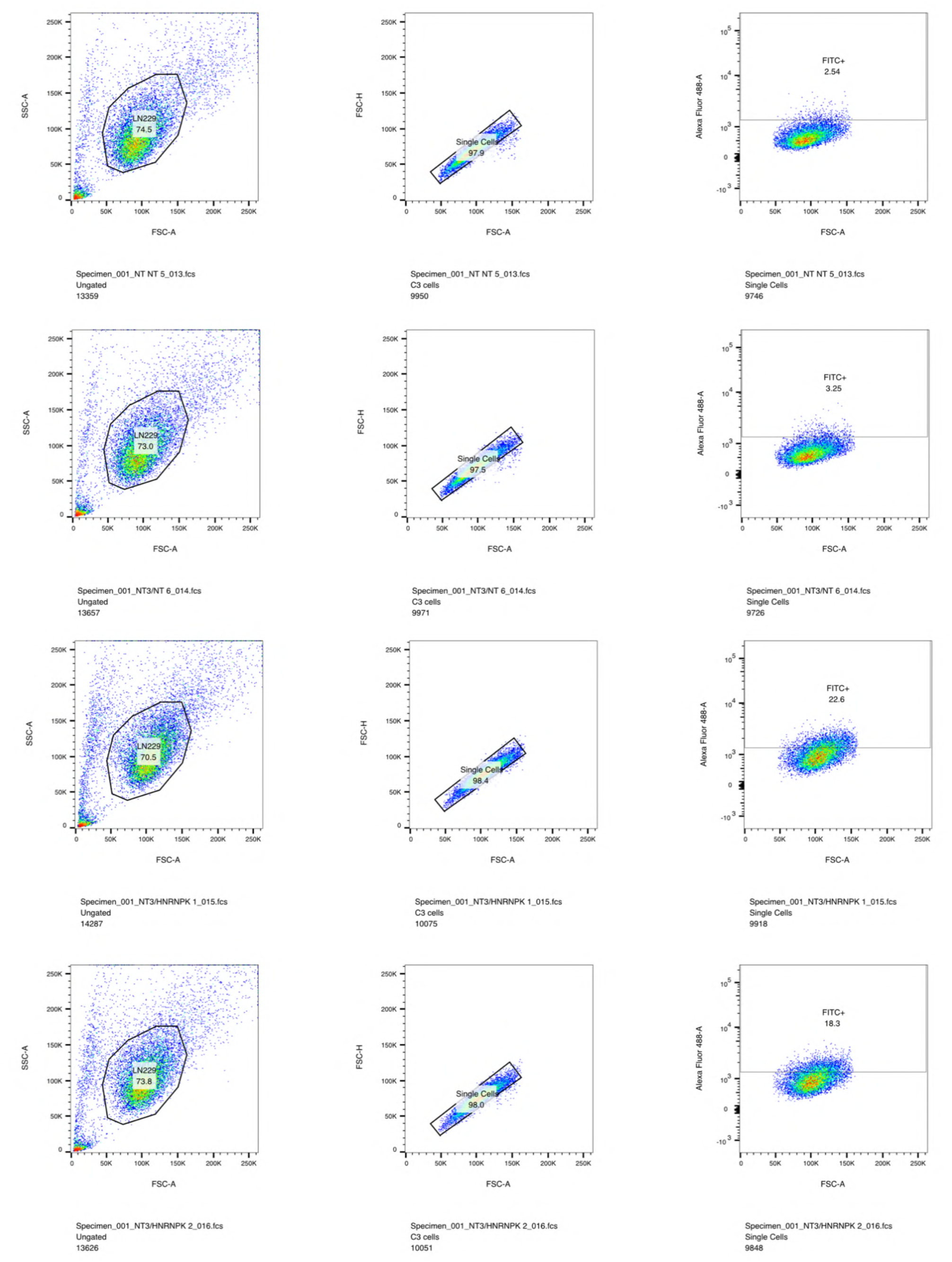
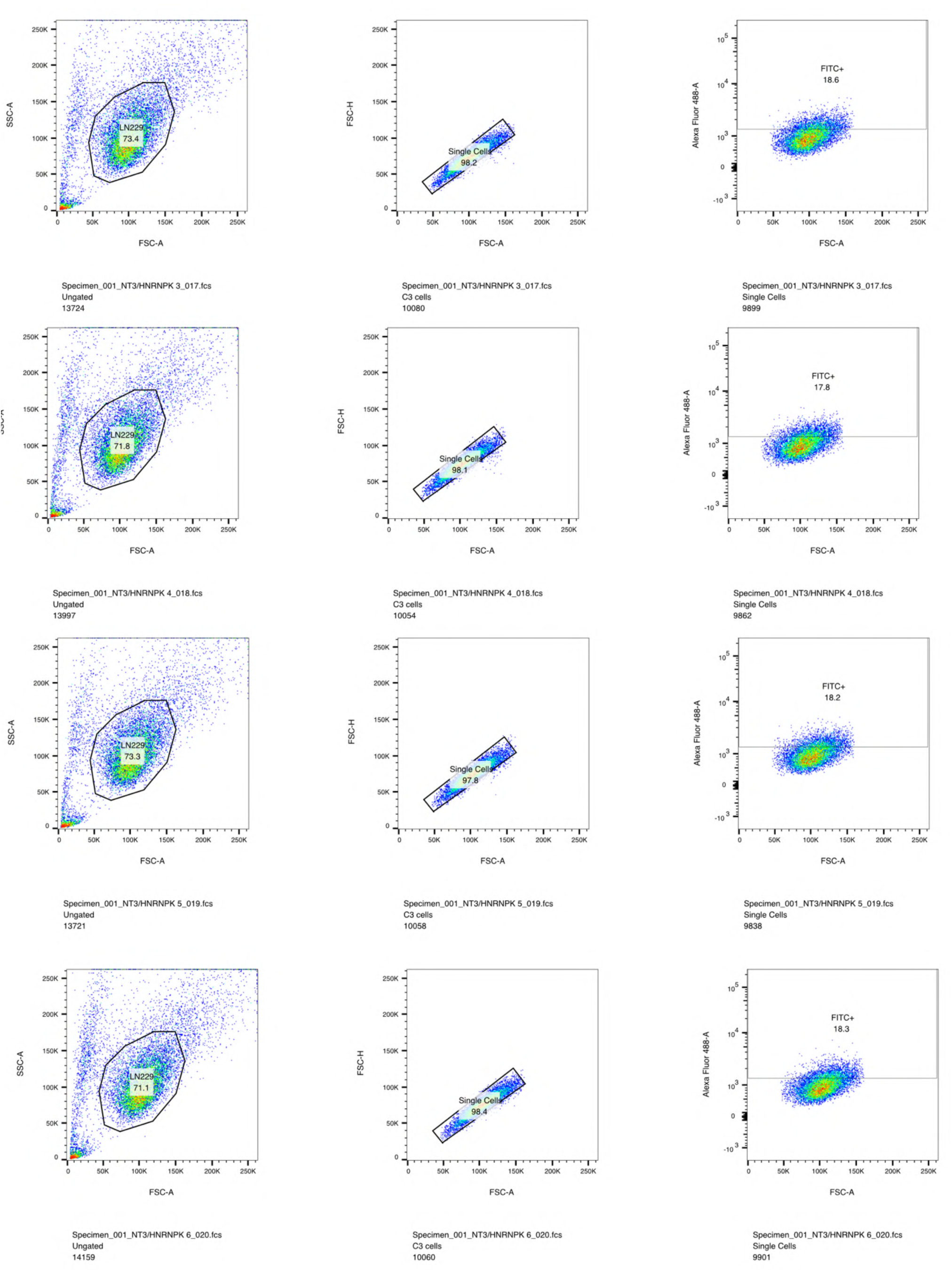
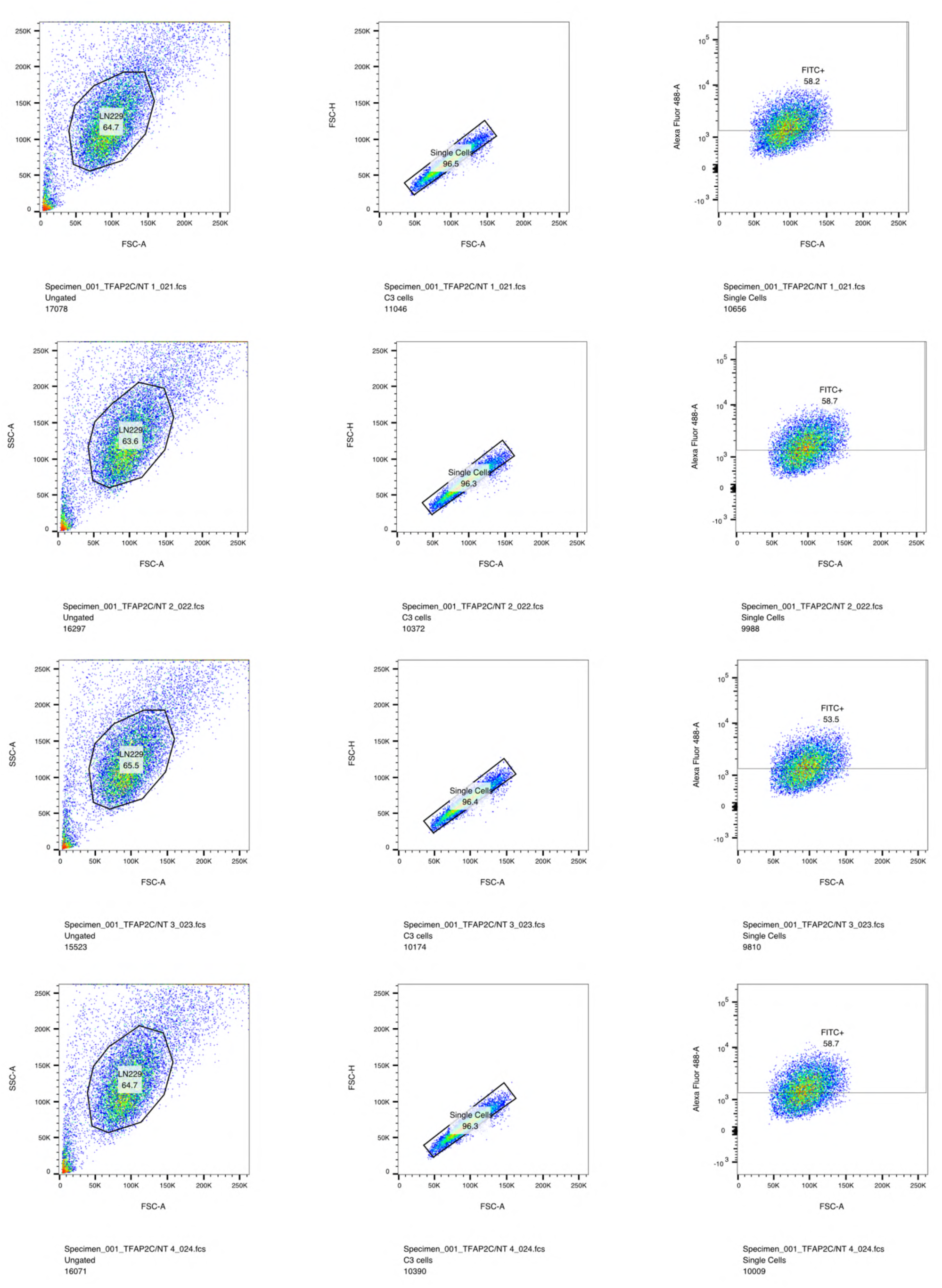
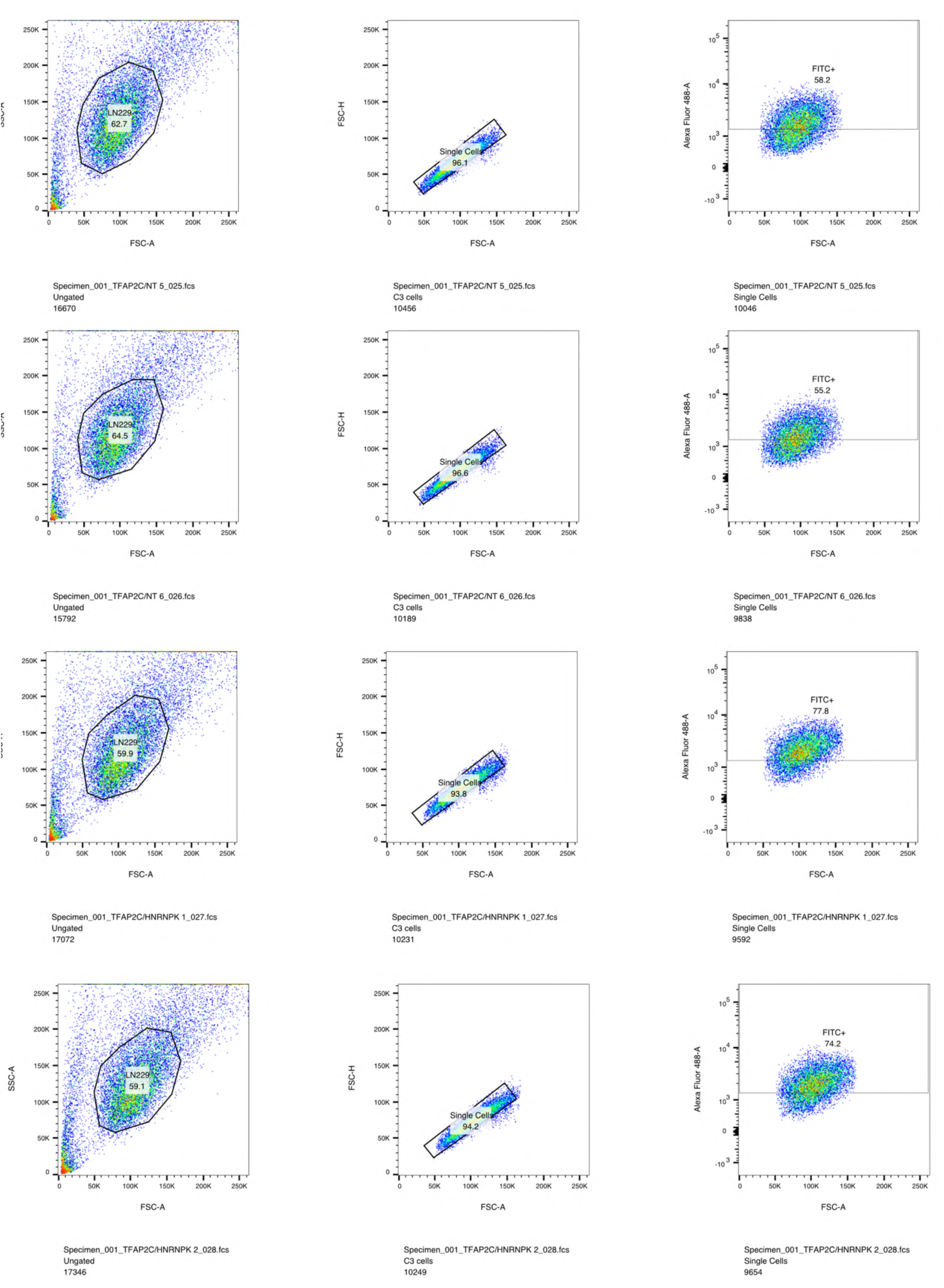
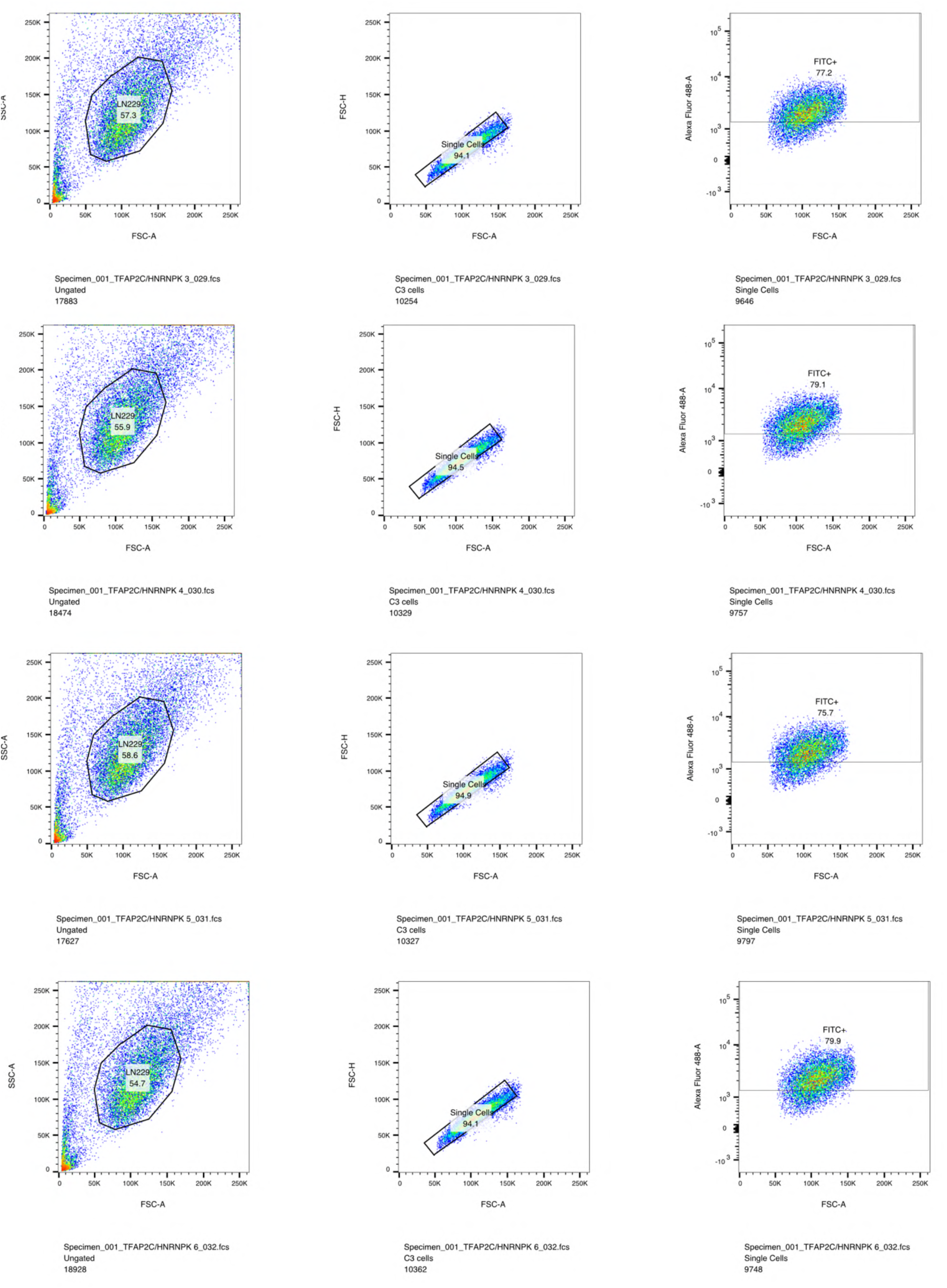
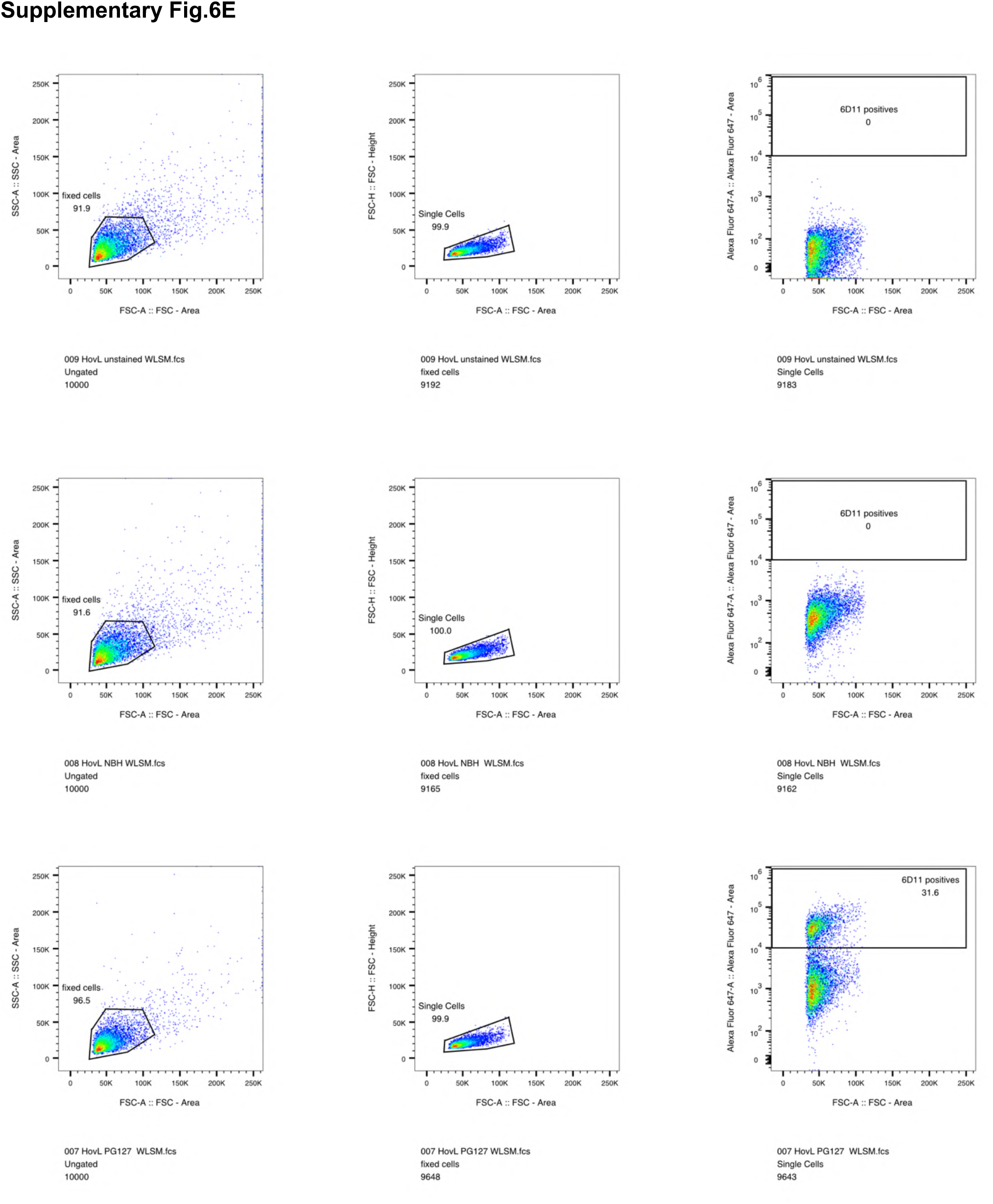

